# In silico codon optimization and phylogenetic analysis of various strains of Sapovirus

**DOI:** 10.64898/2026.10.30.762994

**Authors:** Shinjouli Adhikari, Indra Mani

**Affiliations:** Department of Microbiology, Gargi College, University of Delhi, Siri Fort Road, New Delhi-110049, India

**Keywords:** Sapovirus, CAI, codon optimization, VP1 capsid protein, GC content, DNA, phylogenetic analysis, vaccines

## Abstract

**Introduction:** Sapovirus (SaV) is an emerging enteric pathogen of rising public health importance, responsible for acute gastroenteritis in people of all ages and most often in children under five. Its genome is approximately 7.1-7.7 kb long, positive-sense, single-stranded RNA, comprising two principal open reading frames; processing of the ORF1 product yields seven non-structural proteins and the major capsid protein VP1, while ORF2 encodes the minor structural protein VP2. Given the absence of therapies or an approved vaccine and the low yields of recombinant capsid protein, improving expression remains a priority.

**Methods:** VP1 and VP2 sequences from twelve selected human SaV strains were aligned, phylogenetic trees were reconstructed by maximum likelihood method under the Tamura-Nei model with 1,000 bootstrap replicates, and both genes were optimized *in silico* for *Escherichia coli* using the OPTIMIZER.

**Results:** The trees resolved three genogroups, GI, GII and GV, each strain placed alike by both genes. Every optimized sequence reached the ceiling value of 1.000 for the codon adaptation index (CAI). Mean CAI and GC% increased by 3.07 (207.0%) and 1.11 (11.1%) fold for VP1, and 3.11 (210.9%) and 1.14 (14.1%) fold for VP2, while AT% decreased. All changes were statistically significant (p < 0.001) and the encoded amino acid sequences were unaltered.

**Conclusion:** Our findings indicate that Optmization raised host-matched codon usage in both capsid genes with no truncated proteins produced. The redesigned VP1 and VP2 sequences warrant further investigation as potential candidates for biotherapeutic and vaccine development.

## 1. Introduction

Sapoviruses (SaVs), historically called “Sapporo-like viruses,” carry a non-enveloped icosahedral capsid that encloses a (+) ssRNA genome approximatley 7.1 to 7.7 kb in length (Oka et al., 2016). Distinct calicivirus particles were first observed by electron microscopy in diarrhoeal stools in 1976 (Madeley & Cosgrove, 1976), and in 1977 the virus was linked to an outbreak in an infant home in Sapporo, Japan, from which its name originated (Chiba et al., 1979). The International Committee on Taxonomy of Viruses (ICTV) places the genus within the family *Caliciviridae* as a single recognized species, *Sapovirus sapporoense*, with Sapporo virus as the exemplar isolate (ICTV, 2026; Vinjé et al., 2019). SaV particles measure about 30-40 nm across and possess characteristic cup-like depressions on their capsid surface, producing the distinctive “Star of David” morphology observed by electron microscopy (Madeley, 1979; Saif et al., 1980). Two principal open reading frames occupy the SaV genome, with VPg linked to the 5′ end and polyadenylated at the 3′end .(Hosmillo et al., 2014; Oka et al., 2015).ORF1 encodes a large polyprotein that is subsequently cleaved into seven non-structural proteins (NS1 to NS7), including the RNA-dependent RNA polymerase (RdRp) and the major capsid protein VP1, whereas ORF2 encodes the minor structural protein VP2 (Oka et al., 2015; Li et al., 2017; Tohma et al., 2020). VP1 can also be translated from a subgenomic RNA (Chang et al., 2004; Oka et al., 2009). A third reading frame is also anticipated in certain human and bat SaV genomes, though it has not been fully characterized yet (Oka et al., 2015; Tse et al., 2012).

The complete amino acid sequence of VP1 is used to classify the genus into 19 genogroups, designated GI to GXIX, of which GI, GII, GIV, and GV infect humans, whereas GIII predominates in pigs (Farkas et al., 2004; Oka et al., 2016). Recombination near the RNA-dependent RNA polymerase–VP1 junction further increases diversity and complicates molecular typing (Hansman et al., 2005b).

Acute gastroenteritis (AGE) remains a public health concern, causing more than 2.5 million deaths among children under five (Abdelqader et al., 2025). GI and GII are often reported in AGE and contribute to childhood diarrhoeal disease worldwide (Becker-Dreps et al., 2020). A meta-analysis of sporadic infection put the pooled detection rate at 3.4%, rising to 5.6% in studies that used RT-qPCR and to 4.4% in children under five (Diez Valcarce et al., 2021). In an eight-country birth cohort of 1,715 children, sapovirus was found in 24.7% of diarrhoeal stools, more than 90% of children had been infected by two years of age, and prior infection was associated with a 22% reduction in the risk of reinfection, which indicates that naturally acquired protection is partial (Rouhani et al., 2022). The World Health Organization (WHO) reports that immunity is specific to genotypes, with limited cross-protection even among genogroups (Harada et al., 2012; WHO, 2025). While most infections are mild and self-limiting, with dehydration as the principal complication, severe cases require hospitalization (Bucardo et al., 2014), and deaths have been recorded in outbreaks in long-term care facilities (Lee et al., 2012). No sapovirus-specific mortality estimate exists; the figure of roughly 71,000 under-five deaths attributed to caliciviruses combines norovirus with sapovirus (Lanata et al., 2013), and SaV is one of twelve pathogens modelled in global estimates of under-five diarrhoeal mortality (Black et al., 2024). Where rotavirus vaccination is established, SaV accounts for a larger share of the childhood diarrhoea that remains (Bucardo et al., 2014; Sánchez et al., 2018). Despite this burden, no licensed vaccine and no sapovirus-specific antiviral are available (Becker-Dreps et al., 2019), so control relies on sanitation and hygiene (WHO, 2025).

Transmission occurs via the faecal–oral route, involving direct contact between individuals or exposure to contaminated food, water, and surfaces. Outbreaks have been reported in childcare centres, elderly homes, hospital wards, restaurants, and schools (Becker-Dreps et al., 2020). SaVs have a wide host range, affecting humans alongside many animal species such as bats, swine, minks, dogs, rodents, sea lions, and chimpanzees. (Oka et al., 2016), and viral RNA has been recovered from oysters, which makes shellfish a plausible vehicle for foodborne spread (Ueki et al., 2010), while a foodborne outbreak in Japan followed the consumption of catered box lunches prepared by infected handlers (Kobayashi et al., 2012). The pooled median incubation period is 1.7 days, with a 95% confidence interval of 1.5 to 1.8 days (Lee et al., 2013). Asymptomatic shedding may sustain transmission, with viral shedding continuing for 1–4 weeks after symptom onset (Chiba et al., 1980), a median of 18.5 days in a Peruvian birth cohort (Sánchez et al., 2018), and prolonged excretion in immunocompromised patients (Rubio-Mora et al., 2024). **The prevalence of AGE in children, as reported by WHO, varies from 2% to 19% based on age and setting (WHO, 2025)**. This range includes 8.3% observed in Japanese children (Hoque et al., 2023) and 8% noted in Spain (Oliveira-Tozetto et al., 2021). In India, studies reported SaV in 1.8% of 1,203 children in Kolkata during 2021–2023 and 2.7% of 778 cases in Pune during 2007–2011, documenting genotype diversity and surveillance needs (Lasure & Gopalkrishna, 2017; Saha et al., 2025). Sporadic infection follows a cold-season pattern, peaking between November and March in Japan (Dey et al., 2012) and between November and January in western Canada (Zhuo et al., 2021).

VP1 demonstrates considerable genetic variability, which correlates with antigenic differences among SaV genogroups and genotypes, making it central to molecular classification and surveillance (Hansman et al., 2005a; Oka et al., 2024). Its antigenic diversity also makes VP1 a principal target for antibody studies and recombinant antigen development (Oka et al., 2015; Becker-Dreps et al., 2020). Recombinant VP1 assembles spontaneously into virus-like particles (VLPs) whose morphology and antigenic behaviour resemble those of native SaV virions, supporting their use in serological studies and evaluation of candidate vaccine antigens (Hansman et al., 2005a; Oka et al., 2009). Reverse genetics have confirmed that the virion contains VP2 as a minor structural protein (Li et al., 2022), although its precise functions remain incompletely resolved. Variation in VP2, including deletion patterns, has been reported in porcine SaVs, but its effects on HuSaV antigenicity remain uncertain (Dufkova et al., 2013). VP1 and VP2 are therefore the structural genes of interest for antigen production.

Pathogenesis was difficult to study for decades because HuSaV could not be efficiently propagated in conventional cell culture, until replication was achieved in human duodenal cell lines via bile acid supplementation in 2020 (Takagi et al., 2020), followed by intestinal enteroid (Euller-Nicolas et al., 2023), induced pluripotent stem cell (Matsumoto et al., 2023) and Caco-2 systems (Fukuda et al., 2025). These models demonstrated small-intestinal tropism and bile-acid-dependent replication that, unlike norovirus, is independent of histo-blood group antigens (Euller-Nicolas et al., 2023). Subsequently, in 2026, the scavenger receptor CD36 was identified as a HuSaV receptor with direct binding to viral particles (Haga et al., 2026). Entry also needs conjugated bile acids such as glycocholic acid, which acts after attachment and promotes CD36 internalisation, thereby limiting infection to the small intestine (Takagi et al., 2020; Haga et al., 2026).

Despite the implementation of these systems, recombinant expression continues to be the practical method for producing SaV capsid antigens; however, the yields remain low. Hansman et al. (2006) documented that rVP1 expression in insect cells was lower than that achieved for norovirus, with some strains undetectable, and only a two-to three fold gain was recovered by inserting foreign nucleotides upstream of the VP1 start codon, a modification that did not alter codon usage within the coding sequence itself.

Codon optimization is an established strategy to improve the production of foreign proteins, where the codon composition of a gene is redesigned to suit the translational machinery of another organism, as when a plant gene sequence is modified for expression in human cells or when a human gene sequence is adjusted for expression in bacteria or yeast (Graf et al., 2009). Synonymous codons are not used at equal frequencies, and this codon usage bias varies across species. Consequently, genes carrying codons that are infrequently used by the intended host may be translated less efficiently, while substituting host-favoured alternatives optimizes heterologous expression. (Gustafsson et al., 2004; Basak et al., 2008). Therefore, the present study aimed to optimize the codons of the VP1 and VP2 genes of 12 HuSaV strains spanning genogroups GI, GII, and GV for overexpression in *E. coli* using *in silico* methods and to examine the phylogenetic relationships among the strains analyzed. Only synonymous codons were altered so that the amino acid sequence encoded by each gene remains identical to its native counterpart, with no truncated product expected. This approach is intended to preserve the antigenicity and functional activity of both capsid proteins while enabling the production of recombinant antigens in quantities adequate for immunodiagnostic and immunotherapeutic use.

## 2. Materials and Methods

### 2.1. Collection and analysis of nucleotide sequences

The NCBI GenBank site (https://www.ncbi.nlm.nih.gov/) was used to obtain nucleotide sequences for the VP1 and VP2 capsid protein genes of human Sapovirus. The identity and similarity of the retrieved sequences were assessed against the NCBI nucleotide collection using BLAST (Altschul et al., 1997). Twelve Sapovirus strains were included in the analysis. Tables 1 and 2 provide the accession numbers and genomic coordinates for both genes.

**Table 1.** Human Sapovirus strains analysed in this study and the genomic coordinates of the VP1 coding region.

| **No.** | **Strain** | **Geno-group** | **Geno-type** | **GenBank accession** | **Genome (nt)** | **VP1 start** | **VP1 end** | **VP1 (nt)** | **VP1 (aa)** |
|---|---|---|---|---|---|---|---|---|---|
| 1 | N21 | GI | GI.1 | AY237423.2 | 7429 | 5170 | 6852 | 1686 | 561* |
| 2 | Mc114 | GI | GI.1 | AY237422.3 | 7429 | 5170 | 6852 | 1686 | 561* |
| 3 | NongKhai-50 | GI | GI.1 | AY646853.2 | 7429 | 5170 | 6852 | 1686 | 561* |
| 4 | Chanthaburi-74 | GI | GI.1 | AY646854.2 | 7429 | 5170 | 6852 | 1686 | 561* |
| 5 | ROK62 | GI | GI.1 | KP298674.1 | 7429 | 5170 | 6852 | 1686 | 561* |
| 6 | Dresden | GI | GI.1 | AY694184.2 | 7429 | 5170 | 6852 | 1686 | 561* |
| 7 | Sapporo/MT-2010 | GI | GI.1 | HM002617.1 | 7433 | 5173 | 6855 | 1686 | 561* |
| 8 | Chiba/000496 | GI | GI.4 | AJ606693.2 | 7436 | 5170 | 6852 | 1686 | 561* |
| 9 | Kashiwa1 | GII | GII.5 | LC190463.1 | 7448 | 5174 | 6853 | 1680 | 559 |
| 10 | Mc10 | GII | GII.2 | AY237420.2 | 7458 | 5174 | 6847 | 1677 | 558 |
| 11 | C12 | GII | GII.3 | AY603425.1 | 7476 | 5173 | 6855 | 1686 | 561* |
| 12 | NGY-1 | GV | GV.2 | AB775659.2 | 7521 | 5205 | 6917 | 1716 | 571 |

\* Nucleotide lengths include the terminal stop codon. VP1 comprises 562 codons including the stop codon, giving 561 encoded amino acids. Coordinates were verified individually against each GenBank record.

**Table 2.** Genomic coordinates of the VP2 coding region in the twelve human Sapovirus strains.

| No. | Strain | Geno-group | Geno-type | GenBank accession | Genome (nt) | VP2 start | VP2 end | VP2 (nt) | VP2 (aa) |
|---|---|---|---|---|---|---|---|---|---|
| 1 | N21 | GI | GI.1 | AY237423.2 | 7429 | 6852 | 7349 | 498 | 165 |
| 2 | Mc114 | GI | GI.1 | AY237422.3 | 7429 | 6852 | 7349 | 498 | 165 |
| 3 | NongKhai-50 | GI | GI.1 | AY646853.2 | 7429 | 6852 | 7349 | 498 | 165 |
| 4 | Chanthaburi-74 | GI | GI.1 | AY646854.2 | 7429 | 6852 | 7349 | 498 | 165 |
| 5 | ROK62 | GI | GI.1 | KP298674.1 | 7429 | 6852 | 7349 | 498 | 165 |
| 6 | Dresden | GI | GI.1 | AY694184.2 | 7429 | 6852 | 7349 | 498 | 165 |
| 7 | Sapporo-MT-2010 | GI | GI.1 | HM002617.1 | 7433 | 6855 | 7352 | 498 | 165 |
| 8 | Chiba/000496 | GI | GI.4 | AJ606693.2 | 7436 | 6852 | 7349 | 498 | 165 |
| 9 | Kashiwa1 | GII | GII.5 | LC190463.1 | 7448 | 6853 | 7353 | 501 | 166 |
| 10 | Mc10 | GII | GII.2 | AY237420.2 | 7458 | 6850 | 7350 | 501 | 166 |
| 11 | C12 | GII | GII.3 | AY603425.1 | 7476 | 6858 | 7358 | 501 | 166 |
| 12 | NGY-1 | GV | GV.2 | AB775659.2 | 7521 | 6917 | 7420 | 504 | 167 |

Nucleotide lengths include the terminal stop codon throughout.

### 2.2. Multiple sequence alignment and phylogenetic analysis

The VP1 and VP2 coding sequences were aligned separately using ClustalW. Phylogenetic relationships were then inferred from these alignments by maximum likelihood method, taking the Tamura-Nei substitution model (Tamura and Nei,1993) with 1000 bootstrap replicates. Sites carrying gaps or missing data were handled by partial deletion, so that only positions covered in at least 95% of sequences were retained. All analyses were conducted using MEGA software (version 12) (Kumar et al., 2024).

### 2.3. Codon optimization and analysis

OPTIMIZER (http://genomes.urv.es/OPTIMIZER/), a PHP application designed to predict and raise the gene expression in a heterologous host (Puigbò et al., 2007) was used to optimize and compute the CAI, GC and AT content of the retrieved DNA sequences using *Escherichia coli* str. K-12 substr. MG1655 as the reference host. It is a model organism frequently used for heterologous gene expression. Codon Adaptation Index (CAI) measures the degree of similarity between the codon usage of a gene and that of the reference host (Sharp & Li, 1987) and it was determined for the VP1 and VP2 genes across all 12 strains.

### 2.4. Alignment of native and optimized sequences

Alignments of each native coding sequence against its optimized counterpart were obtained directly from the OPTIMIZER output for both the VP1 and VP2 genes of all twelve strains.

### 2.5. Statistical analysis

Descriptive statistics, including mean, range, and standard deviation, were calculated using GraphPad Prism (version 11). Native and optimized sequence values across various Sapovirus strains were organized into tables and graphically represented to compare their CAI values. For both the VP1 and VP2 genes, variations in CAI, GC, and AT content were evaluated using the Wilcoxon matched-pairs signed rank test, with statistical significance defined by a two-tailed probability of p<0.05.

### 2.6. GenBank Accession Numbers

Codon optimized sequences generated in this study were deposited in NCBI GenBank database under accession numbers QB078484 and QB245777–QB245787 for VP1, and QB245788–QB245799 for VP2.

## 3. Results

### 3.1. Sequence alignment and phylogenetic analysis

The native coding sequences of the VP1 and VP2 genes of all twelve HuSaV strains were aligned separately with ClustalW in MEGA version 12 (Fig. 1A and 1B). The VP1 alignment extended over 1,739 positions, of which 548 (31.5%) carried an identical nucleotide in every strain, while gaps occupied 101 columns (5.8%) spread over 30 distinct regions between positions 15 and 1,311, indicating that indels were distributed throughout the gene rather than confined to one segment. The largest gaps included an 11-nt block at positions 20-30 and a 10-nt block at positions 1,155-1,164. The number of gaps varied among strains, with the GV strain NGY-1 having the fewest at 23 positions, resulting in the longest continuous sequence of 1,716 nucleotides, while the GII strain Mc10 had the most at 62 positions, leading to the shortest sequence of 1,677 nucleotides. Each of the eight GI strains contained 53 gaps.

**Fig. 1.**
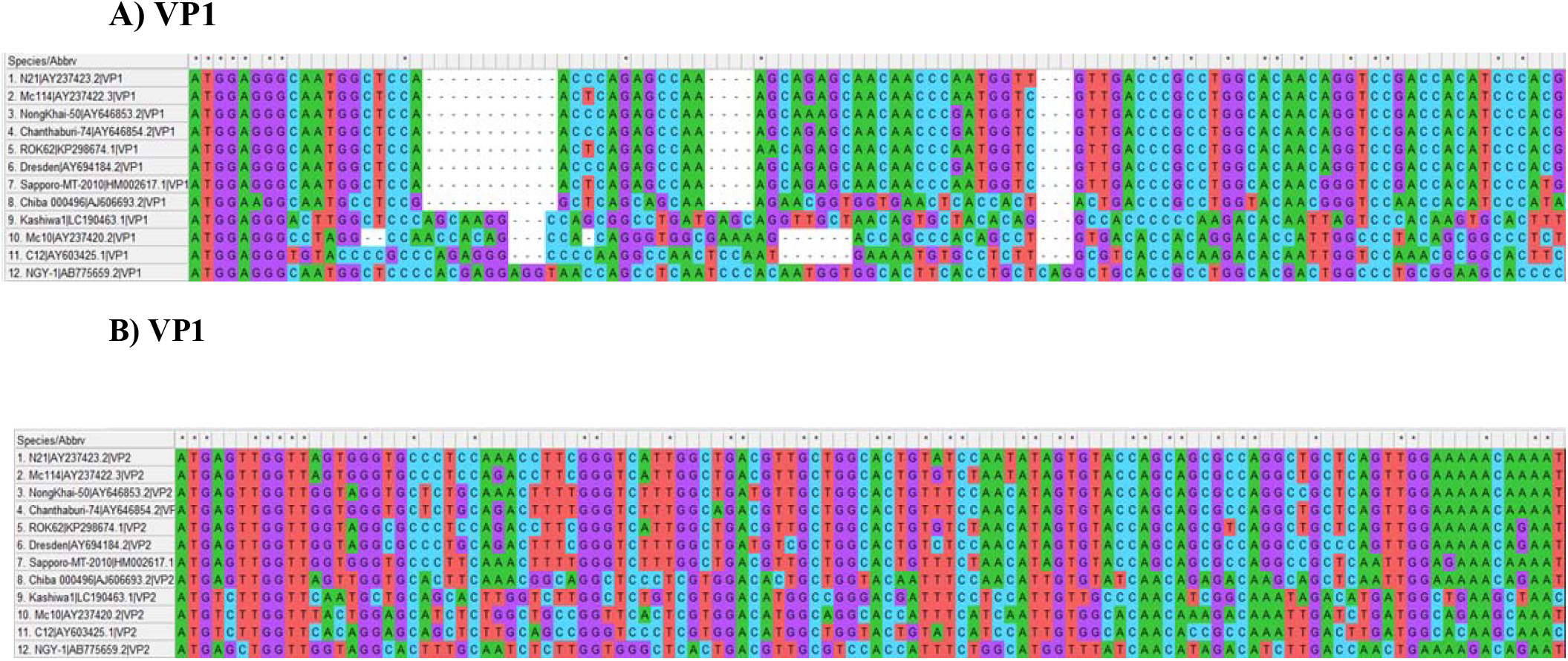
Multiple sequence alignment of the native (A) VP1 and (B) VP2 nucleotide sequences of twelve human Sapovirus strains, using ClustalW in MEGA 12. Only the 5′ terminal regions are shown. Asterisks (*) mark indicates conserved positions; dashes (-) indicate gaps.

In comparison, the VP2 alignment was both shorter and significantly more consistent, covering 509 positions. Among these, 128 positions (25.1%) were identical across all strains, while only 14 positions (2.8%) had a gap. Coding lengths here were categorized into three discrete classes based on genogroup: 498 nt in the eight GI strains, 501 nt in the three GII strains and 504 nt in NGY-1, which alone extended to column 509 of the alignment.

The aligned sequences were then subjected to phylogenetic analysis, in order to construct a phylogenetic tree for both genes using the maximum likelihood (ML) method. The analysis was performed using the Tamura-Nei nucleotide substitution model with a bootstrap value of 1000, as shown in (Fig. 2A for VP1 and 2B for VP2). The starting topology for each search was whichever of a Neighbor-Joining tree from Tamura-Nei pairwise distances and the shortest of ten Maximum Parsimony trees gave the better log likelihood. Partial deletion at a 95% site coverage cutoff removed every column carrying a gap, retaining 1,638 of the 1,739 VP1 positions and 495 of the 509 VP2 positions. The analysis revealed that the twelve strains resolved into three well-defined clades corresponding to genogroups GI, GII, and GV, with each strain’s genogroup classification being concordant between the two genes.

**Fig. 2.**
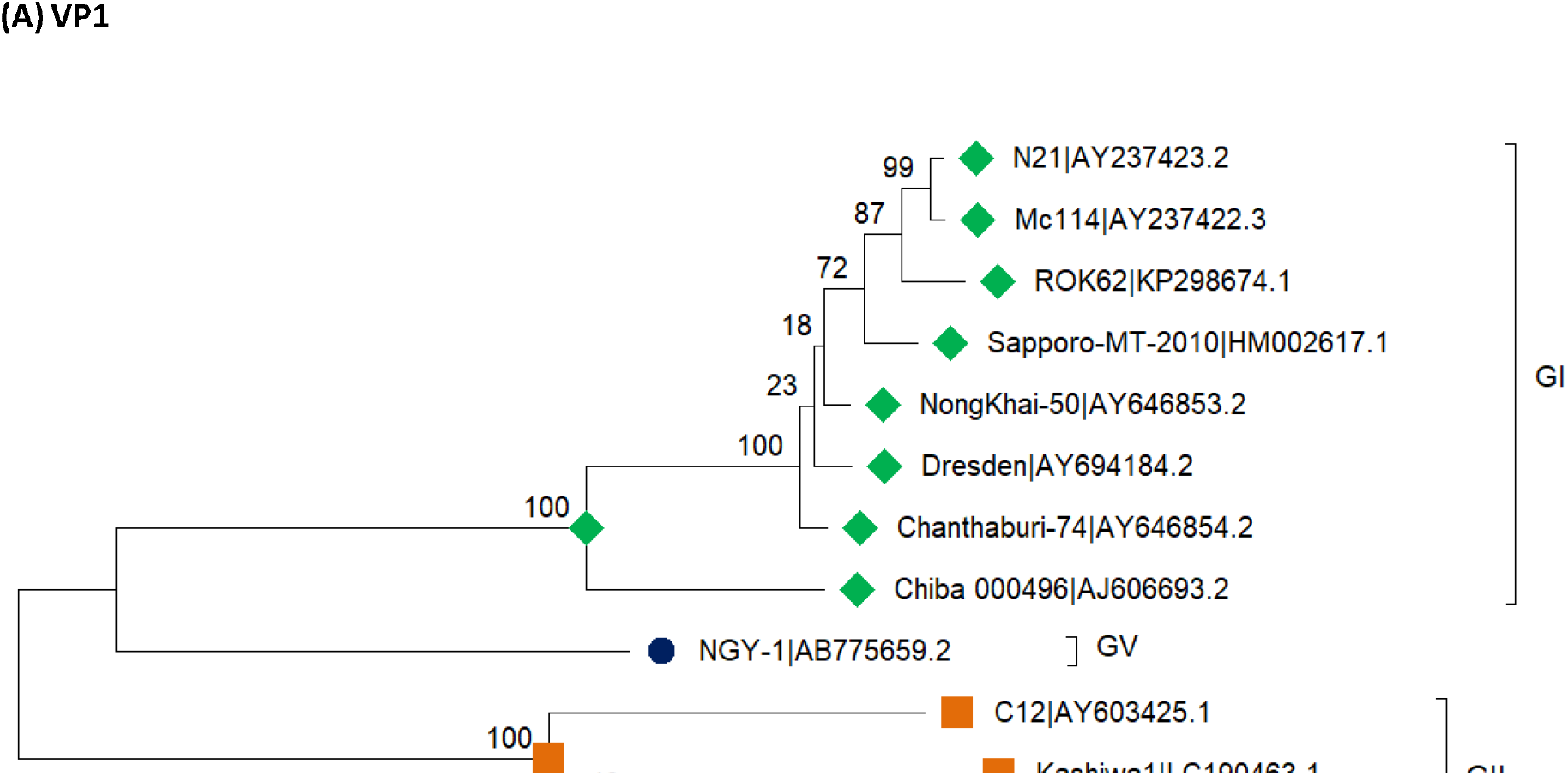
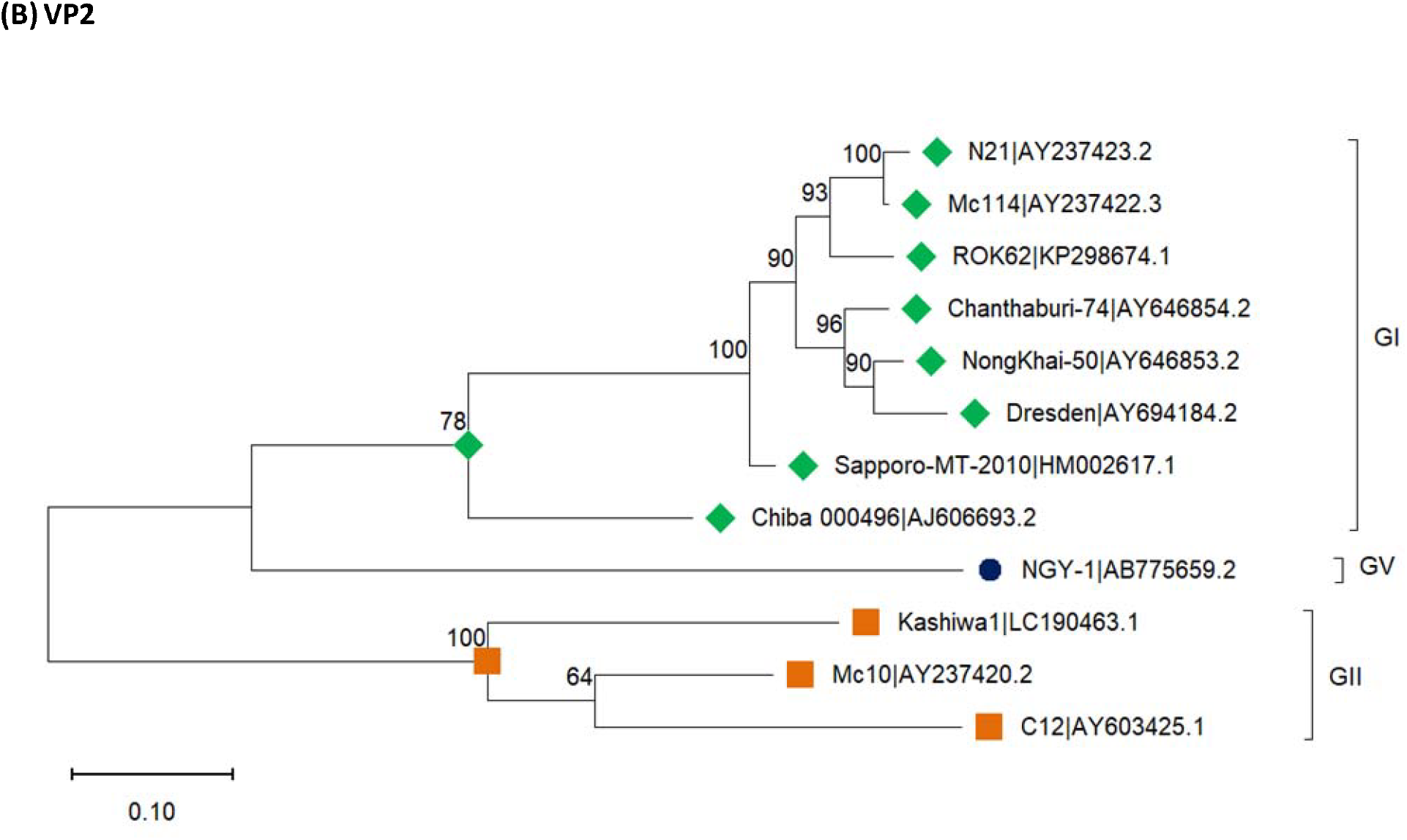
Phylogenetic analysis of the native (A) VP1 and (B) VP2 nucleotide sequences of twelve human Sapovirus strains. Trees were inferred using the Tamura–Nei model with 1,000 bootstrap replicates and 95% site-coverage partial deletion. The final datasets comprised 1,638 VP1 and 495 VP2 positions. Bootstrap support is shown at the nodes. Genogroups: GI (green diamonds), GII (orange squares), GV (blue circle). Scale bar indicates substitutions per site.

For VP1, the final dataset included 1,638 positions, and the tree with the maximum log likelihood of (-11,187.17) is depicted in (Fig. 2A). Within GI, eight strains (N21, Mc114, ROK62, Sapporo-MT-2010, NongKhai-50, Dresden, Chanthaburi-74, and Chiba 000496) were grouped with 100% bootstrap support. Meanwhile, the three GII strains (C12, Kashiwa1, and Mc10) formed a distinct cluster, also supported at 100%. The GV strain, NGY-1, branched separately from both groups. In the GI cluster, N21 and Mc114 were most closely related (99%), with ROK62 (87%) and Sapporo-MT-2010 (72%) joining successively, while the nodes separating NongKhai-50, Dresden, and Chanthaburi-74 had weak support (18% and 23%). Similarly, the internal node of the GII cluster was poorly resolved (40%).

In the case of VP2, the final dataset comprised 495 positions, and the tree with the maximum log likelihood (-3,836.84) is shown in (Fig. 2B). The same three genogroup clusters were recovered, with broader GI grouping supported at 78% and the GII cluster supported at 100%; NGY-1 again forming a separate lineage. Notably, the internal branching within GI was better resolved than in the VP1 tree, where N21 and Mc114 grouped at 100%, ROK62 joined at 93%, Chanthaburi-74 and NongKhai-50 grouped at 96% with Dresden joining at 90%, and Chiba 000496 diverged at the base of the cluster (78%). The internal node of GII reached 64%. Taken together, the two trees illustrate that the strains analysed in this study span three human genogroups, with GII being the most divergent.

### 3.2. Codon optimization

At present, there is neither an authorized vaccine nor a specific antiviral known, and infection management is still primarily supportive (Becker-Dreps et al., 2019). Consequently, this study employed DNA codon optimization to generate sequences capable of yielding sufficient protein in the target host. Both VP1 and VP2 genes across the 12 strains were optimized against *Escherichia coli* K-12 MG1655, as detailed in Tables 3 and 4, respectively. The findings revealed that the CAI of the optimized DNA sequences surpassed that of the corresponding native DNA sequences for both genes, reaching a maximum possible value of 1 in every case. This indicates that each optimized gene uses only the most commonly used synonymous codon of the host at every position.

**Table 3.** Expression levels of the SaV *VP1* gene in *E. coli* for native-type and codon-optimized sequences.

| **Sapovirus strains (GenBank Accession No.)** | **Native type DNA** |  |  | **Optimized DNA** |  |  |
|---|---|---|---|---|---|---|
|  | CAI | GC% | AT% | CAI | GC% | AT% |
| **AY237423.2** | 0.321 | 51.2 | 48.8 | 1 | 56.3 | 43.7 |
| **AY237422.3** | 0.32 | 51.8 | 48.2 | 1 | 56.2 | 43.8 |
| **AY646853.2** | 0.311 | 51.2 | 48.8 | 1 | 56.6 | 43.4 |
| **AY646854.2** | 0.319 | 51.7 | 48.3 | 1 | 56.5 | 43.5 |
| **KP298674.1** | 0.328 | 51.5 | 48.5 | 1 | 56.3 | 43.7 |
| **AY694184.2** | 0.318 | 51.1 | 48.9 | 1 | 56.5 | 43.5 |
| **HM002617.1** | 0.339 | 51.6 | 48.4 | 1 | 56.3 | 43.7 |
| **AJ606693.2** | 0.324 | 51.5 | 48.5 | 1 | 57.4 | 42.6 |
| **LC190463.1** | 0.338 | 52.7 | 47.3 | 1 | 59.2 | 40.8 |
| **AY237420.2** | 0.319 | 51.5 | 48.5 | 1 | 58.9 | 41.1 |
| **AY603425.1** | 0.331 | 52 | 48 | 1 | 58.5 | 41.5 |
| **AB775659.2** | 0.341 | 51 | 49 | 1 | 59 | 41 |
| **N** | 12 | 12 | 12 | 12 | 12 | 12 |
| **Min.** | 0.311 | 51 | 47.3 | 1 | 56.2 | 40.8 |
| **Max.** | 0.341 | 52.7 | 49 | 1 | 59.2 | 43.8 |
| **Mean±SD** | 0.326 ± 0.010 | 51.57 ± 0.464 | 48.43 ± 0.464 | **1.00 ± 0.00**\*\* | **57.31 ± 1.224**\*\* | **42.69 ± 1.224**\*\* |

\*\* p < 0.001 in comparison with the native-type gene sequences (Wilcoxon matched-pairs signed rank test, exact two-tailed P = 0.0005, n = 12 pairs, no ties).

**Table 4.** Expression levels of the SaV VP2 gene in E. coli for native-type and codon-optimized sequences.

| **Sapovirus strains (GenBank Accession No.)** | **Native type DNA** |  |  | **Optimized DNA** |  |  |
|---|---|---|---|---|---|---|
|  | CAI | GC% | AT% | CAI | GC% | AT% |
| **AY237423.2** | 0.308 | 49.4 | 50.6 | 1 | 56.6 | 43.4 |
| **AY237422.3** | 0.317 | 49.8 | 50.2 | 1 | 56.6 | 43.4 |
| **AY646853.2** | 0.332 | 49.2 | 50.8 | 1 | 57 | 43 |
| **AY646854.2** | 0.313 | 48.4 | 51.6 | 1 | 57.2 | 42.8 |
| **KP298674.1** | 0.333 | 49 | 51 | 1 | 56.4 | 43.6 |
| **AY694184.2** | 0.318 | 51.6 | 48.4 | 1 | 57.8 | 42.2 |
| **HM002617.1** | 0.324 | 50.8 | 49.2 | 1 | 57.2 | 42.8 |
| **AJ606693.2** | 0.394 | 50.2 | 49.8 | 1 | 57.6 | 42.4 |
| **LC190463.1** | 0.308 | 53.5 | 46.5 | 1 | 57.5 | 42.5 |
| **AY237420.2** | 0.309 | 53.9 | 46.1 | 1 | 60.9 | 39.1 |
| **AY603425.1** | 0.298 | 54.1 | 45.9 | 1 | 60.1 | 39.9 |
| **AB775659.2** | 0.306 | 48.6 | 51.4 | 1 | 59.5 | 40.5 |
| **N** | 12 | 12 | 12 | 12 | 12 | 12 |
| **Min.** | 0.298 | 48.4 | 45.9 | 1 | 56.4 | 39.1 |
| **Max.** | 0.394 | 54.1 | 51.6 | 1 | 60.9 | 43.6 |
| **Mean±SD** | 0.322 ± 0.025 | 50.71 ± 2.090 | 49.29 ± 2.090 | **1.00 ± 0.00\*\*** | **57.87 ± 1.479\*\*** | **42.13 ± 1.479\*\*** |

\*\* p < 0.001 in comparison with the native-type gene sequences (Wilcoxon matched-pairs signed rank test, exact two-tailed P = 0.0005, n = 12 pairs, no ties).

For the native VP1 gene across all twelve strains, the CAI values, along with the GC and AT content, spanned ranges of 0.311 to 0.341, 51.0 to 52.7, and 47.3 to 49.0, respectively, with corresponding mean values (± standard deviation) of 0.326 (± 0.010), 51.57 (± 0.464), and 48.43 (± 0.464). After optimization, the GC and AT proportions in the modified DNA sequences shifted to ranges of 56.2 to 59.2 and 40.8 to 43.8, with mean values (± standard deviation) of 57.31 (± 1.224) and 42.69 (± 1.224), respectively. Statistical comparison revealed that the mean CAI, GC, and AT indices of the optimized VP1 sequences differed significantly from their native counterparts (p < 0.001). Specifically, the mean CAI and GC% increased by 3.07 (207.0%) fold and 1.11 (11.1%) fold relative to native baselines, whereas the mean AT content declined by 11.9% (Table 3). Variations in CAI values across the evaluated strains are illustrated graphically in Figure 3, where CAI values are plotted against the strain distribution. In addition, the alignment of the native and codon-optimized VP1 sequences is provided in Supplementary Figure S1.

**Fig. 3.**
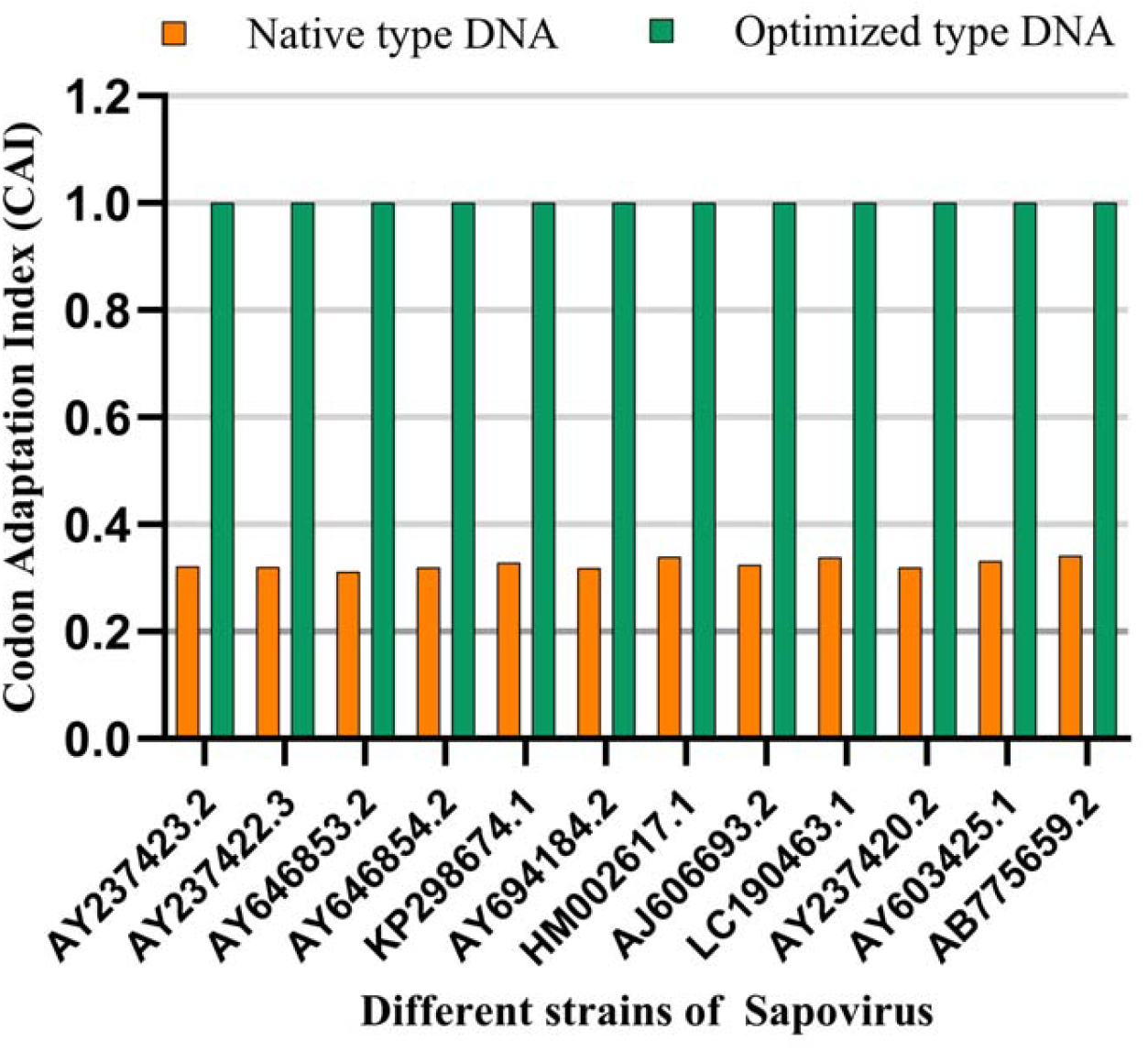
Comparison of the native and codon optimized DNA sequences of VP1 gene across various strains of human Sapovirus.

Likewise, for the native VP2 gene across all twelve strains, the CAI values along with the GC and AT content, spanned ranges of 0.298 to 0.394, 48.4 to 54.1 and 45.9 to 51.6 respectively, with corresponding mean values (± standard deviation) of 0.322 (± 0.025), 50.71 (± 2.090), and 49.29 (± 2.090). Following optimization, the CAI value reached a uniform maximum score of 1.00 across all twelve strains, while the GC and AT proportions in the optmized DNA sequences shifted to ranges of 56.4–60.9 and 39.1–43.6, with mean values (±SD) of 57.87 (± 1.479) and 42.13 (± 1.479) respectively. Statistical evaluation demonstrated that the mean CAI, GC, and AT indices of the optimized VP2 sequences significantly differed from their native counterparts (p < 0.001). in optimized DNA, the mean CAI and GC**%** were found to be 3.11 (210.9%) and 1.14 (14.1%) fold greater than the relative native baselines, whereas the mean AT content decreased by 14.5% (Table 4). Variations in CAI values across the evaluated strains are graphically represented in Figure 4, plotting CAI values against the strain distribution. Furthermore, the alignment between the native and codon-optimized VP2 sequences is provided in Supplementary Figure 2.

**Fig. 4.**
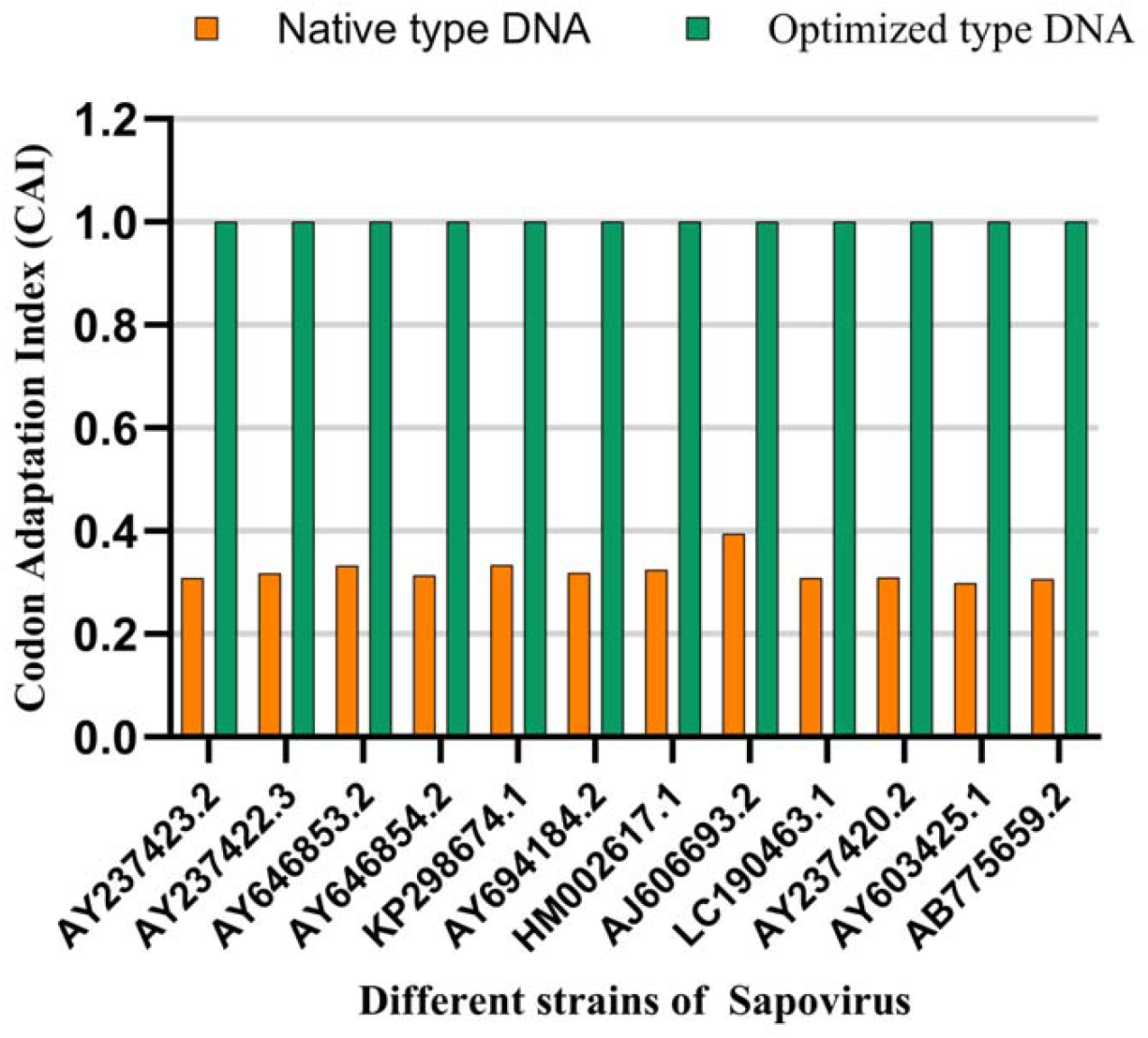
Comparison of the native and codon optimized DNA sequences of VP2 gene across various strains of human Sapovirus.

Interestingly, the baseline variability of the native sequences differed notably between the two genes. Across the twelve strains, native VP2 varied more widely than native VP1 in terms of CAI (SD 0.025 against 0.010) and GC content (SD 2.090 against 0.464). All sequences attained the maximum CAI value of 1, which is the upper limit established by the optimization method. Consequently, the magnitude of the gain was determined by the native value. Furthermore, this extensive codon optimization fully preserved the amino acid sequences encoded by the VP1 and VP2 capsid protein genes without any alterations.

## 4. Discussion

Codon optimization leverages the degeneracy of the genetic code, allowing a single protein to be encoded by multiple alternative nucleotide sequences. These sequences produce the same protein but can differ in production levels. Since many native sequences express poorly in heterologous systems, enhancing their expression can reduce production costs and facilitate the study of low-abundance proteins (Wong et al., 2025). Codon usage is neither random nor consistent across species; for instance, in Escherichia coli and yeast, the codons preferred by genes with high expression levels correspond to the more abundant tRNAs.(Ikemura, 1985), whereas genes expressed at low levels select among synonymous codons in a much more random fashion (Gouy & Gautier, 1982), Thus, synonymous codon choice is a conserved mechanism of expression control that does not change the encoded protein, although the mechanisms underlying codon optimality and its effects remain to be resolved (Barrington et al., 2023; Wong et al., 2025). Given the observed correlation between bias and yield, it is suggested that gene expression can be enhanced by aligning a gene with the host’s codon usage pattern and, the codon adaptation index was created to measure the degree of this alignment (Sharp & Li, 1987). Numerous optimization programs have been developed based on this principle (Puigbo et al., 2007), most of which, for *E. coli*, take the codon usage of the expression host as their reference (Wong et al., 2025). This redesign facilitates reducing tandem repeats, adding or removing restriction sites, and adjusting translational rate. It is now recommended as an initial step in designing high-expression constructs for gene therapy and nucleic acid vaccines (Mauro & Chappell, 2014).

Additionally, research involving various hosts has shown substantial gains. For example, the optimization of a multistage malaria vaccine candidate led to at least a threefold increase in expression within E. coli, the host used in this study (Zhou et al., 2004). A comparison of two strategies in that organism revealed that the selected algorithm determines yield (Menzella, 2011). In mammalian cells, recoding resulted in a six-to ninefold boost in output (Slimko & Lester, 2003). From another perspective, RNA virus genes, hindered at various stages rather than a single one, were expressed with at least fifty times less efficiency from unmodified plasmids compared to optimized ones (Ternette et al., 2007). This advantage is also seen in DNA vaccines, where a codon-optimized HIV-1 gag construct increased gene expression by more than tenfold over the native counterpart (Deml et al., 2001). Beyond protein therapeutics, the same approach has raised yields of synthetic biofuels, enzymes, and vaccines for human use across diverse expression systems (Demissie et al., 2025). For instance, codon optimization of an HIV-1 reverse transcriptase DNA vaccine enhanced antigen expression and increased antibody and cellular immune responses in mice (Latanova et al., 2018). Furthermore, codon usage analysis is not limited to expression purposes; patterns of codon choice have been read to infer which hosts the RNA viruses recovered from acidic hot-spring metagenomes infect, whether those hosts are bacterial, archaeal or eukaryotic (Stedman et al., 2013).

The phylogenetic analysis carried out here placed the twelve strains into three clades corresponding to genogroups GI, GII, and GV, with each strain assigned identically by VP1 and VP2. Concordance between the two separately analyzed datasets supports the VP1-based genogroup classification (Oka et al., 2016), and the VP2 coding lengths further support this, as they fall into three categories that match the same genogroups. A dual typing system, utilizing complete VP1 and RdRp sequences, has since been proposed to enhance Sapovirus genotype classification by incorporating genetic-distance criteria (Zhao et al., 2024). Nonetheless, the internal relationship resolution varied between the two genes, with VP2 providing stronger support for internal GI nodes compared to VP1. Two internal VP1 nodes showed less than 25% bootstrap support, whereas the corresponding VP2 nodes ranged from 90% to 100%, indicating the short internal branches that separate closely related GI genotypes and the uneven evolutionary rate distribution across genome regions and genotypes (Tohma et al., 2020). The observed VP2 coding lengths also differed among the selected genogroups. Although GII was the most divergent group in both trees, additional pairwise genetic distance analysis is necessary to confirm its relative divergence. Comparable genogroup-level divergence, accompanied by variations in capsid region length, has likewise been reported in porcine SaV strains (Liu et al., 2022). Molecular surveillance of strains circulating in young children continues to reveal such diversity in both sporadic infections (Ji et al., 2024) and in outbreaks (Su et al., 2023).

Following this, the study determined the average CAI, GC, and AT compositions and compared them with the values of their respective codon-optimized sequences across all twelve strains. For both the VP1 and VP2 genes, the optimized DNA sequences demonstrated significantly different and elevated values compared to the native-type sequences. Optimization enhanced the mean CAI of the VP1 gene sequences by 3.07 times (an increase of 207.0%), whereas the VP2 gene showed a 3.11-fold increase, which is a 210.9% rise. Additionally, an increase in the GC% was observed in the optimized DNA sequences compared to the native sequences. Specifically, GC proportions were enhanced by 1.11 times (11.1% increase) in the VP1 gene and 1.14 times (14.1% increase) in the VP2 gene for the optimized DNA. Conversely, a systematic decline in AT content was observed compared to the native values. These values reflect the adaptation of each gene to its current host of choice, but adaptation alone is insufficient to ensure improved translational efficiency under selection pressure (Rahman et al., 2018; Feng et al., 2022). A comparable *in silico* approach has been applied to Ebola virus (Mathuria et al., 2024), Zika virus (Hora et al., 2023), Nipah virus (Gupta et al., 2021), SARS-CoV-2 (Al Zamane et al., 2021), and influenza A virus (Mani et al., 2011).

VP1 exhibits high genetic variability and is closely associated with immunogenicity (Oka et al., 2024), though its P2 subdomain is highly constrained against amino acid changes (Yokoyama et al., 2024) and is not affected by synonymous recoding. Variation has also been reported in VP2, and deletion patterns may influence its antigenic epitopes (Dufkova et al., 2013; Liu et al., 2022). Both, therefore, serve as suitable vaccine candidates. There is precedent with the porcine sapovirus where an RNA particle vaccine encoding the VP1 gene raised virus-neutralising (VN) antibody titres in colostrum and milk (Aryal et al., 2026) with no equivalent currently existing for the HuSaV. Furthermore, codon optimization can enhance the expression of these genes in cells, and when combined with immunological studies to identify effective antibodies, it is expected to produce vaccines scalable to industrial volumes. Notably, codon optimization has been shown to improve the stability and immunogenicity of mRNA vaccines (Zhang et al., 2023), and both Moderna and Pfizer/BioNTech COVID-19 vaccines utilize codon-optimized coding sequences (Paremskaia et al., 2024). These examples support advancing the optimized VP1 and VP2 sequences, although their efficient expression and protective efficacy still require experimental validation. On the other hand, profiling preferred and under-represented codons and codon pairs can guide the design of synthetic attenuated viruses for live-attenuated vaccines (Lauring et al., 2010), since genomes rebuilt from rare pairs replicate less efficiently and express less protein without provoking a strong host response (Gaunt & Digard, 2022). Thus, identifying under-represented codon pairs in Sapovirus genomes could aid in guiding future deoptimization strategies for vaccine development, offering a potential approach to help prevent future SaV outbreaks.

Although codon optimization is widely used in recombinant protein therapeutics, nucleic acid treatments, and in ongoing clinical trials globally (Paremskaia et al., 2024), it is important to acknowledge its potential demerits. There is evidence that synonymous codon modifications introduced during codon optimization can have unintended effects, such as disturbing normal tRNA usage and altering protein stability, conformation and function, inadvertently eliciting host immune responses. They may even shift post-transcriptional nucleotide modification sites and synthesize an altered repertoire of proteins (Paremskaia et al., 2024). Additionally, synonymous mutations have been associated with several diseases (Mauro & Chappell, 2014). A study demonstrated these effects by showing that synonymous codon changes altered a protein’s fluorescent properties due to changes in protein folding (Sander et al., 2014). Over-optimization carries its own risk, as both modelling and expression tests in *E. coli* indicate that pushing optimal codon use too highcan lower protein yield while raising cellular burden (Roots et al., 2025). Therefore, codon optimization may reduce the efficacy of the biotech therapeutics it aims to improve and should be reconsidered before *in vivo* use.

## 5. Conclusion

HuSaV is a leading cause of acute gastroenteritis (AGE) across all age groups, with the highest incidence observed in children under five and is clinically indistinguishable from norovirus. Onset is abrupt, featuring vomiting or watery diarrhea, nausea, and abdominal cramps, and coinfection with other enteric pathogens is frequent. The burden has long been overlooked for want of a commercial diagnostic test, with detection rates increasing only after molecular assays became available in clinical laboratories. Therefore, it is necessary to recognize this emerging problem. HuSaVs are highly diverse, and the host factors necessary for their infection and spread are only partly defined. The global demand for effective vaccines and treatment regimes persists due to the absence of immunological therapies and preventive measures, a search that still continues. Vaccine and diagnostic antigens are proteins, and recombinant capsid proteins are seldom obtained in sufficient quantities because viral codon usage differs from that of the expression host. Thus, the present study highlights this concern and optimizes the codons of the VP1 and VP2 genes of twelve HuSaV strains to enhance their expression in *Escherichia coli*. Phylogenetic analysis placed these strains in three genogroups, GI, GII and GV, with both genes assigning each strain identically.The CAI and GC content for both genes increased significantly beyond their native values (p < 0.001), while the amino acid sequences remained unchanged. The optimized DNA can therefore be chemically synthesized and overexpressed in *E. coli*, with the elevated CAI expected to yield higher outputs than native DNA. Considering this study and prior work, we believe that VP1 and VP2 are strong candidates for SaV vaccine formulation. Moving forward, laboratory in vitro assays are essential to validate these computational findings, verify expression yields, and evaluate the safety and immunogenic potency of the expressed antigens. This groundwork can subsequently be scaled up for industrial application. These findings add to the genomic characterization of SaV and provide a basis for future work on vaccine development and viral control.

## Acknowledgments

Authors express their gratitude to the Principal of Gargi College for providing the necessary infrastructural support.

## Supplementary Information

Supplementary figure S1 and S2 provided separately.

