## Supplement Figure S1 for "In silico codon optimization and phylogenetic analysis of various strains of Sapovirus"

### Supplementary Figures

**Supplementary Figure S1.** Sequence alignment of the native-type (Query) and codon-optimized VP1 gene sequences of twelve human Sapovirus strains (GenBank accession numbers AY237423.2, AY237422.3, AY646853.2, AY646854.2, KP298674.1, AY694184.2, HM002617.1, AJ606693.2, LC190463.1, AY237420.2, AY603425.1 and AB775659.2).

**The specific-coloured symbols stand for (***): Transversion change, (#): Transition change and (|): Unchanged nucleotide.*

**N21 (AY237423.2)** – 1,686 nucleotides; 1,294 unchanged, 207 transition and 185 transversion changes.

```
           M   E   G   N   G   S   N   P   E   P   K   Q   S   N   N   P   M   V   V   D
    Query  ATG GAG GGC AAT GGC TCC AAC CCA GAG CCA AAG CAG AGC AAC AAC CCA ATG GTT GTT GAC
           ||| ||# ||# ||# ||# ||# ||| ||# ||# ||# ||# ||| **# ||| ||| ||# ||| ||| ||| |||
Optimized  ATG GAA GGT AAC GGT TCT AAC CCG GAA CCG AAA CAG TCT AAC AAC CCG ATG GTT GTT GAC

           P   P   G   T   T   G   P   T   T   S   H   V   V   V   A   N   P   E   Q   P
    Query  CCG CCT GGC ACA ACA GGT CCG ACC ACA TCC CAC GTT GTT GTT GCT AAT CCG GAG CAA CCC
           ||| ||* ||# ||* ||* ||| ||| ||| ||* ||# ||| ||| ||| ||| ||* ||# ||| ||# ||# ||*
Optimized  CCG CCG GGT ACC ACC GGT CCG ACC ACC TCT CAC GTT GTT GTT GCG AAC CCG GAA CAG CCG

           N   G   A   A   Q   R   L   E   L   A   V   A   T   G   A   I   Q   S   N   V
    Query  AAT GGG GCC GCA CAG CGC CTG GAG TTG GCT GTT GCC ACT GGT GCA ATC CAA TCC AAT GTC
           ||# ||* ||* ||# ||| ||# ||| ||# #|| ||* ||| ||* ||# ||| ||# ||| ||# ||# ||# ||#
Optimized  AAC GGT GCG GCG CAG CGT CTG GAA CTG GCG GTT GCG ACC GGT GCG ATC CAG TCT AAC GTT

           P   E   A   I   R   N   C   F   A   V   F   R   T   F   A   W   N   D   R   M
    Query  CCT GAG GCA ATA CGC AAC TGT TTT GCA GTC TTT CGT ACT TTT GCT TGG AAC GAC AGG ATG
           ||* ||# ||# ||* ||# ||| ||# ||# ||# ||# ||# ||| ||# ||# ||* ||| ||| ||| *|* |||
Optimized  CCG GAA GCG ATC CGT AAC TGC TTC GCG GTT TTC CGT ACC TTC GCG TGG AAC GAC CGT ATG

           P   T   G   T   F   L   G   S   I   S   L   H   P   N   I   N   P   Y   T   S
    Query  CCC ACG GGA ACT TTT CTT GGA TCT ATA TCG CTT CAT CCC AAC ATT AAC CCG TAC ACT TCT
           ||* ||* ||* ||# ||# ||* ||* ||| ||* ||* ||* ||# ||* ||| ||# ||| ||| ||| ||# |||
Optimized  CCG ACC GGT ACC TTC CTG GGT TCT ATC TCT CTG CAC CCG AAC ATC AAC CCG TAC ACC TCT

           H   L   S   G   M   W   A   G   W   G   G   S   F   E   V   R   L   S   I   S
    Query  CAC CTC TCT GGG ATG TGG GCC GGG TGG GGC GGT AGT TTT GAG GTC CGG CTA TCG ATC TCT
           ||| ||* ||| ||* ||| ||| ||* ||* ||| ||# ||| **| ||# ||# ||# ||* ||# ||* ||| |||
Optimized  CAC CTG TCT GGT ATG TGG GCG GGT TGG GGT GGT TCT TTC GAA GTT CGT CTG TCT ATC TCT

           G   S   G   V   F   A   G   R   I   I   A   S   V   I   P   P   G   V   D   P
    Query  GGT TCT GGC GTG TTC GCT GGG CGC ATC ATT GCT TCT GTC ATA CCA CCA GGG GTT GAT CCC
           ||| ||| ||# ||* ||| ||* ||* ||# ||| ||# ||* ||| ||# ||* ||# ||# ||* ||| ||# ||*
Optimized  GGT TCT GGT GTT TTC GCG GGT CGT ATC ATC GCG TCT GTT ATC CCG CCG GGT GTT GAC CCG
```

```
           S   S   I   R   D   P   G   V   L   P   H   A   F   V   D   A   R   I   T   E
Query      TCG TCC ATC AGG GAC CCA GGC GTG TTG CCT CAC GCT TTC GTT GAT GCT CGC ATC ACT GAG
           ||* ||# ||| *|* ||| ||# ||# ||* #|| ||* ||| ||* ||| ||| ||# ||* ||# ||| ||# ||#
Optimized  TCT TCT ATC CGT GAC CCG GGT GTT CTG CCG CAC GCG TTC GTT GAC GCG CGT ATC ACC GAA

           P   V   S   F   M   I   P   D   V   R   A   V   D   Y   H   R   M   D   G   A
Query      CCA GTT TCT TTC ATG ATC CCT GAT GTA AGG GCA GTG GAC TAC CAT AGG ATG GAT GGT GCT
           ||# ||| ||| ||| ||| ||| ||* ||# ||* *|* ||# ||* ||| ||| ||# *|* ||| ||# ||| ||*
Optimized  CCG GTT TCT TTC ATG ATC CCG GAC GTT CGT GCG GTT GAC TAC CAC CGT ATG GAC GGT GCG

           E   P   T   C   S   L   G   F   W   V   Y   Q   P   L   L   N   P   F   S   T
Query      GAA CCC ACA TGC TCC TTG GGG TTT TGG GTT TAT CAA CCA TTA CTC AAC CCC TTC TCC ACC
           ||| ||* ||* ||| ||# #|| ||* ||# ||| ||| ||# ||# ||# #|# ||* ||| ||* ||| ||# |||
Optimized  GAA CCG ACC TGC TCT CTG GGT TTC TGG GTT TAC CAG CCG CTG CTG AAC CCG TTC TCT ACC

           T   A   V   S   T   C   W   V   S   V   E   T   K   P   G   G   D   F   D   F
Query      ACC GCC GTT TCA ACA TGT TGG GTG TCA GTG GAA ACT AAA CCA GGT GGT GAT TTT GAC TTC
           ||| ||* ||| ||* ||* ||# ||| ||* ||* ||* ||| ||# ||| ||# ||| ||| ||# ||# ||| |||
Optimized  ACC GCG GTT TCT ACC TGC TGG GTT TCT GTT GAA ACC AAA CCG GGT GGT GAC TTC GAC TTC

           C   L   L   R   P   P   G   Q   Q   M   E   N   G   V   S   P   E   G   L   L
Query      TGT CTG CTA AGG CCA CCA GGT CAA CAG ATG GAA AAT GGG GTT TCA CCG GAA GGT CTA TTG
           ||# ||| ||# *|* ||# ||# ||| ||# ||| ||| ||| ||# ||* ||| ||* ||| ||| ||| ||# #||
Optimized  TGC CTG CTG CGT CCG CCG GGT CAG CAG ATG GAA AAC GGT GTT TCT CCG GAA GGT CTG CTG

           P   R   R   L   G   Y   S   R   G   N   R   V   G   G   L   V   V   G   M   I
Query      CCT AGG CGC TTG GGT TAC TCG CGT GGT AAC CGT GTT GGT GGC TTG GTG GTT GGC ATG ATT
           ||* *|* ||# #|| ||| ||| ||* ||| ||| ||| ||| ||| ||| ||# #|| ||* ||| ||# ||| ||#
Optimized  CCG CGT CGT CTG GGT TAC TCT CGT GGT AAC CGT GTT GGT GGT CTG GTT GTT GGT ATG ATC

           L   V   A   E   H   K   Q   V   N   R   H   F   N   S   N   S   V   T   F   G
Query      TTG GTA GCT GAG CAT AAG CAG GTC AAT AGA CAC TTT AAT TCC AAT TCA GTC ACC TTT GGT
           #|| ||* ||* ||# ||# ||# ||| ||# ||# *|* ||| ||# ||# ||# ||# ||* ||# ||| ||# |||
Optimized  CTG GTT GCG GAA CAC AAA CAG GTT AAC CGT CAC TTC AAC TCT AAC TCT GTT ACC TTC GGT

           W   S   T   A   P   V   N   P   M   A   A   E   I   V   T   N   Q   A   H   S
Query      TGG TCA ACG GCA CCT GTT AAT CCA ATG GCA GCA GAG ATT GTA ACT AAC CAG GCA CAT TCA
           ||| ||* ||* ||# ||* ||| ||# ||# ||| ||# ||# ||# ||# ||* ||# ||| ||| ||# ||# ||*
Optimized  TGG TCT ACC GCG CCG GTT AAC CCG ATG GCG GCG GAA ATC GTT ACC AAC CAG GCG CAC TCT

           T   S   R   H   A   W   L   S   I   G   A   Q   N   K   G   P   L   F   P   G
Query      ACA TCA CGC CAT GCT TGG CTT TCC ATT GGT GCT CAG AAT AAG GGT CCA TTG TTT CCC GGG
           ||* ||* ||# ||# ||* ||| ||* ||# ||# ||| ||* ||| ||# ||# ||| ||# #|| ||# ||* ||*
Optimized  ACC TCT CGT CAC GCG TGG CTG TCT ATC GGT GCG CAG AAC AAA GGT CCG CTG TTC CCG GGT
```

```
           I   P   N   H   F   P   D   S   C   V   S   T   V   V   G   A   M   D   T   S
    Query  ATA CCC AAC CAC TTC CCC GAC TCG TGT GTA TCA ACA GTC GTG GGG GCC ATG GAC ACC TCT
           ||* ||* ||| ||| ||| ||* ||| ||* ||# ||* ||* ||* ||# ||* ||* ||* ||| ||| ||| |||
Optimized  ATC CCG AAC CAC TTC CCG GAC TCT TGC GTT TCT ACC GTT GTT GGT GCG ATG GAC ACC TCT

           L   G   G   R   P   S   T   G   V   C   G   P   A   I   S   F   Q   N   N   G
    Query  CTT GGC GGC CGG CCA TCC ACT GGG GTG TGT GGG CCT GCC ATA AGC TTT CAA AAC AAT GGT
           ||* ||# ||# ||* ||# ||# ||# ||* ||* ||# ||* ||* ||* ||* **# ||# ||# ||| ||# |||
Optimized  CTG GGT GGT CGT CCG TCT ACC GGT GTT TGC GGT CCG GCG ATC TCT TTC CAG AAC AAC GGT

           D   V   Y   E   N   D   T   P   S   V   M   F   A   T   Y   D   P   L   T   P
    Query  GAT GTG TAC GAA AAT GAT ACA CCC TCA GTC ATG TTT GCC ACT TAC GAC CCT CTC ACG CCA
           ||# ||* ||| ||| ||# ||# ||* ||* ||* ||# ||| ||# ||* ||# ||| ||| ||* ||* ||* ||#
Optimized  GAC GTT TAC GAA AAC GAC ACC CCG TCT GTT ATG TTC GCG ACC TAC GAC CCG CTG ACC CCG

           G   T   G   V   A   L   T   N   S   I   N   P   A   S   L   A   L   V   R   I
    Query  GGC ACG GGG GTT GCT CTC ACC AAC AGC ATT AAC CCT GCA TCC CTG GCG TTG GTA CGC ATA
           ||# ||* ||* ||| ||* ||* ||| ||| **# ||# ||| ||* ||# ||# ||| ||| #|| ||* ||# ||*
Optimized  GGT ACC GGT GTT GCG CTG ACC AAC TCT ATC AAC CCG GCG TCT CTG GCG CTG GTT CGT ATC

           S   N   N   D   F   D   T   S   G   F   A   N   D   K   N   V   V   V   Q   M
    Query  TCC AAC AAT GAC TTT GAC ACC AGT GGC TTT GCC AAC GAC AAG AAT GTA GTG GTG CAA ATG
           ||# ||| ||# ||| ||# ||| ||| **| ||# ||# ||* ||| ||| ||# ||# ||* ||* ||* ||# |||
Optimized  TCT AAC AAC GAC TTC GAC ACC TCT GGT TTC GCG AAC GAC AAA AAC GTT GTT GTT CAG ATG

           S   W   E   M   Y   T   G   T   N   Q   I   R   G   Q   V   T   P   M   S   G
    Query  TCA TGG GAG ATG TAC ACA GGC ACC AAC CAA ATT AGA GGT CAA GTC ACA CCA ATG TCT GGC
           ||* ||| ||# ||| ||| ||* ||# ||| ||| ||# ||# *|* ||| ||# ||# ||* ||# ||| ||| ||#

           T   N   Y   T   F   T   S   T   G   A   N   T   L   V   L   W   Q   E   R   M
    Query  ACC AAC TAC ACC TTT ACT TCC ACA GGT GCG AAC ACT CTT GTG CTG TGG CAG GAG CGC ATG
           ||| ||| ||| ||| ||# ||# ||# ||* ||| ||| ||| ||# ||* ||* ||| ||| ||| ||# ||# |||
Optimized  ACC AAC TAC ACC TTC ACC TCT ACC GGT GCG AAC ACC CTG GTT CTG TGG CAG GAA CGT ATG

           L   S   Y   D   G   H   Q   A   I   L   Y   S   S   Q   L   E   R   T   A   E
    Query  CTC AGC TAT GAT GGA CAC CAA GCA ATA CTG TAC TCG TCA CAA TTG GAG AGA ACG GCT GAG
           ||* **# ||# ||# ||* ||| ||# ||# ||* ||| ||| ||* ||* ||# #|| ||# *|* ||* ||* ||#
Optimized  CTG TCT TAC GAC GGT CAC CAG GCG ATC CTG TAC TCT TCT CAG CTG GAA CGT ACC GCG GAA

           Y   F   Q   N   D   I   V   N   I   P   E   N   S   M   A   V   F   N   V   E
    Query  TAC TTC CAG AAT GAC ATA GTC AAC ATA CCA GAA AAT TCC ATG GCG GTG TTC AAT GTG GAA
           ||| ||| ||| ||# ||| ||* ||# ||| ||* ||# ||| ||# ||# ||| ||| ||* ||| ||# ||* |||
Optimized  TAC TTC CAG AAC GAC ATC GTT AAC ATC CCG GAA AAC TCT ATG GCG GTT TTC AAC GTT GAA
```

```
           T   N   S   A   S   F   Q   I   G   I   R   P   D   G   Y   M   V   T   G   G
    Query  ACC AAC TCC GCA TCC TTC CAA ATT GGC ATT AGA CCT GAT GGT TAC ATG GTG ACA GGT GGA
           ||| ||| ||# ||# ||# ||| ||# ||# ||# ||# *|* ||* ||# ||| ||| ||| ||* ||* ||| ||*
Optimized  ACC AAC TCT GCG TCT TTC CAG ATC GGT ATC CGT CCG GAC GGT TAC ATG GTT ACC GGT GGT

           S   I   G   I   N   V   P   L   E   P   E   T   R   F   Q   Y   V   G   I   L
    Query  TCT ATT GGC ATT AAC GTA CCG CTT GAA CCT GAA ACC AGA TTC CAA TAC GTT GGA ATT CTC
           ||| ||# ||# ||# ||| ||* ||| ||* ||| ||* ||| ||| *|* ||| ||# ||| ||| ||* ||# ||*
Optimized  TCT ATC GGT ATC AAC GTT CCG CTG GAA CCG GAA ACC CGT TTC CAG TAC GTT GGT ATC CTG

           P   L   S   A   A   L   S   G   P   S   G   N   M   G   R   A   K   R   V   F
    Query  CCT TTG TCT GCA GCT TTA TCT GGG CCC AGT GGG AAC ATG GGA AGG GCC AAA CGG GTG TTC
           ||* #|| ||| ||# ||* #|# ||| ||* ||* **| ||* ||| ||| ||* *|* ||* ||| ||* ||* |||
Optimized  CCG CTG TCT GCG GCG CTG TCT GGT CCG TCT GGT AAC ATG GGT CGT GCG AAA CGT GTT TTC

           Q   .
    Query  CAA TGA
           ||# |||
Optimized  CAG TGA
```

**Mc114 (AY237422.3)** – 1,686 nucleotides; 1,293 unchanged, 207 transition and 186 transversion changes.

```
           M   E   G   N   G   S   N   S   E   P   K   Q   S   N   N   P   M   V   V   D
    Query  ATG GAG GGC AAT GGC TCC AAC TCA GAG CCA AAG CAG AGC AAC AAC CCA ATG GTC GTT GAC
           ||| ||# ||# ||# ||# ||# ||| ||* ||# ||# ||# ||| **# ||| ||| ||# ||| ||# ||| |||
Optimized  ATG GAA GGT AAC GGT TCT AAC TCT GAA CCG AAA CAG TCT AAC AAC CCG ATG GTT GTT GAC

           P   P   G   T   T   G   P   T   T   S   H   V   V   V   A   N   P   E   Q   P
    Query  CCG CCT GGC ACA ACA GGT CCG ACC ACA TCC CAC GTT GTT GTT GCT AAT CCG GAG CAA CCC
           ||| ||* ||# ||* ||* ||| ||| ||| ||* ||# ||| ||| ||| ||| ||* ||# ||| ||# ||# ||*
Optimized  CCG CCG GGT ACC ACC GGT CCG ACC ACC TCT CAC GTT GTT GTT GCG AAC CCG GAA CAG CCG

           N   G   A   A   Q   R   L   E   L   A   V   A   T   G   A   I   Q   S   N   V
    Query  AAT GGG GCC GCA CAG CGC CTG GAG TTG GCT GTT GCC ACT GGT GCA ATC CAA TCC AAT GTC
           ||# ||* ||* ||# ||| ||# ||| ||# #|| ||* ||| ||* ||# ||| ||# ||| ||# ||# ||# ||#
Optimized  AAC GGT GCG GCG CAG CGT CTG GAA CTG GCG GTT GCG ACC GGT GCG ATC CAG TCT AAC GTT

           P   E   A   I   R   N   C   F   A   V   F   R   T   F   A   W   N   D   R   M
    Query  CCT GAG GCA ATA CGC AAC TGC TTT GCA GTC TTT CGT ACT TTT GCT TGG AAC GAC AGG ATG
           ||* ||# ||# ||* ||# ||| ||| ||# ||# ||# ||# ||| ||# ||# ||* ||| ||| ||| *|* |||
Optimized  CCG GAA GCG ATC CGT AAC TGC TTC GCG GTT TTC CGT ACC TTC GCG TGG AAC GAC CGT ATG

           P   T   G   T   F   L   G   S   I   S   L   H   P   N   I   N   P   Y   T   S
    Query  CCC ACG GGA ACT TTT CTT GGA TCT ATA TCG CTT CAT CCC AAC ATT AAC CCG TAC ACT TCT
           ||* ||* ||* ||# ||# ||* ||* ||| ||* ||* ||* ||# ||* ||| ||# ||| ||| ||| ||# |||
Optimized  CCG ACC GGT ACC TTC CTG GGT TCT ATC TCT CTG CAC CCG AAC ATC AAC CCG TAC ACC TCT

           H   L   S   G   M   W   A   G   W   G   G   S   F   E   V   R   L   S   I   S
    Query  CAC CTC TCT GGG ATG TGG GCC GGG TGG GGC GGC AGT TTT GAG GTC CGG CTA TCG ATC TCT
           ||| ||* ||| ||* ||| ||| ||* ||* ||| ||# ||# **| ||# ||# ||# ||* ||# ||* ||| |||
Optimized  CAC CTG TCT GGT ATG TGG GCG GGT TGG GGT GGT TCT TTC GAA GTT CGT CTG TCT ATC TCT

           G   S   G   V   F   A   G   R   I   I   A   S   V   I   P   P   G   V   D   P
    Query  GGT TCT GGC GTG TTC GCT GGG CGC ATC ATT GCT TCT GTC ATA CCA CCA GGG GTT GAT CCC
           ||| ||| ||# ||* ||| ||* ||* ||# ||| ||# ||* ||| ||# ||* ||# ||# ||* ||| ||# ||*
Optimized  GGT TCT GGT GTT TTC GCG GGT CGT ATC ATC GCG TCT GTT ATC CCG CCG GGT GTT GAC CCG
```

```
          S   S   I   R   D   P   G   V   L   P   H   A   F   V   D   A   R   I   T   E
    Query TCG TCC ATC AGG GAC CCA GGC GTG TTG CCT CAC GCT TTC GTT GAT GCT CGC ATC ACT GAG
          ||* ||# ||| *|* ||| ||# ||# ||* #|| ||* ||| ||* ||| ||| ||# ||* ||# ||| ||# ||#
Optimized TCT TCT ATC CGT GAC CCG GGT GTT CTG CCG CAC GCG TTC GTT GAC GCG CGT ATC ACC GAA

          P   V   S   F   M   I   P   D   V   R   A   V   D   Y   H   R   M   D   G   A
    Query CCA GTT TCT TTC ATG ATC CCT GAT GTA AGG GCA GTG GAC TAC CAC AGG ATG GAT GGT GCT
          ||# ||| ||| ||| ||| ||| ||* ||# ||* *|* ||# ||* ||| ||| ||| *|* ||| ||# ||| ||*
Optimized CCG GTT TCT TTC ATG ATC CCG GAC GTT CGT GCG GTT GAC TAC CAC CGT ATG GAC GGT GCG

          E   P   T   C   S   L   G   F   W   V   Y   Q   P   L   L   N   P   F   S   T
    Query GAA CCC ACA TGC TCC TTG GGG TTT TGG GTT TAT CAA CCA TTA CTC AAC CCC TTT TCC ACC
          ||| ||* ||* ||| ||# #|| ||* ||# ||| ||| ||# ||# ||# #|# ||* ||| ||* ||# ||# |||
Optimized GAA CCG ACC TGC TCT CTG GGT TTC TGG GTT TAC CAG CCG CTG CTG AAC CCG TTC TCT ACC

          T   A   V   S   T   C   W   V   S   V   E   T   K   P   G   G   D   F   D   F
    Query ACC GCC GTC TCA ACA TGT TGG GTG TCA GTG GAA ACT AAA CCA GGT GGT GAT TTT GAC TTC
          ||| ||* ||# ||* ||* ||# ||| ||* ||* ||* ||| ||# ||| ||# ||| ||| ||# ||# ||| |||
Optimized ACC GCG GTT TCT ACC TGC TGG GTT TCT GTT GAA ACC AAA CCG GGT GGT GAC TTC GAC TTC

          C   L   L   R   P   P   G   Q   Q   M   E   N   G   V   S   P   E   G   L   L
    Query TGT TTG CTA AGG CCA CCA GGT CAA CAG ATG GAA AAT GGG GTT TCA CCA GAA GGC TTA TTG
          ||# #|| ||# *|* ||# ||# ||| ||# ||| ||| ||| ||# ||* ||| ||* ||# ||| ||# #|# #||
Optimized TGC CTG CTG CGT CCG CCG GGT CAG CAG ATG GAA AAC GGT GTT TCT CCG GAA GGT CTG CTG

          P   R   R   L   G   Y   S   R   G   N   R   V   G   G   L   V   V   G   M   I
    Query CCT AGG CGC TTG GGT TAC TCG CGT GGT AAC CGT GTT GGT GGC TTG GTG GTT GGC ATG ATT
          ||* *|* ||# #|| ||| ||| ||* ||| ||| ||| ||| ||| ||| ||# #|| ||* ||| ||# ||| ||#
Optimized CCG CGT CGT CTG GGT TAC TCT CGT GGT AAC CGT GTT GGT GGT CTG GTT GTT GGT ATG ATC

          L   V   A   E   H   K   Q   V   N   R   H   F   N   S   N   S   V   T   F   G
    Query TTG GTA GCT GAG CAT AAG CAG GTC AAT AGA CAC TTT AAT TCC AAC TCA GTC ACC TTT GGT
          #|| ||* ||* ||# ||# ||# ||| ||# ||# *|* ||| ||# ||# ||# ||| ||* ||# ||| ||# |||
Optimized CTG GTT GCG GAA CAC AAA CAG GTT AAC CGT CAC TTC AAC TCT AAC TCT GTT ACC TTC GGT

          W   S   T   A   P   V   N   P   M   A   A   E   I   V   T   N   Q   A   H   S
    Query TGG TCA ACG GCA CCT GTT AAT CCA ATG GCA GCA GAG ATT GTG ACT AAC CAG GCA CAT TCA
          ||| ||* ||* ||# ||* ||| ||# ||# ||| ||# ||# ||# ||# ||* ||# ||| ||| ||# ||# ||*
Optimized TGG TCT ACC GCG CCG GTT AAC CCG ATG GCG GCG GAA ATC GTT ACC AAC CAG GCG CAC TCT

          T   S   R   H   A   W   L   S   I   G   A   Q   N   K   G   P   L   F   P   G
    Query ACA TCA CGC CAT GCT TGG CTT TCC ATT GGT GCT CAG AAT AAG GGT CCA TTG TTC CCC GGG
          ||* ||* ||# ||# ||* ||| ||* ||# ||# ||| ||* ||| ||# ||# ||| ||# #|| ||| ||* ||*
Optimized ACC TCT CGT CAC GCG TGG CTG TCT ATC GGT GCG CAG AAC AAA GGT CCG CTG TTC CCG GGT
```

```
          I   P   N   H   F   P   D   S   C   A   S   T   V   V   G   A   M   D   T   S
    Query ATA CCC AAC CAC TTC CCC GAC TCG TGT GCA TCA ACA GTC GTG GGG GCC ATG GAC ACC TCT
          ||* ||* ||| ||| ||| ||* ||| ||* ||# ||# ||* ||* ||# ||* ||* ||* ||| ||| ||| |||
Optimized ATC CCG AAC CAC TTC CCG GAC TCT TGC GCG TCT ACC GTT GTT GGT GCG ATG GAC ACC TCT

          L   G   G   R   P   S   T   G   V   C   G   P   A   I   S   F   Q   N   N   G
    Query CTT GGC GGC CGG CCA TCC ACT GGG GTG TGT GGG CCT GCC ATA AGC TTC CAA AAC AAT GGT
          ||* ||# ||# ||* ||# ||# ||# ||* ||* ||# ||* ||* ||* ||* **# ||| ||# ||| ||# |||
Optimized CTG GGT GGT CGT CCG TCT ACC GGT GTT TGC GGT CCG GCG ATC TCT TTC CAG AAC AAC GGT

          D   V   Y   E   N   D   T   P   S   V   M   F   A   T   Y   D   P   L   T   S
    Query GAT GTG TAC GAA AAT GAT ACA CCC TCA GTC ATG TTT GCC ACT TAC GAC CCC CTC ACG TCA
          ||# ||* ||| ||| ||# ||# ||* ||* ||* ||# ||| ||# ||* ||# ||| ||| ||* ||* ||* ||*
Optimized GAC GTT TAC GAA AAC GAC ACC CCG TCT GTT ATG TTC GCG ACC TAC GAC CCG CTG ACC TCT

          G   T   G   V   A   L   T   N   S   I   N   P   A   S   L   A   L   V   R   I
    Query GGC ACG GGG GTT GCT CTT ACC AAC AGC ATT AAC CCT GCA TCC CTG GCG TTG GTG CGC ATA
          ||# ||* ||* ||| ||* ||* ||| ||| **# ||# ||| ||* ||# ||# ||| ||| #|| ||* ||# ||*
Optimized GGT ACC GGT GTT GCG CTG ACC AAC TCT ATC AAC CCG GCG TCT CTG GCG CTG GTT CGT ATC

          S   N   N   D   F   D   T   S   G   F   A   N   D   K   N   V   V   V   Q   M
    Query TCC AAC AAT GAC TTT GAC ACC AGT GGC TTT GCC AAC GAC AAG AAT GTA GTG GTG CAA ATG
          ||# ||| ||# ||| ||# ||| ||| **| ||# ||# ||* ||| ||| ||# ||# ||* ||* ||* ||# |||
Optimized TCT AAC AAC GAC TTC GAC ACC TCT GGT TTC GCG AAC GAC AAA AAC GTT GTT GTT CAG ATG

          S   W   E   M   Y   T   G   T   N   Q   I   R   G   Q   V   T   P   M   S   G
    Query TCA TGG GAG ATG TAC ACA GGC ACC AAC CAA ATT AGA GGT CAA GTC ACA CCA ATG TCT GGC
          ||* ||| ||# ||| ||| ||* ||# ||| ||| ||# ||# *|* ||| ||# ||# ||* ||# ||| ||| ||#

          T   N   Y   T   F   T   S   T   G   A   N   T   L   V   L   W   Q   E   R   M
    Query ACC AAC TAC ACC TTT ACT TCC ACA GGT GCG AAC ACT CTT GTG CTG TGG CAG GAG CGC ATG
          ||| ||| ||| ||| ||# ||# ||# ||* ||| ||| ||| ||# ||* ||* ||| ||| ||| ||# ||# |||
Optimized ACC AAC TAC ACC TTC ACC TCT ACC GGT GCG AAC ACC CTG GTT CTG TGG CAG GAA CGT ATG

          L   S   Y   D   G   H   Q   A   I   L   Y   S   S   Q   L   E   R   T   A   E
    Query CTT AGC TAT GAT GGG CAC CAA GCA ATA CTG TAC TCG TCA CAA TTG GAG AGA ACG GCT GAG
          ||* **# ||# ||# ||* ||| ||# ||# ||* ||| ||| ||* ||* ||# #|| ||# *|* ||* ||* ||#
Optimized CTG TCT TAC GAC GGT CAC CAG GCG ATC CTG TAC TCT TCT CAG CTG GAA CGT ACC GCG GAA

          Y   F   Q   N   D   I   V   N   I   P   E   N   S   M   A   V   F   N   V   E
    Query TAC TTC CAG AAT GAC ATA GTC AAC ATA CCA GAA AAT TCC ATG GCG GTG TTC AAT GTG GAA
          ||| ||| ||| ||# ||| ||* ||# ||| ||* ||# ||| ||# ||# ||| ||| ||* ||| ||# ||* |||
Optimized TAC TTC CAG AAC GAC ATC GTT AAC ATC CCG GAA AAC TCT ATG GCG GTT TTC AAC GTT GAA
```

```
           T   N   S   A   S   F   Q   I   G   I   R   P   D   G   Y   M   V   T   G   G
     Query ACC AAC TCC GCA TCC TTC CAA ATT GGC ATC AGG CCT GAT GGT TAC ATG GTG ACA GGT GGA
           ||| ||| ||# ||# ||# ||| ||# ||# ||# ||| *|* ||* ||# ||| ||| ||| ||* ||* ||| ||*
 Optimized ACC AAC TCT GCG TCT TTC CAG ATC GGT ATC CGT CCG GAC GGT TAC ATG GTT ACC GGT GGT

           S   I   G   I   N   V   P   L   E   P   E   T   R   F   Q   Y   V   G   I   L
     Query TCT ATT GGC ATC AAC GTG CCG CTT GAA CCT GAA ACC AGA TTC CAA TAC GTT GGA ATT CTC
           ||| ||# ||# ||| ||| ||* ||| ||* ||| ||* ||| ||| *|* ||| ||# ||| ||| ||* ||# ||*
 Optimized TCT ATC GGT ATC AAC GTT CCG CTG GAA CCG GAA ACC CGT TTC CAG TAC GTT GGT ATC CTG

           P   L   S   A   A   L   S   G   P   S   G   N   M   G   R   A   K   R   V   F
     Query CCT TTG TCT GCA GCT TTA TCT GGG CCC AGT GGG AAC ATG GGA AGG GCC AAA CGG GTG TTC
           ||* #|| ||| ||# ||* #|# ||| ||* ||* **| ||* ||| ||| ||* *|* ||* ||| ||* ||* |||
 Optimized CCG CTG TCT GCG GCG CTG TCT GGT CCG TCT GGT AAC ATG GGT CGT GCG AAA CGT GTT TTC

           Q   .
     Query CAA TGA
           ||# |||
 Optimized CAG TGA
```

**NongKhai-50 (AY646853.2)** – 1,686 nucleotides; 1,293 unchanged, 207 transition and 186 transversion changes.

```
               M   E   G   N   G   S   N   P   E   P   K   Q   S   N   N   P   M   V   V   D
    Query     ATG GAG GGC AAT GGC TCC AAC CCA GAG CCA AAG CAA AGC AAC AAC CCG ATG GTC GTT GAC
              ||| ||# ||# ||# ||# ||# ||| ||# ||# ||# ||# ||# **# ||| ||| ||| ||| ||# ||| |||
Optimized     ATG GAA GGT AAC GGT TCT AAC CCG GAA CCG AAA CAG TCT AAC AAC CCG ATG GTT GTT GAC

               P   P   G   T   T   G   P   T   T   S   H   V   V   V   A   N   P   E   Q   P
    Query     CCG CCT GGC ACA ACA GGT CCG ACC ACA TCC CAC GTT GTT GTT GCT AAT CCG GAG CAA CCC
              ||| ||* ||# ||* ||* ||| ||| ||| ||* ||# ||| ||| ||| ||| ||* ||# ||| ||# ||# ||*
Optimized     CCG CCG GGT ACC ACC GGT CCG ACC ACC TCT CAC GTT GTT GTT GCG AAC CCG GAA CAG CCG

               N   G   A   A   Q   R   L   E   L   A   V   A   T   G   A   I   Q   S   N   V
    Query     AAT GGG GCC GCA CAG CGC CTG GAG TTG GCT GTT GCC ACT GGT GCA ATC CAA TCC AAT GTC
              ||# ||* ||* ||# ||| ||# ||| ||# #|| ||* ||| ||* ||# ||| ||# ||| ||# ||# ||# ||#
Optimized     AAC GGT GCG GCG CAG CGT CTG GAA CTG GCG GTT GCG ACC GGT GCG ATC CAG TCT AAC GTT

               P   E   A   I   R   N   C   F   A   V   F   R   T   F   A   W   N   D   R   M
    Query     CCT GAG GCA ATA CGC AAC TGC TTT GCA GTC TTT CGT ACT TTT GCT TGG AAC GAC AGG ATG
              ||* ||# ||# ||* ||# ||| ||| ||# ||# ||# ||# ||| ||# ||# ||* ||| ||| ||| *|* |||
Optimized     CCG GAA GCG ATC CGT AAC TGC TTC GCG GTT TTC CGT ACC TTC GCG TGG AAC GAC CGT ATG

               P   T   G   T   F   L   G   S   I   S   L   H   P   N   I   N   P   Y   T   A
    Query     CCC ACG GGA ACT TTT CTT GGA TCT ATA TCG CTT CAT CCC AAC ATT AAC CCG TAC ACT GCT
              ||* ||* ||* ||# ||# ||* ||* ||| ||* ||* ||* ||# ||* ||| ||# ||| ||| ||| ||# ||*
Optimized     CCG ACC GGT ACC TTC CTG GGT TCT ATC TCT CTG CAC CCG AAC ATC AAC CCG TAC ACC GCG

               H   L   S   G   M   W   A   G   W   G   G   S   F   E   V   R   L   S   I   S
    Query     CAC CTC TCT GGG ATG TGG GCC GGG TGG GGC GGT AGT TTT GAG GTC CGG CTA TCG ATC TCT
              ||| ||* ||| ||* ||| ||| ||* ||* ||| ||# ||| **| ||# ||# ||# ||* ||# ||* ||| |||
Optimized     CAC CTG TCT GGT ATG TGG GCG GGT TGG GGT GGT TCT TTC GAA GTT CGT CTG TCT ATC TCT

               G   S   G   V   F   A   G   R   I   I   A   S   V   I   P   P   G   V   D   P
    Query     GGT TCT GGC GTG TTC GCT GGG CGC ATC ATT GCT TCT GTC ATA CCA CCA GGG GTT GAT CCC
              ||| ||| ||# ||* ||| ||* ||* ||# ||| ||# ||* ||| ||# ||* ||# ||# ||* ||| ||# ||*
Optimized     GGT TCT GGT GTT TTC GCG GGT CGT ATC ATC GCG TCT GTT ATC CCG CCG GGT GTT GAC CCG
```

```
          S   S   I   R   D   P   G   V   L   P   H   A   F   V   D   A   R   I   T   E
    Query TCG TCC ATC AGG GAC CCA GGC GTG TTG CCT CAC GCT TTC GTT GAT GCT CGC ATC ACT GAG
          ||* ||# ||| *|* ||| ||# ||# ||* #|| ||* ||| ||* ||| ||| ||# ||* ||# ||| ||# ||#
Optimized TCT TCT ATC CGT GAC CCG GGT GTT CTG CCG CAC GCG TTC GTT GAC GCG CGT ATC ACC GAA

          P   V   S   F   M   I   P   D   V   R   A   V   D   Y   H   R   M   D   G   A
    Query CCA GTT TCT TTC ATG ATC CCT GAT GTC AGA GCA GTG GAC TAC CAC AGG ATG GAT GGT GCT
          ||# ||| ||| ||| ||| ||| ||* ||# ||# *|* ||# ||* ||| ||| ||| *|* ||| ||# ||| ||*
Optimized CCG GTT TCT TTC ATG ATC CCG GAC GTT CGT GCG GTT GAC TAC CAC CGT ATG GAC GGT GCG

          E   P   T   C   S   L   G   F   W   V   Y   Q   P   L   L   N   P   F   S   T
    Query GAG CCC ACT TGC TCG TTA GGG TTC TGG GTT TAC CAA CCA TTG CTC AAC CCA TTT TCC ACC
          ||# ||* ||# ||| ||* #|# ||* ||| ||| ||| ||| ||# ||# #|| ||* ||| ||# ||# ||# |||
Optimized GAA CCG ACC TGC TCT CTG GGT TTC TGG GTT TAC CAG CCG CTG CTG AAC CCG TTC TCT ACC

          T   A   V   S   T   C   W   V   S   V   E   T   K   P   G   G   D   F   D   F
    Query ACC GCC GTT TCA ACA TGT TGG GTG TCA GTA GAG ACT AAG CCA GGT GGT GAT TTT GAT TTC
          ||| ||* ||| ||* ||* ||# ||| ||* ||* ||* ||# ||# ||# ||# ||| ||| ||# ||# ||# |||
Optimized ACC GCG GTT TCT ACC TGC TGG GTT TCT GTT GAA ACC AAA CCG GGT GGT GAC TTC GAC TTC

          C   L   L   R   P   P   G   Q   Q   M   E   N   G   V   S   P   E   G   L   L
    Query TGC CTA CTA AGA CCA CCG GGT CAA CAG ATG GAA AAT GGG GTC TCA CCA GAA GGC CTA TTG
          ||| ||# ||# *|* ||# ||| ||| ||# ||| ||| ||| ||# ||* ||# ||* ||# ||| ||# ||# #||
Optimized TGC CTG CTG CGT CCG CCG GGT CAG CAG ATG GAA AAC GGT GTT TCT CCG GAA GGT CTG CTG

          P   R   R   L   G   Y   S   R   G   N   R   V   G   G   L   V   V   G   M   I
    Query CCC AGA CGC CTA GGT TAC TCG CGT GGT AAT CGT GTT GGT GGC TTG GTG GTT GGT ATG ATC
          ||* *|* ||# ||# ||| ||| ||* ||| ||| ||# ||| ||| ||| ||# #|| ||* ||| ||| ||| |||
Optimized CCG CGT CGT CTG GGT TAC TCT CGT GGT AAC CGT GTT GGT GGT CTG GTT GTT GGT ATG ATC

          L   V   A   E   H   K   Q   V   N   R   H   F   N   S   N   S   V   T   F   G
    Query TTG GTG GCT GAA CAT AAG CAG GTC AAT AGA CAC TTC AAC TCC AAT TCA GTC ACT TTT GGT
          #|| ||* ||* ||| ||# ||# ||| ||# ||# *|* ||| ||| ||| ||# ||# ||* ||# ||# ||# |||
Optimized CTG GTT GCG GAA CAC AAA CAG GTT AAC CGT CAC TTC AAC TCT AAC TCT GTT ACC TTC GGT

          W   S   T   A   P   V   N   P   M   A   A   E   I   V   T   N   Q   A   H   S
    Query TGG TCC ACG GCA CCT GTC AAC CCA ATG GCA GCA GAA ATT GTG ACA AAC CAG GCA CAT TCA
          ||| ||# ||* ||# ||* ||# ||| ||# ||| ||# ||# ||| ||# ||* ||* ||| ||| ||# ||# ||*
Optimized TGG TCT ACC GCG CCG GTT AAC CCG ATG GCG GCG GAA ATC GTT ACC AAC CAG GCG CAC TCT

          T   S   R   H   A   W   L   S   I   G   A   Q   N   K   G   P   L   F   P   G
    Query ACA TCA CGA CAT GCT TGG CTT TCC ATT GGT GCT CAA AAC AAG GGT CCA TTG TTT CCT GGA
          ||* ||* ||* ||# ||* ||| ||* ||# ||# ||| ||* ||# ||| ||# ||| ||# #|| ||# ||* ||*
Optimized ACC TCT CGT CAC GCG TGG CTG TCT ATC GGT GCG CAG AAC AAA GGT CCG CTG TTC CCG GGT
```

```
           I   P   N   H   F   P   D   S   C   A   S   T   I   V   G   A   M   D   T   S
    Query  ATA CCT AAC CAC TTC CCC GAC TCG TGT GCA TCA ACA ATC GTT GGG GCC ATG GAC ACC TCC
           ||* ||* ||| ||| ||| ||* ||| ||* ||# ||# ||* ||* ||| ||| ||* ||* ||| ||| ||| ||#
Optimized  ATC CCG AAC CAC TTC CCG GAC TCT TGC GCG TCT ACC ATC GTT GGT GCG ATG GAC ACC TCT

           L   G   G   R   P   S   T   G   V   C   G   P   A   I   S   F   Q   N   N   G
    Query  CTT GGA GGC CGG CCG TCC ACT GGG GTG TGT GGA CCT GCC ATA AGC TTT CAA AAT AAT GGT
           ||* ||* ||# ||* ||| ||# ||# ||* ||* ||# ||* ||* ||* ||* **# ||# ||# ||# ||# |||
Optimized  CTG GGT GGT CGT CCG TCT ACC GGT GTT TGC GGT CCG GCG ATC TCT TTC CAG AAC AAC GGT

           D   V   Y   E   N   D   T   P   S   V   M   F   A   T   Y   D   P   L   T   S
    Query  GAT GTG TAC GAG AAT GAC ACA CCC TCA GTC ATG TTT GCC ACT TAT GAC CCA CTC ACG TCA
           ||# ||* ||| ||# ||# ||| ||* ||* ||* ||# ||| ||# ||* ||# ||# ||| ||# ||* ||* ||*
Optimized  GAC GTT TAC GAA AAC GAC ACC CCG TCT GTT ATG TTC GCG ACC TAC GAC CCG CTG ACC TCT

           G   T   G   V   A   L   T   N   S   I   N   P   A   S   L   A   L   V   R   I
    Query  GGC ACG GGG GTT GCT CTT ACC AAC AGC ATC AAC CCC GCA TCC CTG GCG TTG GTG CGC ATA
           ||# ||* ||* ||| ||* ||* ||| ||| **# ||| ||| ||* ||# ||# ||| ||| #|| ||* ||# ||*
Optimized  GGT ACC GGT GTT GCG CTG ACC AAC TCT ATC AAC CCG GCG TCT CTG GCG CTG GTT CGT ATC

           S   N   N   D   F   D   T   S   G   F   A   N   D   K   N   V   V   V   Q   M
    Query  TCC AAT AAT GAC TTT GAC ACT AGT GGC TTT GCC AAC GAT AAG AAC GTG GTG GTT CAA ATG
           ||# ||# ||# ||| ||# ||| ||# **| ||# ||# ||* ||| ||# ||# ||| ||* ||* ||| ||# |||
Optimized  TCT AAC AAC GAC TTC GAC ACC TCT GGT TTC GCG AAC GAC AAA AAC GTT GTT GTT CAG ATG

           S   W   E   M   Y   T   G   T   N   Q   I   R   G   Q   V   T   P   M   S   G
    Query  TCA TGG GAG ATG TAT ACA GGC ACC AAT CAA ATT AGA GGT CAG GTT ACA CCA ATG TCT GGC
           ||* ||| ||# ||| ||# ||* ||# ||| ||# ||# ||# *|* ||| ||| ||| ||* ||# ||| ||| ||#

           T   N   Y   T   F   T   S   T   G   A   N   T   L   V   L   W   Q   E   R   M
    Query  ACC AAC TAC ACC TTC ACA TCA ACA GGT GCA AAC ACT CTT GTG TTG TGG CAG GAG CGT ATG
           ||| ||| ||| ||| ||| ||* ||* ||* ||| ||# ||| ||# ||* ||* #|| ||| ||| ||# ||| |||
Optimized  ACC AAC TAC ACC TTC ACC TCT ACC GGT GCG AAC ACC CTG GTT CTG TGG CAG GAA CGT ATG

           L   S   Y   D   G   H   Q   A   I   L   Y   S   S   Q   L   E   R   T   A   E
    Query  CTC AGC TAT GAT GGA CAC CAA GCA ATA CTG TAC TCG TCA CAA TTG GAA AGG ACG GCT GAG
           ||* **# ||# ||# ||* ||| ||# ||# ||* ||| ||| ||* ||* ||# #|| ||| *|* ||* ||* ||#
Optimized  CTG TCT TAC GAC GGT CAC CAG GCG ATC CTG TAC TCT TCT CAG CTG GAA CGT ACC GCG GAA

           Y   F   Q   N   D   I   V   N   I   P   E   N   S   M   A   V   F   N   V   E
    Query  TAC TTT CAG AAT GAC ATA GTT AAC ATA CCT GAA AAC TCC ATG GCA GTG TTC AAT GTG GAA
           ||| ||# ||| ||# ||| ||* ||| ||| ||* ||* ||| ||| ||# ||| ||# ||* ||| ||# ||* |||
Optimized  TAC TTC CAG AAC GAC ATC GTT AAC ATC CCG GAA AAC TCT ATG GCG GTT TTC AAC GTT GAA
```

```
           T   N   S   A   S   F   Q   I   G   I   R   P   D   G   Y   M   V   T   G   G
    Query  ACC AAC TCT GCA TCT TTC CAA ATT GGC ATT AGA CCT GAT GGT TAC ATG GTG ACA GGT GGC
           ||| ||| ||| ||# ||| ||| ||# ||# ||# ||# *|* ||* ||# ||| ||| ||| ||* ||* ||| ||#
Optimized  ACC AAC TCT GCG TCT TTC CAG ATC GGT ATC CGT CCG GAC GGT TAC ATG GTT ACC GGT GGT

           S   I   G   I   N   V   P   L   E   P   E   T   R   F   Q   Y   V   G   I   L
    Query  TCC ATT GGT ATT AAT GTG CCG CTT GAG CCT GAA ACC CGA TTC CAA TAC GTT GGA ATT CTC
           ||# ||# ||| ||# ||# ||* ||| ||* ||# ||* ||| ||| ||* ||| ||# ||| ||| ||* ||# ||*
Optimized  TCT ATC GGT ATC AAC GTT CCG CTG GAA CCG GAA ACC CGT TTC CAG TAC GTT GGT ATC CTG

           P   L   S   A   A   L   S   G   P   S   G   N   M   G   R   A   R   R   V   F
    Query  CCT TTG TCC GCA GCT TTG TCT GGG CCC AGT GGG AAC ATG GGA AGG GCC AGA CGG GTG TTC
           ||* #|| ||# ||# ||* #|| ||| ||* ||* **| ||* ||| ||| ||* *|* ||* *|* ||* ||* |||
Optimized  CCG CTG TCT GCG GCG CTG TCT GGT CCG TCT GGT AAC ATG GGT CGT GCG CGT CGT GTT TTC

           Q   .
    Query  CAA TGA
           ||# |||
Optimized  CAG TGA
```

**Chanthaburi-74 (AY646854.2)** – 1,686 nucleotides; 1,300 unchanged, 202 transition and 184 transversion changes.

M E G N G S N P E P K Q S N N P M V V D
Query ATG GAG GGC AAT GGC TCC AAC CCA GAG CCA AAG CAG AGC AAC AAC CCG ATG GTC GTT GAC
||| ||# ||# ||# ||# ||# ||| ||# ||# ||# ||# ||| **# ||| ||| ||| ||| ||# ||| |||
Optimized ATG GAA GGT AAC GGT TCT AAC CCG GAA CCG AAA CAG TCT AAC AAC CCG ATG GTT GTT GAC

P P G T T G P T T S H V V V A N P E Q P
Query CCG CCT GGC ACA ACA GGT CCG ACC ACA TCC CAC GTT GTT GTT GCT AAT CCG GAG CAA CCC
||| ||* ||# ||* ||* ||| ||| ||| ||* ||# ||| ||| ||| ||| ||* ||# ||| ||# ||# ||*
Optimized CCG CCG GGT ACC ACC GGT CCG ACC ACC TCT CAC GTT GTT GTT GCG AAC CCG GAA CAG CCG

N G A A Q R L E L A V A T G A I Q S N V
Query AAT GGG GCC GCA CAG CGC CTG GAG CTG GCT GTT GCC ACT GGT GCA ATC CAA TCC AAT GTC
||# ||* ||* ||# ||| ||# ||| ||# ||| ||* ||| ||* ||# ||| ||# ||| ||# ||# ||# ||#
Optimized AAC GGT GCG GCG CAG CGT CTG GAA CTG GCG GTT GCG ACC GGT GCG ATC CAG TCT AAC GTT

P E A I R N C F A V F R T F A W N D R M
Query CCT GAG GCA ATA CGC AAC TGC TTT GCA GTC TTT CGT ACT TTT GCT TGG AAC GAC AGG ATG
||* ||# ||# ||* ||# ||| ||| ||# ||# ||# ||# ||| ||# ||# ||* ||| ||| ||| *|* |||
Optimized CCG GAA GCG ATC CGT AAC TGC TTC GCG GTT TTC CGT ACC TTC GCG TGG AAC GAC CGT ATG

P T G T F L G S I S L H P N I N P Y T A
Query CCC ACG GGA ACT TTT CTT GGA TCT ATA TCG CTT CAT CCC AAC ATT AAC CCG TAC ACT GCT
||* ||* ||* ||# ||# ||* ||* ||| ||* ||* ||* ||# ||* ||| ||# ||| ||| ||| ||# ||*
Optimized CCG ACC GGT ACC TTC CTG GGT TCT ATC TCT CTG CAC CCG AAC ATC AAC CCG TAC ACC GCG

H L S G M W A G W G G S F E V R L S I S
Query CAC CTC TCT GGG ATG TGG GCC GGG TGG GGC GGT AGT TTT GAG GTC CGG CTA TCG ATC TCT
||| ||* ||| ||* ||| ||| ||* ||* ||| ||# ||| **| ||# ||# ||# ||* ||# ||* ||| |||
Optimized CAC CTG TCT GGT ATG TGG GCG GGT TGG GGT GGT TCT TTC GAA GTT CGT CTG TCT ATC TCT

G S G V F A G R I I A S V I P P G V D P
Query GGT TCT GGC GTG TTC GCT GGG CGC ATC ATT GCT TCT GTC ATA CCA CCA GGG GTT GAT CCC
||| ||| ||# ||* ||| ||* ||* ||# ||| ||# ||* ||| ||# ||* ||# ||# ||* ||| ||# ||*
Optimized GGT TCT GGT GTT TTC GCG GGT CGT ATC ATC GCG TCT GTT ATC CCG CCG GGT GTT GAC CCG

```
          S   S   I   R   D   P   G   V   L   P   H   A   F   V   D   A   R   I   T   E
Query     TCG TCC ATC AGG GAC CCA GGC GTG TTG CCT CAC GCT TTC GTT GAT GCT CGC ATC ACT GAG
          ||* ||# ||| *|* ||| ||# ||# ||* #|| ||* ||| ||* ||| ||| ||# ||* ||# ||| ||# ||#
Optimized TCT TCT ATC CGT GAC CCG GGT GTT CTG CCG CAC GCG TTC GTT GAC GCG CGT ATC ACC GAA

          P   V   S   F   M   I   P   D   V   R   A   V   D   Y   H   R   M   D   G   A
Query     CCA GTT TCT TTC ATG ATC CCT GAT GTC AGA GCA GTG GAC TAC CAC AGG ATG GAT GGT GCT
          ||# ||| ||| ||| ||| ||| ||* ||# ||# *|* ||# ||* ||| ||| ||| *|* ||| ||# ||| ||*
Optimized CCG GTT TCT TTC ATG ATC CCG GAC GTT CGT GCG GTT GAC TAC CAC CGT ATG GAC GGT GCG

          E   P   T   C   S   L   G   F   W   V   Y   Q   P   L   L   N   P   F   S   T
Query     GAG CCC ACT TGC TCG TTA GGA TTT TGG GTT TAC CAA CCA TTG CTC AAC CCG TTT TCC ACC
          ||# ||* ||# ||| ||* #|# ||* ||# ||| ||| ||| ||# ||# #|| ||* ||| ||| ||# ||# |||
Optimized GAA CCG ACC TGC TCT CTG GGT TTC TGG GTT TAC CAG CCG CTG CTG AAC CCG TTC TCT ACC

          T   A   V   S   T   C   W   V   S   V   E   T   K   P   G   G   D   F   D   F
Query     ACC GCC GTT TCA ACA TGC TGG GTG TCA GTG GAG ACC AAA CCA GGT GGT GAT TTT GAT TTT
          ||| ||* ||| ||* ||* ||| ||| ||* ||* ||* ||# ||| ||| ||# ||| ||| ||# ||# ||# ||#
Optimized ACC GCG GTT TCT ACC TGC TGG GTT TCT GTT GAA ACC AAA CCG GGT GGT GAC TTC GAC TTC

          C   L   L   R   P   P   G   Q   Q   M   E   N   G   V   S   P   E   G   L   L
Query     TGC CTA CTA AGG CCA CCA GGT CAG CAG ATG GAA AAT GGG GTT TCA CCA GAA GGC CTG TTG
          ||| ||# ||# *|* ||# ||# ||| ||| ||| ||| ||| ||# ||* ||| ||* ||# ||| ||# ||| #||
Optimized TGC CTG CTG CGT CCG CCG GGT CAG CAG ATG GAA AAC GGT GTT TCT CCG GAA GGT CTG CTG

          P   R   R   L   G   Y   S   R   G   N   R   V   G   G   L   V   V   G   M   I
Query     CCC AGG CGC CTA GGT TAC TCA CGC GGT AAC CGT GTT GGT GGC TTG GTG GTT GGT ATG ATC
          ||* *|* ||# ||# ||| ||| ||* ||# ||| ||| ||| ||| ||| ||# #|| ||* ||| ||| ||| |||
Optimized CCG CGT CGT CTG GGT TAC TCT CGT GGT AAC CGT GTT GGT GGT CTG GTT GTT GGT ATG ATC

          L   V   A   E   H   K   Q   V   N   R   H   F   N   S   N   S   V   T   F   G
Query     TTG GTG GCT GAG CAC AAG CAG GTC AAC AGG CAT TTT AAC TCC AAT TCA GTC ACT TTT GGT
          #|| ||* ||* ||# ||| ||# ||| ||# ||| *|* ||# ||# ||| ||# ||# ||* ||# ||# ||# |||
Optimized CTG GTT GCG GAA CAC AAA CAG GTT AAC CGT CAC TTC AAC TCT AAC TCT GTT ACC TTC GGT

          W   S   T   A   P   V   N   P   M   A   A   E   I   V   T   N   Q   A   H   S
Query     TGG TCC ACG GCA CCT GTC AAC CCA ATG GCG GCA GAA ATT GTG ACA AAC CAA GCA CAT TCA
          ||| ||# ||* ||# ||* ||# ||| ||# ||| ||| ||# ||| ||# ||* ||* ||| ||# ||# ||# ||*
Optimized TGG TCT ACC GCG CCG GTT AAC CCG ATG GCG GCG GAA ATC GTT ACC AAC CAG GCG CAC TCT

          T   S   R   H   A   W   L   S   I   G   A   Q   N   K   G   P   L   F   P   G
Query     ACA TCG CGC CAT GCT TGG CTT TCT ATT GGT GCT CAA AAC AAG GGT CCA TTG TTT CCC GGG
          ||* ||* ||# ||# ||* ||| ||* ||| ||# ||| ||* ||# ||| ||# ||| ||# #|| ||# ||* ||*
Optimized ACC TCT CGT CAC GCG TGG CTG TCT ATC GGT GCG CAG AAC AAA GGT CCG CTG TTC CCG GGT
```

```
                I   P   N   H   F   P   D   S   C   A   S   T   I   V   G   A   M   D   T   S
    Query       ATA CCT AAC CAT TTC CCC GAC TCG TGT GCA TCA ACA ATC GTG GGG GCC ATG GAC ACC TCC
                ||* ||* ||| ||# ||| ||* ||| ||* ||# ||# ||* ||* ||| ||* ||* ||* ||| ||| ||| ||#
Optimized       ATC CCG AAC CAC TTC CCG GAC TCT TGC GCG TCT ACC ATC GTT GGT GCG ATG GAC ACC TCT

                L   G   G   R   P   S   T   G   V   C   G   P   A   I   S   F   Q   N   N   G
    Query       CTT GGT GGC CGG CCG TCT ACT GGG GTA TGT GGA CCT GCC ATA AGT TTT CAA AAC AAT GGT
                ||* ||| ||# ||* ||| ||| ||# ||* ||* ||# ||* ||* ||* ||* **| ||# ||# ||| ||# |||
Optimized       CTG GGT GGT CGT CCG TCT ACC GGT GTT TGC GGT CCG GCG ATC TCT TTC CAG AAC AAC GGT

                D   V   Y   E   N   D   T   P   S   V   M   F   A   T   Y   D   P   L   T   S
    Query       GAT GTG TAC GAG AAC GAC ACA CCC TCA GTC ATG TTT GCC ACT TAT GAC CCA CTT ACG TCA
                ||# ||* ||| ||# ||| ||| ||* ||* ||* ||# ||| ||# ||* ||# ||# ||| ||# ||* ||* ||*
Optimized       GAC GTT TAC GAA AAC GAC ACC CCG TCT GTT ATG TTC GCG ACC TAC GAC CCG CTG ACC TCT

                G   T   G   V   A   L   T   N   S   I   N   P   A   S   L   A   L   V   R   I
    Query       GGC ACG GGG GTT GCT CTT ACC AAC AGC ATC AAC CCC GCA TCC CTG GCG TTG GTG CGC ATA
                ||# ||* ||* ||| ||* ||* ||| ||| **# ||| ||| ||* ||# ||# ||| ||| #|| ||* ||# ||*
Optimized       GGT ACC GGT GTT GCG CTG ACC AAC TCT ATC AAC CCG GCG TCT CTG GCG CTG GTT CGT ATC

                S   N   N   D   F   D   T   S   G   F   A   N   D   K   N   V   V   V   Q   M
    Query       TCT AAT AAT GAC TTT GAC ACT AGT GGC TTT GCC AAC GAT AAG AAC GTG GTG GTT CAA ATG
                ||| ||# ||# ||| ||# ||| ||# **| ||# ||# ||* ||| ||# ||# ||| ||* ||* ||| ||# |||
Optimized       TCT AAC AAC GAC TTC GAC ACC TCT GGT TTC GCG AAC GAC AAA AAC GTT GTT GTT CAG ATG

                S   W   E   M   Y   T   G   T   N   Q   I   R   G   Q   V   T   P   M   S   G
    Query       TCA TGG GAA ATG TAT ACA GGC ACC AAT CAA ATT AGA GGT CAG GTC ACA CCA ATG TCT GGC
                ||* ||| ||| ||| ||# ||* ||# ||| ||# ||# ||# *|* ||| ||| ||# ||* ||# ||| ||| ||#

                T   N   Y   T   F   T   S   T   G   A   N   T   L   V   L   W   Q   E   R   M
    Query       ACT AAC TAC ACC TTC ACA TCC ACA GGT GCA AAC ACC CTT GTA TTG TGG CAA GAG CGC ATG
                ||# ||| ||| ||| ||| ||* ||# ||* ||| ||# ||| ||| ||* ||* #|| ||| ||# ||# ||# |||
Optimized       ACC AAC TAC ACC TTC ACC TCT ACC GGT GCG AAC ACC CTG GTT CTG TGG CAG GAA CGT ATG

                L   S   Y   D   G   H   Q   A   I   L   Y   S   S   Q   L   E   R   T   A   E
    Query       CTC AGC TAT GAT GGA CAC CAA GCA ATA CTG TAC TCG TCA CAA TTG GAG AGG ACG GCC GAG
                ||* **# ||# ||# ||* ||| ||# ||# ||* ||| ||| ||* ||* ||# #|| ||# *|* ||* ||* ||#
Optimized       CTG TCT TAC GAC GGT CAC CAG GCG ATC CTG TAC TCT TCT CAG CTG GAA CGT ACC GCG GAA

                Y   F   Q   N   D   I   V   N   I   P   E   N   S   M   A   V   F   N   V   E
    Query       TAC TTT CAG AAT GAT ATA GTC AAC ATA CCT GAA AAC TCT ATG GCA GTG TTC AAT GTG GAA
                ||| ||# ||| ||# ||# ||* ||# ||| ||* ||* ||| ||| ||| ||| ||# ||* ||| ||# ||* |||
Optimized       TAC TTC CAG AAC GAC ATC GTT AAC ATC CCG GAA AAC TCT ATG GCG GTT TTC AAC GTT GAA
```

```
          T   N   S   A   S   F   Q   I   G   I   R   P   D   G   Y   M   V   T   G   G
Query     ACC AAC TCT GCA TCT TTC CAA ATT GGC ATC AGA CCT GAT GGT TAT ATG GTG ACA GGT GGC
          ||| ||| ||| ||# ||| ||| ||# ||# ||# ||| *|* ||* ||# ||| ||# ||| ||* ||* ||| ||#
Optimized ACC AAC TCT GCG TCT TTC CAG ATC GGT ATC CGT CCG GAC GGT TAC ATG GTT ACC GGT GGT

          S   I   G   I   N   V   P   L   E   P   E   T   R   F   Q   Y   V   G   I   L
Query     TCC ATT GGT ATC AAT GTG CCG CTC GAA CCC GAA ACC CGA TTC CAA TAC GTT GGA ATT CTC
          ||# ||# ||| ||| ||# ||* ||| ||* ||| ||* ||| ||| ||* ||| ||# ||| ||| ||* ||# ||*
Optimized TCT ATC GGT ATC AAC GTT CCG CTG GAA CCG GAA ACC CGT TTC CAG TAC GTT GGT ATC CTG

          P   L   S   A   A   L   S   G   P   S   G   N   M   G   R   A   R   R   V   F
Query     CCT TTG TCC GCA GCT TTA TCT GGG CCC AGT GGG AAC ATG GGA AGG GCC AGA CGG GTG TTC
          ||* #|| ||# ||# ||* #|# ||| ||* ||* **| ||* ||| ||| ||* *|* ||* *|* ||* ||* |||
Optimized CCG CTG TCT GCG GCG CTG TCT GGT CCG TCT GGT AAC ATG GGT CGT GCG CGT CGT GTT TTC

          Q   .
Query     CAA TGA
          ||# |||
Optimized CAG TGA
```

**ROK62 (KP298674.1)** – 1,686 nucleotides; 1,306 unchanged, 195 transition and 185 transversion changes.

```
           M   E   G   N   G   S   N   S   E   P   K   Q   S   N   N   P   M   V   V   D
    Query  ATG GAG GGC AAT GGC TCC AAC TCA GAG CCA AAA CAG AGC AAC AAC CCA ATG GTC GTT GAC
           ||| ||# ||# ||# ||# ||# ||| ||* ||# ||# ||| ||| **# ||| ||| ||# ||| ||# ||| |||
Optimized  ATG GAA GGT AAC GGT TCT AAC TCT GAA CCG AAA CAG TCT AAC AAC CCG ATG GTT GTT GAC

           P   P   G   T   T   G   P   T   T   S   H   V   V   V   A   N   P   E   Q   P
    Query  CCG CCT GGC ACA ACA GGT CCG ACC ACA TCC CAC GTT GTT GTT GCT AAT CCG GAG CAA CCC
           ||| ||* ||# ||* ||* ||| ||| ||| ||* ||# ||| ||| ||| ||| ||* ||# ||| ||# ||# ||*
Optimized  CCG CCG GGT ACC ACC GGT CCG ACC ACC TCT CAC GTT GTT GTT GCG AAC CCG GAA CAG CCG

           N   G   A   A   Q   R   L   E   L   A   V   A   T   G   A   I   Q   S   N   V
    Query  AAT GGG GCC GCA CAG CGC CTG GAG TTG GCT GTT GCC ACT GGT GCA ATC CAA TCC AAT GTC
           ||# ||* ||* ||# ||| ||# ||| ||# #|| ||* ||| ||* ||# ||| ||# ||| ||# ||# ||# ||#
Optimized  AAC GGT GCG GCG CAG CGT CTG GAA CTG GCG GTT GCG ACC GGT GCG ATC CAG TCT AAC GTT

           P   E   A   I   R   N   C   F   A   V   F   R   T   F   A   W   N   D   R   M
    Query  CCT GAG GCA ATA CGC AAC TGC TTT GCA GTC TTT CGT ACT TTT GCT TGG AAC GAC AGG ATG
           ||* ||# ||# ||* ||# ||| ||| ||# ||# ||# ||# ||| ||# ||# ||* ||| ||| ||| *|* |||
Optimized  CCG GAA GCG ATC CGT AAC TGC TTC GCG GTT TTC CGT ACC TTC GCG TGG AAC GAC CGT ATG

           P   T   G   T   F   L   G   S   I   S   L   H   P   N   I   N   P   Y   T   S
    Query  CCC ACG GGA ACT TTT CTT GGA TCT ATA TCG CTT CAT CCC AAC ATT AAC CCG TAC ACT TCT
           ||* ||* ||* ||# ||# ||* ||* ||| ||* ||* ||* ||# ||* ||| ||# ||| ||| ||| ||# |||
Optimized  CCG ACC GGT ACC TTC CTG GGT TCT ATC TCT CTG CAC CCG AAC ATC AAC CCG TAC ACC TCT

           H   L   S   G   M   W   A   G   W   G   G   S   F   E   V   R   L   S   I   S
    Query  CAC CTC TCT GGG ATG TGG GCC GGG TGG GGC GGT AGC TTT GAG GTC CGG CTA TCG ATC TCT
           ||| ||* ||| ||* ||| ||| ||* ||* ||| ||# ||| **# ||# ||# ||# ||* ||# ||* ||| |||
Optimized  CAC CTG TCT GGT ATG TGG GCG GGT TGG GGT GGT TCT TTC GAA GTT CGT CTG TCT ATC TCT

           G   S   G   V   F   A   G   R   I   I   A   S   V   I   P   P   G   V   D   P
    Query  GGC TCT GGC GTG TTC GCT GGG CGC ATC ATT GCT TCT GTC ATA CCA CCA GGG GTT GAT CCC
           ||# ||| ||# ||* ||| ||* ||* ||# ||| ||# ||* ||| ||# ||* ||# ||# ||* ||| ||# ||*
Optimized  GGT TCT GGT GTT TTC GCG GGT CGT ATC ATC GCG TCT GTT ATC CCG CCG GGT GTT GAC CCG
```

```
           S   S   I   R   D   P   G   V   L   P   H   A   F   V   D   A   R   I   T   E
    Query  TCG TCC ATC AGG GAC CCA GGC GTG TTG CCT CAC GCT TTC GTT GAT GCT CGC ATC ACT GAG
           ||* ||# ||| *|* ||| ||# ||# ||* #|| ||* ||| ||* ||| ||| ||# ||* ||# ||| ||# ||#
Optimized  TCT TCT ATC CGT GAC CCG GGT GTT CTG CCG CAC GCG TTC GTT GAC GCG CGT ATC ACC GAA

           P   V   S   F   M   I   P   D   V   R   A   V   D   Y   H   R   M   D   G   A
    Query  CCA GTT TCT TTC ATG ATT CCT GAT GTC AGA GCA GTG GAC TAC CAT AGA ATG GAT GGT GCT
           ||# ||| ||| ||| ||| ||# ||* ||# ||# *|* ||# ||* ||| ||| ||# *|* ||| ||# ||| ||*
Optimized  CCG GTT TCT TTC ATG ATC CCG GAC GTT CGT GCG GTT GAC TAC CAC CGT ATG GAC GGT GCG

           E   P   T   C   S   L   G   F   W   V   Y   Q   P   L   L   N   P   F   S   T
    Query  GAA CCC ACA TGT TCC TTG GGG TTC TGG GTC TAC CAA CCA TTA CTC AAC CCC TTC TCC ACC
           ||| ||* ||* ||# ||# #|| ||* ||| ||| ||# ||| ||# ||# #|# ||* ||| ||* ||| ||# |||
Optimized  GAA CCG ACC TGC TCT CTG GGT TTC TGG GTT TAC CAG CCG CTG CTG AAC CCG TTC TCT ACC

           T   A   V   S   T   C   W   V   S   V   E   T   K   P   G   G   D   F   D   F
    Query  ACC GCC GTG TCA ACA TGT TGG GTG TCA GTG GAA ACT AAA CCA GGT GGT GAT TTT GAC TTC
           ||| ||* ||* ||* ||* ||# ||| ||* ||* ||* ||| ||# ||| ||# ||| ||| ||# ||# ||| |||
Optimized  ACC GCG GTT TCT ACC TGC TGG GTT TCT GTT GAA ACC AAA CCG GGT GGT GAC TTC GAC TTC

           C   L   L   R   P   P   G   Q   Q   M   E   N   G   V   S   P   E   G   L   L
    Query  TGT TTG CTA AGG CCA CCA GGT CAA CAG ATG GAA AAT GGG GTT TCA CCA GAA GGC CTG CTG
           ||# #|| ||# *|* ||# ||# ||| ||# ||| ||| ||| ||# ||* ||| ||* ||# ||| ||# ||| |||
Optimized  TGC CTG CTG CGT CCG CCG GGT CAG CAG ATG GAA AAC GGT GTT TCT CCG GAA GGT CTG CTG

           P   R   R   L   G   Y   S   R   G   N   R   V   G   G   L   V   V   G   M   I
    Query  CCC AGG CGT TTG GGT TAC TCA CGT GGT AAT CGT GTT GGT GGT CTG GTG GTT GGC ATG ATT
           ||* *|* ||| #|| ||| ||| ||* ||| ||| ||# ||| ||| ||| ||| ||| ||* ||| ||# ||| ||#
Optimized  CCG CGT CGT CTG GGT TAC TCT CGT GGT AAC CGT GTT GGT GGT CTG GTT GTT GGT ATG ATC

           L   V   A   E   H   K   Q   V   N   R   H   F   N   S   N   S   V   T   F   G
    Query  TTG GTG GCT GAA CAC AAG CAG GTC AAC AGG CAT TTT AAC TCC AAT TCT GTC ACA TTT GGT
           #|| ||* ||* ||| ||| ||# ||| ||# ||| *|* ||# ||# ||| ||# ||# ||| ||# ||* ||# |||
Optimized  CTG GTT GCG GAA CAC AAA CAG GTT AAC CGT CAC TTC AAC TCT AAC TCT GTT ACC TTC GGT

           W   S   T   A   P   V   N   P   M   A   A   E   I   V   T   N   Q   A   H   S
    Query  TGG TCA ACC GCA CCT GTT AAC CCA ATG GCA GCA GAG ATC GTG ACC AAC CAG GCA CAT TCG
           ||| ||* ||| ||# ||* ||| ||| ||# ||| ||# ||# ||# ||| ||* ||| ||| ||| ||# ||# ||*
Optimized  TGG TCT ACC GCG CCG GTT AAC CCG ATG GCG GCG GAA ATC GTT ACC AAC CAG GCG CAC TCT

           T   S   R   H   A   W   L   S   I   G   A   Q   N   K   G   P   L   F   P   G
    Query  ACA TCA CGT CAT GCT TGG CTC TCC ATT GGT GCT CAG AAT AAG GGT CCA TTG TTC CCC GGG
           ||* ||* ||| ||# ||* ||| ||* ||# ||# ||| ||* ||| ||# ||# ||| ||# #|| ||| ||* ||*
Optimized  ACC TCT CGT CAC GCG TGG CTG TCT ATC GGT GCG CAG AAC AAA GGT CCG CTG TTC CCG GGT
```

```
           I   P   N   H   F   P   D   S   C   A   S   T   V   V   G   A   M   D   T   S
Query      ATA CCC AAC CAT TTC CCC GAC TCG TGT GCA TCA ACA GTC GTG GGG GCC ATG GAC ACC TCT
           ||* ||* ||| ||# ||| ||* ||| ||* ||# ||# ||* ||* ||# ||* ||* ||* ||| ||| ||| |||
Optimized  ATC CCG AAC CAC TTC CCG GAC TCT TGC GCG TCT ACC GTT GTT GGT GCG ATG GAC ACC TCT

           L   G   G   R   P   S   T   G   V   C   G   P   A   I   S   F   Q   N   N   G
Query      CTT GGC GGC CGG CCG TCC ACT GGG GTG TGT GGG CCT GCC ATA AGC TTC CAA AAC AAT GGT
           ||* ||# ||# ||* ||| ||# ||# ||* ||* ||# ||* ||* ||* ||* **# ||| ||# ||| ||# |||
Optimized  CTG GGT GGT CGT CCG TCT ACC GGT GTT TGC GGT CCG GCG ATC TCT TTC CAG AAC AAC GGT

           D   V   Y   E   N   D   T   P   S   V   M   F   A   T   Y   D   P   L   T   S
Query      GAC GTT TAC GAA AAT GAC ACA CCC TCA GTC ATG TTT GCC ACC TAC GAC CCT CTC ACG TCC
           ||| ||| ||| ||| ||# ||| ||* ||* ||* ||# ||| ||# ||* ||| ||| ||| ||* ||* ||* ||#
Optimized  GAC GTT TAC GAA AAC GAC ACC CCG TCT GTT ATG TTC GCG ACC TAC GAC CCG CTG ACC TCT

           G   T   G   V   A   L   T   N   S   I   N   P   A   S   L   A   L   V   R   I
Query      GGC ACG GGG GTT GCT CTT ACC AAT AGC ATC AAC CCT GCA TCT CTG GCA TTG GTG CGC ATA
           ||# ||* ||* ||| ||* ||* ||| ||# **# ||| ||| ||* ||# ||| ||| ||# #|| ||* ||# ||*
Optimized  GGT ACC GGT GTT GCG CTG ACC AAC TCT ATC AAC CCG GCG TCT CTG GCG CTG GTT CGT ATC

           S   N   N   D   F   D   T   S   G   F   A   N   D   K   N   V   V   V   Q   M
Query      TCC AAC AAC GAC TTT GAC ACC AGT GGT TTT GCC AAC GAT AAA AAT GTG GTG GTG CAA ATG
           ||# ||| ||| ||| ||# ||| ||| **| ||| ||# ||* ||| ||# ||| ||# ||* ||* ||* ||# |||
Optimized  TCT AAC AAC GAC TTC GAC ACC TCT GGT TTC GCG AAC GAC AAA AAC GTT GTT GTT CAG ATG

           S   W   E   M   Y   T   G   T   N   Q   I   R   G   Q   V   T   P   M   S   G
Query      TCA TGG GAG ATG TAC ACA GGC ACC AAC CAA ATT AGA GGT CAA GTC ACA CCA ATG TCT GGT
           ||* ||| ||# ||| ||| ||* ||# ||| ||| ||# ||# *|* ||| ||# ||# ||* ||# ||| ||| |||

           T   N   Y   T   F   T   S   T   G   A   N   T   L   V   L   W   Q   E   R   M
Query      ACC AAC TAC ACC TTT ACT TCC ACA GGT GCG AAC ACT CTT GTG TTG TGG CAG GAG CGT ATG
           ||| ||| ||| ||| ||# ||# ||# ||* ||| ||| ||| ||# ||* ||* #|| ||| ||| ||# ||| |||
Optimized  ACC AAC TAC ACC TTC ACC TCT ACC GGT GCG AAC ACC CTG GTT CTG TGG CAG GAA CGT ATG

           L   S   Y   D   G   H   Q   A   I   L   Y   S   S   Q   L   E   R   T   A   E
Query      CTC AGC TAC GAT GGA CAC CAA GCA ATA TTG TAC TCG TCA CAA TTG GAG AGG ACG GCT GAG
           ||* **# ||| ||# ||* ||| ||# ||# ||* #|| ||| ||* ||* ||# #|| ||# *|* ||* ||* ||#
Optimized  CTG TCT TAC GAC GGT CAC CAG GCG ATC CTG TAC TCT TCT CAG CTG GAA CGT ACC GCG GAA

           Y   F   Q   N   D   I   V   N   I   P   E   N   S   M   A   V   F   N   V   E
Query      TAC TTC CAG AAT GAC ATA GTT AAC ATA CCT GAA AAT TCC ATG GCA GTG TTC AAT GTG GAA
           ||| ||| ||| ||# ||| ||* ||| ||| ||* ||* ||| ||# ||# ||| ||# ||* ||| ||# ||* |||
Optimized  TAC TTC CAG AAC GAC ATC GTT AAC ATC CCG GAA AAC TCT ATG GCG GTT TTC AAC GTT GAA
```

```
           T   N   S   A   S   F   Q   I   G   I   R   P   D   G   Y   M   V   T   G   G
    Query  ACC AAT TCT GCA TCT TTT CAA ATT GGC ATT AGA CCT GAT GGT TAT ATG GTG ACA GGT GGA
           ||| ||# ||| ||# ||| ||# ||# ||# ||# ||# *|* ||* ||# ||| ||# ||| ||* ||* ||| ||*
Optimized  ACC AAC TCT GCG TCT TTC CAG ATC GGT ATC CGT CCG GAC GGT TAC ATG GTT ACC GGT GGT

           S   I   G   V   N   V   P   L   E   P   E   T   R   F   Q   Y   V   G   I   L
    Query  TCT ATT GGC GTC AAT GTG CCG CTT GAA CCT GAA ACT AGA TTC CAA TAC GTT GGT ATC CTC
           ||| ||# ||# ||# ||# ||* ||| ||* ||| ||* ||| ||# *|* ||| ||# ||| ||| ||| ||| ||*
Optimized  TCT ATC GGT GTT AAC GTT CCG CTG GAA CCG GAA ACC CGT TTC CAG TAC GTT GGT ATC CTG

           P   L   S   A   A   L   S   G   P   S   G   N   M   G   R   A   R   R   V   F
    Query  CCT TTG TCT GCA GCT TTA TCT GGG CCC AGT GGG AAC ATG GGA AGG GCC AGA CGG GTG TTC
           ||* #|| ||| ||# ||* #|# ||| ||* ||* **| ||* ||| ||| ||* *|* ||* *|* ||* ||* |||
Optimized  CCG CTG TCT GCG GCG CTG TCT GGT CCG TCT GGT AAC ATG GGT CGT GCG CGT CGT GTT TTC

           Q   .
    Query  CAA TGA
           ||# |||
Optimized  CAG TGA
```

**Dresden (AY694184.2)** – 1,686 nucleotides; 1,296 unchanged, 207 transition and 183 transversion changes.

```
           M   E   G   N   G   S   N   P   E   P   K   Q   S   N   N   P   M   V   V   D
    Query  ATG GAG GGC AAT GGC TCC AAC CCA GAG CCA AAG CAG AGC AAC AAC CCG ATG GTC GTT GAC
           ||| ||# ||# ||# ||# ||# ||| ||# ||# ||# ||# ||| **# ||| ||| ||| ||| ||# ||| |||
Optimized  ATG GAA GGT AAC GGT TCT AAC CCG GAA CCG AAA CAG TCT AAC AAC CCG ATG GTT GTT GAC

           P   P   G   T   T   G   P   T   T   S   H   V   V   V   A   N   P   E   Q   P
    Query  CCG CCT GGC ACA ACA GGT CCG ACC ACA TCC CAC GTT GTT GTT GCT AAT CCG GAG CAA CCC
           ||| ||* ||# ||* ||* ||| ||| ||| ||* ||# ||| ||| ||| ||| ||* ||# ||| ||# ||# ||*
Optimized  CCG CCG GGT ACC ACC GGT CCG ACC ACC TCT CAC GTT GTT GTT GCG AAC CCG GAA CAG CCG

           N   G   A   A   Q   R   L   E   L   A   V   A   T   G   A   I   Q   S   N   V
    Query  AAT GGG GCC GCA CAG CGC CTG GAG TTG GCT GTT GCC ACT GGT GCA ATC CAA TCC AAT GTC
           ||# ||* ||* ||# ||| ||# ||| ||# #|| ||* ||| ||* ||# ||| ||# ||| ||# ||# ||# ||#
Optimized  AAC GGT GCG GCG CAG CGT CTG GAA CTG GCG GTT GCG ACC GGT GCG ATC CAG TCT AAC GTT

           P   E   A   I   R   N   C   F   A   V   F   R   T   F   A   W   N   D   R   M
    Query  CCT GAG GCA ATA CGC AAC TGC TTT GCA GTC TTT CGT ACT TTT GCT TGG AAC GAC AGG ATG
           ||* ||# ||# ||* ||# ||| ||| ||# ||# ||# ||# ||| ||# ||# ||* ||| ||| ||| *|* |||
Optimized  CCG GAA GCG ATC CGT AAC TGC TTC GCG GTT TTC CGT ACC TTC GCG TGG AAC GAC CGT ATG

           P   T   G   T   F   L   G   S   I   S   L   H   P   N   I   N   P   Y   T   A
    Query  CCC ACG GGA ACT TTT CTT GGA TCT ATA TCG CTT CAT CCC AAC ATT AAC CCG TAC ACT GCT
           ||* ||* ||* ||# ||# ||* ||* ||| ||* ||* ||* ||# ||* ||| ||# ||| ||| ||| ||# ||*
Optimized  CCG ACC GGT ACC TTC CTG GGT TCT ATC TCT CTG CAC CCG AAC ATC AAC CCG TAC ACC GCG

           H   L   S   G   M   W   A   G   W   G   G   S   F   E   V   R   L   S   I   S
    Query  CAC CTC TCT GGG ATG TGG GCC GGG TGG GGC GGT AGT TTT GAG GTC CGG CTA TCG ATC TCT
           ||| ||* ||| ||* ||| ||| ||* ||* ||| ||# ||| **| ||# ||# ||# ||* ||# ||* ||| |||
Optimized  CAC CTG TCT GGT ATG TGG GCG GGT TGG GGT GGT TCT TTC GAA GTT CGT CTG TCT ATC TCT

           G   S   G   V   F   A   G   R   I   I   A   S   V   I   P   P   G   V   D   P
    Query  GGT TCT GGC GTG TTC GCT GGG CGT ATC ATT GCT TCT GTC ATA CCA CCA GGG GTT GAT CCC
           ||| ||| ||# ||* ||| ||* ||* ||| ||| ||# ||* ||| ||# ||* ||# ||# ||* ||| ||# ||*
Optimized  GGT TCT GGT GTT TTC GCG GGT CGT ATC ATC GCG TCT GTT ATC CCG CCG GGT GTT GAC CCG
```

```
               S   S   I   R   D   P   G   V   L   P   H   A   F   V   D   A   R   I   T   E
    Query      TCG TCC ATC AGG GAC CCA GGC GTG TTG CCT CAC GCT TTC GTT GAT GCT CGC ATC ACT GAG
               ||* ||# ||| *|* ||| ||# ||# ||* #|| ||* ||| ||* ||| ||| ||# ||* ||# ||| ||# ||#
Optimized      TCT TCT ATC CGT GAC CCG GGT GTT CTG CCG CAC GCG TTC GTT GAC GCG CGT ATC ACC GAA

               P   V   S   F   M   I   P   D   V   R   A   V   D   Y   H   R   M   D   G   A
    Query      CCA GTT TCT TTC ATG ATC CCT GAT GTT AGA GCA GTG GAC TAC CAT AGG ATG GAC GGT GCT
               ||# ||| ||| ||| ||| ||| ||* ||# ||| *|* ||# ||* ||| ||| ||# *|* ||| ||| ||| ||*
Optimized      CCG GTT TCT TTC ATG ATC CCG GAC GTT CGT GCG GTT GAC TAC CAC CGT ATG GAC GGT GCG

               E   P   T   C   S   L   G   F   W   V   Y   Q   P   L   L   N   P   F   S   T
    Query      GAG CCC ACT TGC TCA TTA GGA TTT TGG GTT TAC CAA CCA TTG CTC AAC CCG TTT TCC ACC
               ||# ||* ||# ||| ||* #|# ||* ||# ||| ||| ||| ||# ||# #|| ||* ||| ||| ||# ||# |||
Optimized      GAA CCG ACC TGC TCT CTG GGT TTC TGG GTT TAC CAG CCG CTG CTG AAC CCG TTC TCT ACC

               T   A   V   S   T   C   W   V   S   V   E   T   K   P   G   G   D   F   D   F
    Query      ACC GCC GTT TCA ACA TGT TGG GTG TCA GTA GAG ACT AAA CCA GGC GGT GAT TTT GAT TTC
               ||| ||* ||| ||* ||* ||# ||| ||* ||* ||* ||# ||# ||| ||# ||# ||| ||# ||# ||# |||
Optimized      ACC GCG GTT TCT ACC TGC TGG GTT TCT GTT GAA ACC AAA CCG GGT GGT GAC TTC GAC TTC

               C   L   L   R   P   P   G   Q   Q   M   E   N   G   V   S   P   E   G   L   L
    Query      TGC CTA CTA AGG CCA CCA GGT CAA CAG ATG GAG AAT GGG GTT TCA CCA GAA GGT CTG CTG
               ||| ||# ||# *|* ||# ||# ||| ||# ||| ||| ||# ||# ||* ||| ||* ||# ||| ||| ||| |||
Optimized      TGC CTG CTG CGT CCG CCG GGT CAG CAG ATG GAA AAC GGT GTT TCT CCG GAA GGT CTG CTG

               P   R   R   L   G   Y   S   R   G   N   R   V   G   G   L   V   V   G   M   V
    Query      CCC AGG CGC CTA GGT TAC TCG CGT GGT AAC CGT GTT GGT GGT TTG GTG GTT GGC ATG GTC
               ||* *|* ||# ||# ||| ||| ||* ||| ||| ||| ||| ||| ||| ||| #|| ||* ||| ||# ||| ||#
Optimized      CCG CGT CGT CTG GGT TAC TCT CGT GGT AAC CGT GTT GGT GGT CTG GTT GTT GGT ATG GTT

               L   V   A   E   H   K   Q   V   N   R   H   F   N   S   N   S   V   T   F   G
    Query      TTG GTG GCT GAG CAC AAA CAG GTT AAT AGA CAC TTC AAC TCC AAT TCA GTC ACT TTT GGT
               #|| ||* ||* ||# ||| ||| ||| ||| ||# *|* ||| ||| ||| ||# ||# ||* ||# ||# ||# |||
Optimized      CTG GTT GCG GAA CAC AAA CAG GTT AAC CGT CAC TTC AAC TCT AAC TCT GTT ACC TTC GGT

               W   S   T   A   P   V   N   P   M   A   A   E   I   V   T   N   Q   A   H   S
    Query      TGG TCC ACG GCA CCC GTC AAC CCA ATG GCA GCA GAA ATT GTG ACA AAC CAG GCA CAT TCA
               ||| ||# ||* ||# ||* ||# ||| ||# ||| ||# ||# ||| ||# ||* ||* ||| ||| ||# ||# ||*
Optimized      TGG TCT ACC GCG CCG GTT AAC CCG ATG GCG GCG GAA ATC GTT ACC AAC CAG GCG CAC TCT

               T   S   R   H   A   W   L   S   I   G   A   Q   N   K   G   P   L   F   P   G
    Query      ACG TCA CGT CAT GCT TGG CTT TCC ATT GGT GCT CAA AAC AAG GGT CCA TTG TTT CCT GGA
               ||* ||* ||| ||# ||* ||| ||* ||# ||# ||| ||* ||# ||| ||# ||| ||# #|| ||# ||* ||*
Optimized      ACC TCT CGT CAC GCG TGG CTG TCT ATC GGT GCG CAG AAC AAA GGT CCG CTG TTC CCG GGT
```

```
           I   P   N   H   F   P   D   S   C   A   S   T   I   V   G   A   M   D   T   S
Query      ATA CCC AAC CAT TTC CCC GAC TCA TGT GCA TCA ACA ATC GTG GGG GCC ATG GAC ACC TCC
           ||* ||* ||| ||# ||| ||* ||| ||* ||# ||# ||* ||* ||| ||* ||* ||* ||| ||| ||| ||#
Optimized  ATC CCG AAC CAC TTC CCG GAC TCT TGC GCG TCT ACC ATC GTT GGT GCG ATG GAC ACC TCT

           L   G   G   R   P   S   T   G   V   C   G   P   A   I   S   F   Q   N   N   G
Query      CTT GGT GGC CGG CCG TCT ACT GGG GTA TGT GGA CCT GCC ATA AGC TTT CAA AAT AAT GGT
           ||* ||| ||# ||* ||| ||| ||# ||* ||* ||# ||* ||* ||* ||* **# ||# ||# ||# ||# |||
Optimized  CTG GGT GGT CGT CCG TCT ACC GGT GTT TGC GGT CCG GCG ATC TCT TTC CAG AAC AAC GGT

           D   V   Y   E   N   D   T   P   S   V   M   F   A   T   Y   D   P   L   T   S
Query      GAT GTG TAC GAG AAT GAT ACA CCC TCA GTC ATG TTT GCC ACT TAT GAC CCA CTT ACG TCA
           ||# ||* ||| ||# ||# ||# ||* ||* ||* ||# ||| ||# ||* ||# ||# ||| ||# ||* ||* ||*
Optimized  GAC GTT TAC GAA AAC GAC ACC CCG TCT GTT ATG TTC GCG ACC TAC GAC CCG CTG ACC TCT

           G   T   G   V   A   L   T   N   S   I   N   P   A   S   L   A   L   V   R   I
Query      GGC ACG GGG GTT GCT CTT ACC AAC AGT ATC AAC CCC GCA TCC CTG GCA TTG GTG CGC ATA
           ||# ||* ||* ||| ||* ||* ||| ||| **| ||| ||| ||* ||# ||# ||| ||# #|| ||* ||# ||*
Optimized  GGT ACC GGT GTT GCG CTG ACC AAC TCT ATC AAC CCG GCG TCT CTG GCG CTG GTT CGT ATC

           S   N   N   D   F   D   T   S   G   F   A   N   D   K   N   V   V   V   Q   M
Query      TCC AAT AAT GAC TTT GAC ACC AGT GGC TTT GCC AAT GAC AAG AAC GTG GTG GTT CAA ATG
           ||# ||# ||# ||| ||# ||| ||| **| ||# ||# ||* ||# ||| ||# ||| ||* ||* ||| ||# |||
Optimized  TCT AAC AAC GAC TTC GAC ACC TCT GGT TTC GCG AAC GAC AAA AAC GTT GTT GTT CAG ATG

           S   W   E   M   Y   T   G   T   N   Q   I   R   G   Q   V   T   P   M   S   G
Query      TCA TGG GAG ATG TAT ACA GGC ACC AAT CAA ATT AGA GGT CAG GTC ACA CCA ATG TCT GGC
           ||* ||| ||# ||| ||# ||* ||# ||| ||# ||# ||# *|* ||| ||| ||# ||* ||# ||| ||| ||#

           T   N   Y   T   F   T   S   T   G   A   N   T   L   V   L   W   Q   E   R   M
Query      ACT AAC TAT ACC TTC ACA TCA ACC GGT GCA AAC ACT CTT GTG TTG TGG CAG GAG CGC ATG
           ||# ||| ||# ||| ||| ||* ||* ||| ||| ||# ||| ||# ||* ||* #|| ||| ||| ||# ||# |||
Optimized  ACC AAC TAC ACC TTC ACC TCT ACC GGT GCG AAC ACC CTG GTT CTG TGG CAG GAA CGT ATG

           L   S   Y   D   G   H   Q   A   I   L   Y   S   S   Q   L   E   R   T   A   E
Query      CTC AGC TAT GAT GGA CAC CAA GCA ATA CTG TAC TCG TCA CAA TTG GAG AGG ACG GCT GAG
           ||* **# ||# ||# ||* ||| ||# ||# ||* ||| ||| ||* ||* ||# #|| ||# *|* ||* ||* ||#
Optimized  CTG TCT TAC GAC GGT CAC CAG GCG ATC CTG TAC TCT TCT CAG CTG GAA CGT ACC GCG GAA

           Y   F   Q   N   D   I   V   N   I   P   E   N   S   M   A   V   F   N   V   E
Query      TAC TTT CAG AAC GAC ATA GTT AAC ATA CCT GAA AAC TCT ATG GCA GTG TTC AAT GTG GAA
           ||| ||# ||| ||| ||| ||* ||| ||| ||* ||* ||| ||| ||| ||| ||# ||* ||| ||# ||* |||
Optimized  TAC TTC CAG AAC GAC ATC GTT AAC ATC CCG GAA AAC TCT ATG GCG GTT TTC AAC GTT GAA
```

```
           T   N   S   A   S   F   Q   I   G   I   R   P   D   G   Y   M   V   T   G   G
     Query ACC AAC TCT GCA TCT TTC CAA ATT GGC ATT AGA CCT GAT GGC TAT ATG GTA ACA GGT GGC
           ||| ||| ||| ||# ||| ||| ||# ||# ||# ||# *|* ||* ||# ||# ||# ||| ||* ||* ||| ||#
 Optimized ACC AAC TCT GCG TCT TTC CAG ATC GGT ATC CGT CCG GAC GGT TAC ATG GTT ACC GGT GGT

           S   I   G   I   N   V   P   L   E   P   E   T   R   F   Q   Y   V   G   I   L
     Query TCC ATT GGC ATT AAT GTG CCG CTT GAA CCA GAA ACC CGA TTC CAA TAC GTT GGA ATT CTC
           ||# ||# ||# ||# ||# ||* ||| ||* ||| ||# ||| ||| ||* ||| ||# ||| ||| ||* ||# ||*
 Optimized TCT ATC GGT ATC AAC GTT CCG CTG GAA CCG GAA ACC CGT TTC CAG TAC GTT GGT ATC CTG

           P   L   S   A   A   L   S   G   P   S   G   N   M   G   R   A   R   R   V   F
     Query CCT TTG TCC GCA GCT TTA TCT GGG CCC AGT GGG AAC ATG GGA AGG GCC AGA CGG GTG TTC
           ||* #|| ||# ||# ||* #|# ||| ||* ||* **| ||* ||| ||| ||* *|* ||* *|* ||* ||* |||
 Optimized CCG CTG TCT GCG GCG CTG TCT GGT CCG TCT GGT AAC ATG GGT CGT GCG CGT CGT GTT TTC

           Q   .
     Query CAA TGA
           ||# |||
 Optimized CAG TGA
```

**Sapporo-MT-2010 (HM002617.1)** – 1,686 nucleotides; 1,303 unchanged, 202 transition and 181 transversion changes.

```
              M   E   G   N   G   S   N   S   E   P   K   Q   S   N   N   P   M   V   V   D
    Query    ATG GAG GGC AAT GGC TCC AAC TCA GAG CCA AAG CAG AGC AAC AAC CCA ATG GTC GTT GAC
             ||| ||# ||# ||# ||# ||# ||| ||* ||# ||# ||# ||| **# ||| ||| ||# ||| ||# ||| |||
Optimized    ATG GAA GGT AAC GGT TCT AAC TCT GAA CCG AAA CAG TCT AAC AAC CCG ATG GTT GTT GAC

              P   P   G   T   T   G   P   T   T   S   H   V   V   V   A   N   P   E   Q   P
    Query    CCG CCT GGC ACA ACG GGT CCG ACC ACA TCC CAT GTT GTT GTT GCT AAT CCG GAG CAA CCC
             ||| ||* ||# ||* ||* ||| ||| ||| ||* ||# ||# ||| ||| ||| ||* ||# ||| ||# ||# ||*
Optimized    CCG CCG GGT ACC ACC GGT CCG ACC ACC TCT CAC GTT GTT GTT GCG AAC CCG GAA CAG CCG

              N   G   A   A   Q   R   L   E   L   A   V   A   T   G   A   I   Q   S   N   V
    Query    AAT GGG GCC GCA CAG CGC CTG GAG TTG GCT GTT GCC ACT GGT GCA ATC CAA TCC AAT GTC
             ||# ||* ||* ||# ||| ||# ||| ||# #|| ||* ||| ||* ||# ||| ||# ||| ||# ||# ||# ||#
Optimized    AAC GGT GCG GCG CAG CGT CTG GAA CTG GCG GTT GCG ACC GGT GCG ATC CAG TCT AAC GTT

              P   E   A   I   R   N   C   F   A   V   F   R   T   F   A   W   N   D   R   M
    Query    CCT GAG GCA ATA CGC AAC TGC TTT GCA GTC TTT CGC ACT TTT GCT TGG AAT GAC AGG ATG
             ||* ||# ||# ||* ||# ||| ||| ||# ||# ||# ||# ||# ||# ||# ||* ||| ||# ||| *|* |||
Optimized    CCG GAA GCG ATC CGT AAC TGC TTC GCG GTT TTC CGT ACC TTC GCG TGG AAC GAC CGT ATG

              P   T   G   T   F   L   G   S   I   S   L   H   P   N   I   N   P   Y   T   S
    Query    CCC ACG GGA ACT TTT CTT GGA TCT ATA TCG CTT CAT CCC AAC ATT AAC CCG TAC ACT TCT
             ||* ||* ||* ||# ||# ||* ||* ||| ||* ||* ||* ||# ||* ||| ||# ||| ||| ||| ||# |||
Optimized    CCG ACC GGT ACC TTC CTG GGT TCT ATC TCT CTG CAC CCG AAC ATC AAC CCG TAC ACC TCT

              H   L   S   G   M   W   A   G   W   G   G   S   F   E   V   R   L   S   I   S
    Query    CAC CTC TCT GGG ATG TGG GCC GGG TGG GGC GGT AGC TTT GAG GTC CGG CTG TCG ATC TCT
             ||| ||* ||| ||* ||| ||| ||* ||* ||| ||# ||| **# ||# ||# ||# ||* ||| ||* ||| |||
Optimized    CAC CTG TCT GGT ATG TGG GCG GGT TGG GGT GGT TCT TTC GAA GTT CGT CTG TCT ATC TCT

              G   S   G   V   F   A   G   R   I   I   A   S   V   I   P   P   G   V   D   P
    Query    GGT TCT GGC GTG TTC GCT GGG CGC ATC ATT GCT TCT GTC ATA CCA CCA GGG GTT GAT CCC
             ||| ||| ||# ||* ||| ||* ||* ||# ||| ||# ||* ||| ||# ||* ||# ||# ||* ||| ||# ||*
Optimized    GGT TCT GGT GTT TTC GCG GGT CGT ATC ATC GCG TCT GTT ATC CCG CCG GGT GTT GAC CCG
```

```
           S   S   I   R   D   P   G   V   L   P   H   A   F   V   D   A   R   I   T   E
Query      TCG TCC ATC AGG GAC CCA GGC GTG TTG CCT CAC GCT TTC GTT GAT GCT CGC ATC ACT GAG
           ||* ||# ||| *|* ||| ||# ||# ||* #|| ||* ||| ||* ||| ||| ||# ||* ||# ||| ||# ||#
Optimized  TCT TCT ATC CGT GAC CCG GGT GTT CTG CCG CAC GCG TTC GTT GAC GCG CGT ATC ACC GAA

           P   V   S   F   M   I   P   D   V   R   A   V   D   Y   H   R   M   D   G   A
Query      CCA GTT TCT TTC ATG ATC CCT GAT GTT AGA GCA GTG GAC TAC CAC AGG ATG GAT GGT GCT
           ||# ||| ||| ||| ||| ||| ||* ||# ||| *|* ||# ||* ||| ||| ||| *|* ||| ||# ||| ||*
Optimized  CCG GTT TCT TTC ATG ATC CCG GAC GTT CGT GCG GTT GAC TAC CAC CGT ATG GAC GGT GCG

           E   P   T   C   S   L   G   F   W   V   Y   Q   P   L   L   N   P   F   S   T
Query      GAA CCT ACA TGC TCC TTA GGG TTC TGG GTT TAC CAG CCA TTG CTG AAC CCG TTT TCC ACC
           ||| ||* ||* ||| ||# #|# ||* ||| ||| ||| ||| ||| ||# #|| ||| ||| ||| ||# ||# |||
Optimized  GAA CCG ACC TGC TCT CTG GGT TTC TGG GTT TAC CAG CCG CTG CTG AAC CCG TTC TCT ACC

           T   A   V   S   T   C   W   V   S   V   E   T   K   P   G   G   D   F   D   F
Query      ACC GCC GTT TCA ACA TGT TGG GTA TCA GTG GAA ACT AAA CCA GGC GGC GAC TTT GAC TTC
           ||| ||* ||| ||* ||* ||# ||| ||* ||* ||* ||| ||# ||| ||# ||# ||# ||| ||# ||| |||
Optimized  ACC GCG GTT TCT ACC TGC TGG GTT TCT GTT GAA ACC AAA CCG GGT GGT GAC TTC GAC TTC

           C   L   L   R   P   P   G   Q   Q   M   E   N   G   V   S   P   E   G   L   L
Query      TGT CTG CTA AGG CCA CCA GGT CAA CAA ATG GAA AAT GGG GTT TCA CCA GAA GGC CTG TTG
           ||# ||| ||# *|* ||# ||# ||| ||# ||# ||| ||| ||# ||* ||| ||* ||# ||| ||# ||| #||
Optimized  TGC CTG CTG CGT CCG CCG GGT CAG CAG ATG GAA AAC GGT GTT TCT CCG GAA GGT CTG CTG

           P   R   R   L   G   Y   S   R   G   N   R   V   G   G   L   V   V   G   M   I
Query      CCC AGG CGC TTA GGT TAC TCG CGT GGT AAC CGT GTT GGT GGC TTG GTG GTT GGC ATG ATT
           ||* *|* ||# #|# ||| ||| ||* ||| ||| ||| ||| ||| ||| ||# #|| ||* ||| ||# ||| ||#
Optimized  CCG CGT CGT CTG GGT TAC TCT CGT GGT AAC CGT GTT GGT GGT CTG GTT GTT GGT ATG ATC

           L   V   A   E   H   K   Q   V   N   R   H   F   N   S   N   S   V   T   F   G
Query      TTG GTG GCT GAG CAT AAA CAG GTC AAC AGA CAC TTT AAT TCC AAC TCA GTC ACT TTT GGT
           #|| ||* ||* ||# ||# ||| ||| ||# ||| *|* ||| ||# ||# ||# ||| ||* ||# ||# ||# |||
Optimized  CTG GTT GCG GAA CAC AAA CAG GTT AAC CGT CAC TTC AAC TCT AAC TCT GTT ACC TTC GGT

           W   S   T   A   P   V   N   P   M   A   A   E   I   V   T   N   Q   A   H   S
Query      TGG TCA ACG GCA CCT GTC AAC CCA ATG GCA GCA GAA ATC GTG ACA AAC CAG GCA CAC TCA
           ||| ||* ||* ||# ||* ||# ||| ||# ||| ||# ||# ||| ||| ||* ||* ||| ||| ||# ||| ||*
Optimized  TGG TCT ACC GCG CCG GTT AAC CCG ATG GCG GCG GAA ATC GTT ACC AAC CAG GCG CAC TCT

           T   S   R   H   A   W   L   S   I   G   A   Q   N   K   G   P   L   F   P   G
Query      ACA TCA CGC CAC GCT TGG CTC TCC ATT GGT GCT CAA AAT AAG GGT CCA CTG TTT CCT GGA
           ||* ||* ||# ||| ||* ||| ||* ||# ||# ||| ||* ||# ||# ||# ||| ||# ||| ||# ||* ||*
Optimized  ACC TCT CGT CAC GCG TGG CTG TCT ATC GGT GCG CAG AAC AAA GGT CCG CTG TTC CCG GGT
```

```
           I   P   N   H   F   P   D   S   C   A   S   T   V   V   G   A   M   D   T   S
    Query  ATA CCC AAC CAT TTT CCC GAC TCG TGT GCA TCA ACA GTT GTA GGG GCC ATG GAT ACC TCC
           ||* ||* ||| ||# ||# ||* ||| ||* ||# ||# ||* ||* ||| ||* ||* ||* ||| ||# ||| ||#
Optimized  ATC CCG AAC CAC TTC CCG GAC TCT TGC GCG TCT ACC GTT GTT GGT GCG ATG GAC ACC TCT

           L   G   G   R   P   S   T   G   V   C   G   P   A   I   S   F   Q   N   N   G
    Query  CTT GGT GGC CGG CCA TCC ACT GGG GTG TGT GGG CCT GCC ATA AGC TTT CAA AAC AAT GGT
           ||* ||| ||# ||* ||# ||# ||# ||* ||* ||# ||* ||* ||* ||* **# ||# ||# ||| ||# |||
Optimized  CTG GGT GGT CGT CCG TCT ACC GGT GTT TGC GGT CCG GCG ATC TCT TTC CAG AAC AAC GGT

           D   V   Y   E   N   D   T   P   S   V   M   F   A   T   Y   D   P   L   T   S
    Query  GAT GTG TAT GAG AAT GAC ACA CCC TCA GTC ATG TTT GCC ACT TAT GAC CCA CTC ACG TCA
           ||# ||* ||# ||# ||# ||| ||* ||* ||* ||# ||| ||# ||* ||# ||# ||| ||# ||* ||* ||*
Optimized  GAC GTT TAC GAA AAC GAC ACC CCG TCT GTT ATG TTC GCG ACC TAC GAC CCG CTG ACC TCT

           G   T   G   V   A   L   T   N   S   I   N   P   A   S   L   A   L   V   R   I
    Query  GGT ACG GGG GTT GCT CTC ACC AAC AGC ATT AAC CCC GCA TCC CTG GCG TTG GTG CGC ATT
           ||| ||* ||* ||| ||* ||* ||| ||| **# ||# ||| ||* ||# ||# ||| ||| #|| ||* ||# ||#
Optimized  GGT ACC GGT GTT GCG CTG ACC AAC TCT ATC AAC CCG GCG TCT CTG GCG CTG GTT CGT ATC

           S   N   N   D   F   D   T   S   G   F   A   N   D   K   N   V   V   V   Q   M
    Query  TCC AAC AAT GAC TTT GAC ACC AGT GGC TTT GCC AAC GAT AAA AAC GTA GTG GTT CAA ATG
           ||# ||| ||# ||| ||# ||| ||| **| ||# ||# ||* ||| ||# ||| ||| ||* ||* ||| ||# |||
Optimized  TCT AAC AAC GAC TTC GAC ACC TCT GGT TTC GCG AAC GAC AAA AAC GTT GTT GTT CAG ATG

           S   W   E   M   Y   T   G   T   N   Q   I   R   G   Q   V   T   P   M   S   G
    Query  TCG TGG GAG ATG TAC ACA GGT ACC AAC CAA ATT AGG GGT CAG GTC ACA CCA ATG TCT GGT
           ||* ||| ||# ||| ||| ||* ||| ||| ||| ||# ||# *|* ||| ||| ||# ||* ||# ||| ||| |||

           T   N   Y   T   F   T   S   T   G   A   N   T   L   V   L   W   Q   E   R   M
    Query  ACC AAC TAC ACC TTC ACT TCT ACA GGT GCA AAC ACA CTT GTG TTG TGG CAG GAG CGC ATG
           ||| ||| ||| ||| ||| ||# ||| ||* ||| ||# ||| ||* ||* ||* #|| ||| ||| ||# ||# |||
Optimized  ACC AAC TAC ACC TTC ACC TCT ACC GGT GCG AAC ACC CTG GTT CTG TGG CAG GAA CGT ATG

           L   S   Y   D   G   H   Q   A   I   L   Y   S   S   Q   L   E   R   T   A   E
    Query  CTT AGC TAT GAT GGT CAC CAA GCA ATA CTG TAC TCG TCA CAA TTG GAG AGG ACG GCT GAG
           ||* **# ||# ||# ||| ||| ||# ||# ||* ||| ||| ||* ||* ||# #|| ||# *|* ||* ||* ||#
Optimized  CTG TCT TAC GAC GGT CAC CAG GCG ATC CTG TAC TCT TCT CAG CTG GAA CGT ACC GCG GAA

           Y   F   Q   N   D   I   V   N   I   P   E   N   S   M   A   V   F   N   V   E
    Query  TAC TTT CAG AAT GAC ATA GTC AAC ATA CCA GAA AAT TCC ATG GCA GTG TTC AAT GTG GAA
           ||| ||# ||| ||# ||| ||* ||# ||| ||* ||# ||| ||# ||# ||| ||# ||* ||| ||# ||* |||
Optimized  TAC TTC CAG AAC GAC ATC GTT AAC ATC CCG GAA AAC TCT ATG GCG GTT TTC AAC GTT GAA
```

```
           T   N   S   A   A   F   Q   I   G   I   R   P   D   G   Y   M   V   T   G   G
    Query  ACC AAC TCT GCA GCT TTC CAA ATT GGC ATT AGA CCT GAC GGT TAC ATG GTG ACG GGT GGA
           ||| ||| ||| ||# ||* ||| ||# ||# ||# ||# *|* ||* ||| ||| ||| ||| ||* ||* ||| ||*
Optimized  ACC AAC TCT GCG GCG TTC CAG ATC GGT ATC CGT CCG GAC GGT TAC ATG GTT ACC GGT GGT

           S   I   G   V   N   V   P   L   E   P   E   T   R   F   Q   Y   V   G   I   L
    Query  TCC ATT GGT GTT AAT GTG CCG CTT GAA CCT GAA ACC CGA TTC CAA TAC GTT GGA ATC CTC
           ||# ||# ||| ||| ||# ||* ||| ||* ||| ||* ||| ||| ||* ||| ||# ||| ||| ||* ||| ||*
Optimized  TCT ATC GGT GTT AAC GTT CCG CTG GAA CCG GAA ACC CGT TTC CAG TAC GTT GGT ATC CTG

           P   L   S   A   A   L   S   G   P   S   G   N   M   G   R   A   K   R   V   F
    Query  CCT TTG TCC GCA GCT TTA TCT GGG CCC AGT GGG AAC ATG GGA AGG GCC AAA CGG GTG TTC
           ||* #|| ||# ||# ||* #|# ||| ||* ||* **| ||* ||| ||| ||* *|* ||* ||| ||* ||* |||
Optimized  CCG CTG TCT GCG GCG CTG TCT GGT CCG TCT GGT AAC ATG GGT CGT GCG AAA CGT GTT TTC

           Q   .
    Query  CAA TGA
           ||# |||
Optimized  CAG TGA
```

**Chiba/000496 (AJ606693.2)** – 1,686 nucleotides; 1,311 unchanged, 202 transition and 173 transversion changes.

```
           M   E   G   N   A   S   G   S   A   A   K   N   G   G   E   L   T   T   T   D
    Query  ATG GAA GGC AAT GCC TCC GGC TCA GCA GCA AAG AAC GGT GGT GAA CTC ACC ACT ACT GAC
           ||| ||| ||# ||# ||* ||# ||# ||* ||# ||# ||# ||| ||| ||| ||| ||* ||| ||# ||# |||
Optimized  ATG GAA GGT AAC GCG TCT GGT TCT GCG GCG AAA AAC GGT GGT GAA CTG ACC ACC ACC GAC

           P   P   G   T   T   G   P   T   T   S   H   T   V   V   A   N   P   E   Q   P
    Query  CCG CCT GGT ACA ACG GGT CCA ACC ACA TCC CAT ACT GTT GTT GCT AAT CCG GAA CAA CCC
           ||| ||* ||| ||* ||* ||| ||# ||| ||* ||# ||# ||# ||| ||| ||* ||# ||| ||| ||# ||*
Optimized  CCG CCG GGT ACC ACC GGT CCG ACC ACC TCT CAC ACC GTT GTT GCG AAC CCG GAA CAG CCG

           N   G   A   A   Q   R   L   E   M   A   V   A   T   G   A   I   Q   S   N   V
    Query  AAT GGG GCC GCA CAG CGC TTG GAG ATG GCT GTT GCC ACT GGT GCA ATC CAA TCG AAT GTC
           ||# ||* ||* ||# ||| ||# #|| ||# ||| ||* ||| ||* ||# ||| ||# ||| ||# ||* ||# ||#
Optimized  AAC GGT GCG GCG CAG CGT CTG GAA ATG GCG GTT GCG ACC GGT GCG ATC CAG TCT AAC GTT

           P   E   A   I   R   N   C   F   A   V   F   R   T   F   A   W   N   D   R   M
    Query  CCT GAG GCA ATA CGG AAC TGC TTT GCA GTC TTT CGT ACT TTT GCT TGG AAT GAC AGG ATG
           ||* ||# ||# ||* ||* ||| ||| ||# ||# ||# ||# ||| ||# ||# ||* ||| ||# ||| *|* |||
Optimized  CCG GAA GCG ATC CGT AAC TGC TTC GCG GTT TTC CGT ACC TTC GCG TGG AAC GAC CGT ATG

           P   T   G   T   F   L   G   S   V   S   L   H   P   N   I   N   P   Y   T   S
    Query  CCC ACG GGA ACT TTT CTT GGA TCT GTA TCG CTT CAC CCC AAC ATT AAT CCG TAC ACT TCT
           ||* ||* ||* ||# ||# ||* ||* ||| ||* ||* ||* ||| ||* ||| ||# ||# ||| ||| ||# |||
Optimized  CCG ACC GGT ACC TTC CTG GGT TCT GTT TCT CTG CAC CCG AAC ATC AAC CCG TAC ACC TCT

           H   L   S   A   M   W   A   G   W   G   G   G   F   E   V   R   L   S   I   S
    Query  CAT CTC TCA GCA ATG TGG GCA GGA TGG GGG GGT GGT TTT GAG GTC CGG CTA TCG ATC TCT
           ||# ||* ||* ||# ||| ||| ||# ||* ||| ||* ||| ||| ||# ||# ||# ||* ||# ||* ||| |||
Optimized  CAC CTG TCT GCG ATG TGG GCG GGT TGG GGT GGT GGT TTC GAA GTT CGT CTG TCT ATC TCT

           G   S   G   V   F   A   G   R   I   I   S   S   V   I   P   P   G   V   D   P
    Query  GGG TCT GGC GTC TTT GCT GGG CGT ATC ATT TCT TCT GTC ATA CCA CCT GGG GTT GAT CCA
           ||* ||| ||# ||# ||# ||* ||* ||| ||| ||# ||| ||| ||# ||* ||# ||* ||* ||| ||# ||#
Optimized  GGT TCT GGT GTT TTC GCG GGT CGT ATC ATC TCT TCT GTT ATC CCG CCG GGT GTT GAC CCG
```

```
              T   S   I   R   D   P   G   V   L   P   H   A   F   V   D   A   R   I   T   E
    Query     ACG TCC ATC AGG GAC CCA GGC GTG TTG CCT CAC GCT TTC GTT GAT GCT CGC ATC ACT GAG
              ||* ||# ||| *|* ||| ||# ||# ||* #|| ||* ||| ||* ||| ||| ||# ||* ||# ||| ||# ||#
Optimized     ACC TCT ATC CGT GAC CCG GGT GTT CTG CCG CAC GCG TTC GTT GAC GCG CGT ATC ACC GAA

              P   V   S   F   L   I   P   D   V   R   A   V   D   Y   H   R   M   D   G   E
    Query     CCA GTA TCT TTC CTG ATC CCA GAT GTG CGT GCG GTG GAT TAC CAC CGT ATG GAC GGC GAA
              ||# ||* ||| ||| ||| ||| ||# ||# ||* ||| ||| ||* ||# ||| ||| ||| ||| ||| ||# |||
Optimized     CCG GTT TCT TTC CTG ATC CCG GAC GTT CGT GCG GTT GAC TAC CAC CGT ATG GAC GGT GAA

              E   P   T   C   S   L   G   F   W   V   Y   Q   P   L   L   N   P   F   S   T
    Query     GAG CCC ACA TGC TCG CTG GGG TTC TGG GTG TAC CAG CCA CTG CTA AAC CCA TTT TCC ACA
              ||# ||* ||* ||| ||* ||| ||* ||| ||| ||* ||| ||| ||# ||| ||# ||| ||# ||# ||# ||*
Optimized     GAA CCG ACC TGC TCT CTG GGT TTC TGG GTT TAC CAG CCG CTG CTG AAC CCG TTC TCT ACC

              T   A   V   S   T   C   W   V   S   V   E   T   K   P   G   G   D   F   D   F
    Query     ACA GCA GTT TCA ACC TGT TGG GTG TCG GTT GAG ACG AAG CCG GGT GGT GAT TTT GAT TTC
              ||* ||# ||| ||* ||| ||# ||| ||* ||* ||| ||# ||* ||# ||| ||| ||| ||# ||# ||# |||
Optimized     ACC GCG GTT TCT ACC TGC TGG GTT TCT GTT GAA ACC AAA CCG GGT GGT GAC TTC GAC TTC

              C   L   L   R   P   P   G   Q   R   M   D   N   G   I   T   P   E   G   L   L
    Query     TGC TTG CTG CGC CCA CCA GGA CAA CGC ATG GAC AAC GGC ATC ACA CCC GAG GGT CTG TTG
              ||| #|| ||| ||# ||# ||# ||* ||# ||# ||| ||| ||| ||# ||| ||* ||* ||# ||| ||| #||
Optimized     TGC CTG CTG CGT CCG CCG GGT CAG CGT ATG GAC AAC GGT ATC ACC CCG GAA GGT CTG CTG

              P   R   R   L   G   Y   A   R   G   N   R   V   G   G   L   V   V   G   M   V
    Query     CCG CGG CGC TTG GGG TAT GCG CGT GGT AAC CGT GTT GGA GGC CTT GTG GTG GGC ATG GTC
              ||| ||* ||# #|| ||* ||# ||| ||| ||| ||| ||| ||| ||* ||# ||* ||* ||* ||# ||| ||#
Optimized     CCG CGT CGT CTG GGT TAC GCG CGT GGT AAC CGT GTT GGT GGT CTG GTT GTT GGT ATG GTT

              L   V   A   E   H   K   Q   V   N   R   H   F   N   S   D   S   V   T   L   G
    Query     CTT GTT GCA GAG CAC AAG CAA GTC AAC CGG CAT TTC AAT TCA GAT TCT GTC ACC CTT GGA
              ||* ||| ||# ||# ||| ||# ||# ||# ||| ||* ||# ||| ||# ||* ||# ||| ||# ||| ||* ||*
Optimized     CTG GTT GCG GAA CAC AAA CAG GTT AAC CGT CAC TTC AAC TCT GAC TCT GTT ACC CTG GGT

              W   S   T   A   P   V   N   P   M   A   A   E   I   V   T   N   T   P   R   T
    Query     TGG TCT ACC GCA CCA GTC AAC CCA ATG GCA GCA GAG ATA GTC ACA AAC ACA CCT CGC ACC
              ||| ||| ||| ||# ||# ||# ||| ||# ||| ||# ||# ||# ||* ||# ||* ||| ||* ||* ||# |||
Optimized     TGG TCT ACC GCG CCG GTT AAC CCG ATG GCG GCG GAA ATC GTT ACC AAC ACC CCG CGT ACC

              S   A   R   H   A   W   L   S   I   G   A   Q   G   K   G   P   L   F   P   G
    Query     AGT GCA CGA CAC GCT TGG CTA TCC ATT GGT GCG CAA GGA AAG GGA CCA CTT TTT CCC GGT
              **| ||# ||* ||| ||* ||| ||# ||# ||# ||| ||| ||# ||* ||# ||* ||# ||* ||# ||* |||
Optimized     TCT GCG CGT CAC GCG TGG CTG TCT ATC GGT GCG CAG GGT AAA GGT CCG CTG TTC CCG GGT
```

```
               I   P   N   H   F   P   D   S   C   A   S   T   V   V   G   A   M   N   T   S
     Query     ATA CCT AAC CAC TTT CCC GAC TCC TGT GCC TCC ACA GTA GTT GGT GCA ATG AAC ACA TCT
               ||* ||* ||| ||| ||# ||* ||| ||# ||# ||* ||# ||* ||* ||| ||| ||# ||| ||| ||* |||
 Optimized     ATC CCG AAC CAC TTC CCG GAC TCT TGC GCG TCT ACC GTT GTT GGT GCG ATG AAC ACC TCT

               L   G   G   M   P   S   T   G   I   C   G   P   V   I   G   F   Q   D   N   G
     Query     CTT GGT GGT ATG CCC TCA ACA GGA ATT TGT GGC CCT GTA ATT GGT TTC CAA GAC AAC GGA
               ||* ||| ||| ||| ||* ||* ||* ||* ||# ||# ||# ||* ||* ||# ||| ||| ||# ||| ||| ||*
 Optimized     CTG GGT GGT ATG CCG TCT ACC GGT ATC TGC GGT CCG GTT ATC GGT TTC CAG GAC AAC GGT

               D   V   F   E   T   D   T   P   S   A   M   F   A   T   Y   D   P   L   T   S
     Query     GAC GTG TTT GAA ACA GAC ACA CCA TCA GCC ATG TTT GCC ACA TAT GAT CCC CTC ACC TCA
               ||| ||* ||# ||| ||* ||| ||* ||# ||* ||* ||| ||# ||* ||* ||# ||# ||* ||* ||| ||*
 Optimized     GAC GTT TTC GAA ACC GAC ACC CCG TCT GCG ATG TTC GCG ACC TAC GAC CCG CTG ACC TCT

               G   D   T   V   A   L   S   N   T   I   N   P   L   S   L   A   L   V   R   I
     Query     GGT GAT ACG GTT GCA CTG TCC AAT ACA ATC AAC CCA CTC TCT CTC GCT TTG GTT CGC ATA
               ||| ||# ||* ||| ||# ||| ||# ||# ||* ||| ||| ||# ||* ||| ||* ||* #|| ||| ||# ||*
 Optimized     GGT GAC ACC GTT GCG CTG TCT AAC ACC ATC AAC CCG CTG TCT CTG GCG CTG GTT CGT ATC

               S   N   N   D   F   D   A   T   G   F   S   N   N   K   N   V   V   A   Q   M
     Query     TCA AAC AAT GAC TTT GAT GCA ACC GGG TTT TCC AAC AAT AAG AAT GTT GTG GCT CAG ATG
               ||* ||| ||# ||| ||# ||# ||# ||| ||* ||# ||# ||| ||# ||# ||# ||| ||* ||* ||| |||
 Optimized     TCT AAC AAC GAC TTC GAC GCG ACC GGT TTC TCT AAC AAC AAA AAC GTT GTT GCG CAG ATG

               S   W   E   M   Y   S   G   T   N   Q   I   Q   G   N   V   T   P   M   T   G
     Query     TCA TGG GAA ATG TAT TCT GGC ACC AAC CAA ATT CAG GGC AAC GTG ACT CCA ATG ACA GGC
               ||* ||| ||| ||| ||# ||| ||# ||| ||| ||# ||# ||| ||# ||| ||* ||# ||# ||| ||* ||#

               T   N   Y   T   F   S   S   S   G   A   N   T   L   V   L   W   Q   E   R   M
     Query     ACA AAT TAC ACC TTT TCC TCG TCA GGT GCT AAC ACA CTA GTG TTG TGG CAG GAG CGC ATG
               ||* ||# ||| ||| ||# ||# ||* ||* ||| ||* ||| ||* ||# ||* #|| ||| ||| ||# ||# |||
 Optimized     ACC AAC TAC ACC TTC TCT TCT TCT GGT GCG AAC ACC CTG GTT CTG TGG CAG GAA CGT ATG

               L   S   F   N   E   H   Q   A   I   L   Y   S   S   Q   L   E   R   T   A   E
     Query     CTT AGC TTC AAT GAG CAT CAG GCC ATT TTG TAC TCA TCA CAA TTG GAG CGA ACT GCT GAG
               ||* **# ||| ||# ||# ||# ||| ||* ||# #|| ||| ||* ||* ||# #|| ||# ||* ||# ||* ||#
 Optimized     CTG TCT TTC AAC GAA CAC CAG GCG ATC CTG TAC TCT TCT CAG CTG GAA CGT ACC GCG GAA

               Y   F   Q   N   D   V   V   N   I   P   H   N   T   M   A   V   F   N   V   E
     Query     TAC TTC CAA AAT GAT GTG GTT AAC ATA CCC CAT AAC ACA ATG GCA GTT TTC AAC GTG GAA
               ||| ||| ||# ||# ||# ||* ||| ||| ||* ||* ||# ||| ||* ||| ||# ||| ||| ||| ||* |||
 Optimized     TAC TTC CAG AAC GAC GTT GTT AAC ATC CCG CAC AAC ACC ATG GCG GTT TTC AAC GTT GAA
```

```
           T   N   S   A   S   F   Q   V   G   I   R   A   D   G   Y   M   V   T   G   G
    Query  ACA AAT TCG GCT TCA TTT CAG GTT GGC ATC CGT GCT GAT GGC TAC ATG GTT ACT GGA GGT
           ||* ||# ||* ||* ||* ||# ||| ||| ||# ||| ||| ||* ||# ||# ||| ||| ||| ||# ||* |||
Optimized  ACC AAC TCT GCG TCT TTC CAG GTT GGT ATC CGT GCG GAC GGT TAC ATG GTT ACC GGT GGT

           A   V   G   T   I   V   P   L   E   P   E   T   R   F   Q   Y   V   G   I   L
    Query  GCC GTT GGC ACC ATT GTG CCA CTT GAA CCT GAA ACC CGC TTC CAA TAT GTT GGA ATT CTC
           ||* ||| ||# ||| ||# ||* ||# ||* ||| ||* ||| ||| ||# ||| ||# ||# ||| ||* ||# ||*
Optimized  GCG GTT GGT ACC ATC GTT CCG CTG GAA CCG GAA ACC CGT TTC CAG TAC GTT GGT ATC CTG

           P   L   S   A   A   L   S   G   P   S   G   N   L   G   R   A   R   R   V   F
    Query  CCC CTT TCT GCA GCA TTG TCA GGG CCC AGT GGG AAT TTG GGT AGG GCC AGA AGA GTG TTC
           ||* ||* ||| ||# ||# #|| ||* ||* ||* **| ||* ||# #|| ||| *|* ||* *|* *|* ||* |||
Optimized  CCG CTG TCT GCG GCG CTG TCT GGT CCG TCT GGT AAC CTG GGT CGT GCG CGT CGT GTT TTC

           Q   .
    Query  CAA TGA
           ||# |||
Optimized  CAG TGA
```

**Kashiwa1 (LC190463.1)** – 1,680 nucleotides; 1,290 unchanged, 199 transition and 191 transversion changes.

```
           M   E   G   L   G   S   Q   Q   G   Q   R   P   D   E   Q   V   A   N   S   A
    Query  ATG GAG GGA CTT GGC TCC CAG CAA GGC CAG CGG CCT GAT GAG CAG GTT GCT AAC AGT GCT
           ||| ||# ||* ||* ||# ||# ||| ||# ||# ||| ||* ||* ||# ||# ||| ||| ||* ||| **| ||*
Optimized  ATG GAA GGT CTG GGT TCT CAG CAG GGT CAG CGT CCG GAC GAA CAG GTT GCG AAC TCT GCG

           T   Q   A   T   P   Q   D   T   I   S   P   T   S   A   L   L   L   P   T   Q
    Query  ACA CAG GCC ACC CCC CAA GAC ACA ATT AGT CCC ACA AGT GCA CTT TTG TTG CCA ACT CAA
           ||* ||| ||* ||| ||* ||# ||| ||* ||# **| ||* ||* **| ||# ||* #|| #|| ||# ||# ||#
Optimized  ACC CAG GCG ACC CCG CAG GAC ACC ATC TCT CCG ACC TCT GCG CTG CTG CTG CCG ACC CAG

           I   E   V   P   N   A   T   A   Q   R   V   E   L   A   A   A   T   G   A   I
    Query  ATT GAG GTG CCA AAC GCC ACT GCC CAG CGT GTT GAG TTG GCT GCA GCA ACA GGG GCC ATT
           ||# ||# ||* ||# ||| ||* ||# ||* ||| ||| ||| ||# #|| ||* ||# ||# ||* ||* ||* ||#
Optimized  ATC GAA GTT CCG AAC GCG ACC GCG CAG CGT GTT GAA CTG GCG GCG GCG ACC GGT GCG ATC

           T   S   N   V   P   N   C   V   R   E   C   F   A   A   V   T   T   I   P   W
    Query  ACC AGC AAT GTC CCA AAC TGT GTC CGT GAG TGC TTT GCC GCT GTA ACA ACA ATT CCA TGG
           ||| **# ||# ||# ||# ||| ||# ||# ||| ||# ||| ||# ||* ||* ||* ||* ||* ||# ||# |||
Optimized  ACC TCT AAC GTT CCG AAC TGC GTT CGT GAA TGC TTC GCG GCG GTT ACC ACC ATC CCG TGG

           T   T   R   Q   A   A   N   T   F   L   G   A   I   H   L   G   P   R   I   N
    Query  ACT ACA CGG CAG GCC GCC AAC ACA TTC TTG GGT GCC ATC CAC CTT GGT CCA CGT ATC AAC
           ||# ||* ||* ||| ||* ||* ||| ||* ||| #|| ||| ||* ||| ||| ||* ||| ||# ||| ||| |||
Optimized  ACC ACC CGT CAG GCG GCG AAC ACC TTC CTG GGT GCG ATC CAC CTG GGT CCG CGT ATC AAC

           P   Y   T   A   H   L   S   A   M   F   A   G   W   G   G   G   F   Q   V   R
    Query  CCA TAC ACT GCC CAT CTG AGT GCC ATG TTT GCA GGC TGG GGA GGT GGA TTC CAA GTT CGT
           ||# ||| ||# ||* ||# ||| **| ||* ||| ||# ||# ||# ||| ||* ||| ||* ||| ||# ||| |||
Optimized  CCG TAC ACC GCG CAC CTG TCT GCG ATG TTC GCG GGT TGG GGT GGT GGT TTC CAG GTT CGT

           V   T   L   S   G   S   G   L   Y   A   G   R   V   V   T   A   I   L   P   P
    Query  GTC ACA TTA TCT GGA TCT GGG TTG TAT GCC GGT CGT GTG GTC ACT GCC ATC CTG CCA CCT
           ||# ||* #|# ||| ||* ||| ||* #|| ||# ||* ||| ||| ||* ||# ||# ||* ||| ||| ||# ||*
Optimized  GTT ACC CTG TCT GGT TCT GGT CTG TAC GCG GGT CGT GTT GTT ACC GCG ATC CTG CCG CCG
```

```
           G   V   N   P   A   A   V   Q   N   P   G   V   F   P   H   A   F   I   D   A
    Query  GGC GTC AAT CCG GCT GCG GTG CAA AAC CCT GGA GTG TTC CCA CAC GCG TTC ATT GAC GCG
           ||# ||# ||# ||| ||* ||| ||* ||# ||| ||* ||* ||* ||| ||# ||| ||| ||| ||# ||| |||
Optimized  GGT GTT AAC CCG GCG GCG GTT CAG AAC CCG GGT GTT TTC CCG CAC GCG TTC ATC GAC GCG

           R   T   T   A   P   I   L   I   N   L   P   D   I   R   P   V   D   F   H   R
    Query  CGC ACC ACA GCC CCA ATA TTG ATC AAC CTG CCA GAC ATC CGA CCA GTG GAC TTC CAC CGG
           ||# ||| ||* ||* ||# ||* #|| ||| ||| ||| ||# ||| ||| ||* ||# ||* ||| ||| ||| ||*
Optimized  CGT ACC ACC GCG CCG ATC CTG ATC AAC CTG CCG GAC ATC CGT CCG GTT GAC TTC CAC CGT

           V   D   G   D   D   A   T   A   T   V   G   L   W   V   A   Q   P   L   I   N
    Query  GTG GAT GGT GAT GAT GCC ACA GCA ACG GTC GGG CTG TGG GTG GCC CAA CCA TTG ATC AAC
           ||* ||# ||| ||# ||# ||* ||* ||# ||* ||# ||* ||| ||| ||* ||* ||# ||# #|| ||| |||
Optimized  GTT GAC GGT GAC GAC GCG ACC GCG ACC GTT GGT CTG TGG GTT GCG CAG CCG CTG ATC AAC

           P   F   Q   I   G   S   V   S   T   C   W   L   S   F   E   T   R   P   G   P
    Query  CCT TTC CAG ATT GGC TCA GTG TCA ACT TGT TGG CTA AGC TTT GAG ACC AGG CCA GGC CCG
           ||* ||| ||| ||# ||# ||* ||* ||* ||# ||# ||| ||# **# ||# ||# ||| *|* ||# ||# |||
Optimized  CCG TTC CAG ATC GGT TCT GTT TCT ACC TGC TGG CTG TCT TTC GAA ACC CGT CCG GGT CCG

           D   F   D   F   C   L   L   K   A   P   E   Q   E   M   D   N   G   I   S   P
    Query  GAC TTT GAC TTC TGC CTC TTG AAG GCC CCA GAA CAG GAG ATG GAC AAC GGC ATA TCA CCT

           A   A   L   L   P   R   R   L   G   R   S   K   G   N   R   M   G   G   R   V
    Query  GCC GCT CTC CTC CCG CGT CGC CTC GGG CGT TCG AAA GGT AAT CGC ATG GGA GGT AGG GTT
           ||* ||* ||* ||* ||| ||| ||# ||* ||* ||| ||* ||| ||| ||# ||# ||| ||* ||| *|* |||
Optimized  GCG GCG CTG CTG CCG CGT CGT CTG GGT CGT TCT AAA GGT AAC CGT ATG GGT GGT CGT GTT

           V   G   I   V   V   V   A   R   A   E   Q   V   N   H   H   F   S   A   S   S
    Query  GTG GGC ATT GTT GTT GTC GCA CGT GCA GAG CAG GTC AAC CAC CAC TTT TCT GCT AGT TCC
           ||* ||# ||# ||| ||| ||# ||# ||| ||# ||# ||| ||# ||| ||| ||| ||# ||| ||* **| ||#
Optimized  GTT GGT ATC GTT GTT GTT GCG CGT GCG GAA CAG GTT AAC CAC CAC TTC TCT GCG TCT TCT

           T   T   L   G   W   S   T   L   P   L   E   P   M   A   G   D   V   Y   W   H
    Query  ACC ACA CTC GGC TGG TCA ACA CTA CCA CTA GAA CCT ATG GCT GGG GAC GTG TAC TGG CAT
           ||| ||* ||* ||# ||| ||* ||* ||# ||# ||# ||| ||* ||| ||* ||* ||| ||* ||| ||| ||#
Optimized  ACC ACC CTG GGT TGG TCT ACC CTG CCG CTG GAA CCG ATG GCG GGT GAC GTT TAC TGG CAC

           G   T   D   D   S   G   K   I   R   G   L   V   S   A   A   G   K   G   I   I
    Query  GGC ACT GAC GAC TCT GGA AAA ATT CGT GGC CTT GTC AGT GCA GCA GGA AAG GGC ATC ATC
           ||# ||# ||| ||| ||| ||* ||| ||# ||| ||# ||* ||# **| ||# ||# ||* ||# ||# ||| |||
Optimized  GGT ACC GAC GAC TCT GGT AAA ATC CGT GGT CTG GTT TCT GCG GCG GGT AAA GGT ATC ATC
```

```
          F   P   N   I   V   N   H   W   T   D   V   A   L   S   A   K   T   T   G   N
Query     TTC CCC AAC ATC GTC AAC CAT TGG ACC GAT GTT GCC CTG TCT GCA AAG ACG ACG GGC AAC
          ||| ||* ||| ||| ||# ||| ||# ||| ||| ||# ||| ||* ||| ||| ||# ||# ||* ||* ||# |||
Optimized TTC CCG AAC ATC GTT AAC CAC TGG ACC GAC GTT GCG CTG TCT GCG AAA ACC ACC GGT AAC

          T   T   I   P   T   S   D   F   S   A   S   P   G   A   S   G   P   L   V   T
Query     ACA ACT ATT CCA ACC AGT GAT TTC TCC GCT TCA CCC GGA GCC TCT GGT CCT TTG GTG ACG
          ||* ||# ||# ||# ||| **| ||# ||| ||# ||* ||* ||* ||* ||* ||| ||| ||* #|| ||* ||*
Optimized ACC ACC ATC CCG ACC TCT GAC TTC TCT GCG TCT CCG GGT GCG TCT GGT CCG CTG GTT ACC

          F   A   D   N   G   D   V   N   E   Q   T   A   R   N   A   I   F   T   A   A
Query     TTT GCG GAC AAT GGC GAC GTG AAC GAG CAG ACA GCA CGA AAT GCC ATT TTC ACC GCT GCT
          ||# ||| ||| ||# ||# ||| ||* ||| ||# ||| ||* ||# ||* ||# ||* ||# ||| ||| ||* ||*
Optimized TTC GCG GAC AAC GGT GAC GTT AAC GAA CAG ACC GCG CGT AAC GCG ATC TTC ACC GCG GCG

          T   Q   D   F   T   A   L   S   Q   T   F   D   A   A   G   M   W   V   W   L
Query     ACT CAA GAT TTC ACT GCT TTG TCC CAG ACC TTT GAT GCC GCC GGC ATG TGG GTG TGG TTG
          ||# ||# ||# ||| ||# ||* #|| ||# ||| ||| ||# ||# ||* ||* ||# ||| ||| ||* ||| #||
Optimized ACC CAG GAC TTC ACC GCG CTG TCT CAG ACC TTC GAC GCG GCG GGT ATG TGG GTT TGG CTG

          P   W   T   S   Q   K   P   G   Q   S   D   S   N   I   Y   I   N   P   T   W
Query     CCC TGG ACA TCA CAA AAA CCT GGT CAA TCA GAC AGT AAC ATC TAC ATC AAT CCC ACC TGG
          ||* ||| ||* ||* ||# ||| ||* ||| ||# ||* ||| **| ||| ||| ||| ||| ||# ||* ||| |||
Optimized CCG TGG ACC TCT CAG AAA CCG GGT CAG TCT GAC TCT AAC ATC TAC ATC AAC CCG ACC TGG

          I   N   G   N   P   A   H   A   I   Q   E   R   C   T   N   M   I   G   T   N
Query     ATT AAT GGT AAT CCT GCT CAT GCA ATC CAG GAG CGT TGC ACC AAT ATG ATT GGC ACC AAT
          ||# ||# ||| ||# ||* ||* ||# ||# ||| ||| ||# ||| ||| ||| ||# ||| ||# ||# ||| ||#
Optimized ATC AAC GGT AAC CCG GCG CAC GCG ATC CAG GAA CGT TGC ACC AAC ATG ATC GGT ACC AAC

          F   Q   F   G   G   T   G   T   N   N   I   M   L   W   E   E   Q   H   F   T
Query     TTC CAA TTT GGG GGT ACA GGC ACA AAC AAC ATA ATG CTG TGG GAG GAG CAG CAC TTC ACT
          ||| ||# ||# ||* ||| ||* ||# ||* ||| ||| ||* ||| ||| ||| ||# ||# ||| ||| ||| ||#
Optimized TTC CAG TTC GGT GGT ACC GGT ACC AAC AAC ATC ATG CTG TGG GAA GAA CAG CAC TTC ACC

          S   W   P   G   A   A   R   I   Y   C   S   Q   L   E   S   T   A   E   I   F
Query     TCC TGG CCT GGT GCT GCT CGT ATA TAT TGT TCA CAA TTG GAG AGT ACT GCT GAA ATT TTC
          ||# ||| ||* ||| ||* ||* ||| ||* ||# ||# ||* ||# #|| ||# **| ||# ||* ||| ||# |||
Optimized TCT TGG CCG GGT GCG GCG CGT ATC TAC TGC TCT CAG CTG GAA TCT ACC GCG GAA ATC TTC

          Q   N   N   I   V   N   I   P   A   N   Q   M   A   V   F   N   V   E   T   A
Query     CAA AAC AAC ATA GTG AAC ATC CCA GCC AAC CAA ATG GCA GTG TTT AAT GTG GAA ACA GCT
          ||# ||| ||| ||* ||* ||| ||| ||# ||* ||| ||# ||| ||# ||* ||# ||# ||* ||| ||* ||*
Optimized CAG AAC AAC ATC GTT AAC ATC CCG GCG AAC CAG ATG GCG GTT TTC AAC GTT GAA ACC GCG
```

```
           G   N   S   F   Q   I   G   I   L   P   N   G   Y   S   V   T   N   A   T   I
    Query  GGG AAT TCA TTT CAA ATT GGC ATC CTC CCA AAT GGT TAC TCA GTC ACC AAT GCC ACG ATT
           ||* ||# ||* ||# ||# ||# ||# ||| ||* ||# ||# ||| ||| ||* ||# ||| ||# ||* ||* ||#
Optimized  GGT AAC TCT TTC CAG ATC GGT ATC CTG CCG AAC GGT TAC TCT GTT ACC AAC GCG ACC ATC

           G   T   H   Q   L   L   D   Y   E   T   S   F   K   F   V   G   L   F   P   Q
    Query  GGA ACA CAC CAA CTG TTG GAC TAT GAA ACT AGC TTC AAG TTT GTT GGA CTC TTT CCC CAA
           ||* ||* ||| ||# ||| #|| ||| ||# ||| ||# **# ||| ||# ||# ||| ||* ||* ||# ||* ||#
Optimized  GGT ACC CAC CAG CTG CTG GAC TAC GAA ACC TCT TTC AAA TTC GTT GGT CTG TTC CCG CAG

           S   T   S   L   Q   G   P   N   G   N   S   G   R   A   V   R   F   L   E   .
    Query  AGC ACT TCT CTT CAG GGG CCC AAT GGG AAC AGT GGA CGG GCC GTC AGG TTC CTA GAA TAA
           **# ||# ||| ||* ||| ||* ||* ||# ||* ||| **| ||* ||* ||* ||# *|* ||| ||# ||| |||
Optimized  TCT ACC TCT CTG CAG GGT CCG AAC GGT AAC TCT GGT CGT GCG GTT CGT TTC CTG GAA TAA
```

**Mc10 (AY237420.2)** – 1,677 nucleotides; 1,285 unchanged, 200 transition and 192 transversion changes.

```
           M   E   G   L   G   Q   P   Q   P   Q   G   G   E   K   T   S   P   Q   P   V
    Query  ATG GAG GGC CTA GGC CAA CCA CAG CCA CAG GGT GGC GAA AAG ACC AGC CCA CAG CCT GTG
           ||| ||# ||# ||# ||# ||# ||# ||| ||# ||| ||| ||# ||| ||# ||| **# ||# ||| ||* ||*
Optimized  ATG GAA GGT CTG GGT CAG CCG CAG CCG CAG GGT GGT GAA AAA ACC TCT CCG CAG CCG GTT

           T   P   Q   D   T   I   G   P   T   A   A   L   L   L   P   T   Q   I   E   T
    Query  ACA CCA CAG GAC ACC ATT GGC CCT ACA GCG GCC CTC TTA CTT CCA ACT CAG ATT GAA ACA
           ||* ||# ||| ||| ||| ||# ||# ||* ||* ||| ||* ||* #|# ||* ||# ||# ||| ||# ||| ||*
Optimized  ACC CCG CAG GAC ACC ATC GGT CCG ACC GCG GCG CTG CTG CTG CCG ACC CAG ATC GAA ACC

           P   N   A   S   A   Q   R   L   E   L   A   M   A   T   G   A   V   T   S   N
    Query  CCA AAC GCA AGT GCA CAG CGC TTG GAG TTG GCC ATG GCC ACA GGG GCA GTC ACG AGT AAT
           ||# ||| ||# **| ||# ||| ||# #|| ||# #|| ||* ||| ||* ||* ||* ||# ||# ||* **| ||#
Optimized  CCG AAC GCG TCT GCG CAG CGT CTG GAA CTG GCG ATG GCG ACC GGT GCG GTT ACC TCT AAC

           V   P   N   C   I   R   E   C   F   A   S   V   T   T   I   P   W   T   T   R
    Query  GTA CCA AAC TGT ATT CGT GAG TGT TTT GCC TCT GTT ACC ACA ATC CCC TGG ACA ACT CGA
           ||* ||# ||| ||# ||# ||| ||# ||# ||# ||* ||| ||| ||| ||* ||| ||* ||| ||* ||# ||*
Optimized  GTT CCG AAC TGC ATC CGT GAA TGC TTC GCG TCT GTT ACC ACC ATC CCG TGG ACC ACC CGT

           Q   A   A   N   T   F   L   G   A   I   H   L   G   P   R   I   N   P   Y   T
    Query  CAG GCC GCT AAC ACT TTC CTT GGG GCT ATC CAC CTT GGC CCA CGC ATC AAC CCA TAC ACC
           ||| ||* ||* ||| ||# ||| ||* ||* ||* ||| ||| ||* ||# ||# ||# ||| ||| ||# ||| |||
Optimized  CAG GCG GCG AAC ACC TTC CTG GGT GCG ATC CAC CTG GGT CCG CGT ATC AAC CCG TAC ACC

           A   H   L   S   A   M   F   A   G   W   G   G   G   F   Q   V   R   V   T   I
    Query  GCT CAC CTG AGC GCA ATG TTT GCT GGG TGG GGT GGT GGT TTT CAA GTG AGA GTG ACT ATA
           ||* ||| ||| **# ||# ||| ||# ||* ||* ||| ||| ||| ||| ||# ||# ||* *|* ||* ||# ||*
Optimized  GCG CAC CTG TCT GCG ATG TTC GCG GGT TGG GGT GGT GGT TTC CAG GTT CGT GTT ACC ATC

           S   G   S   G   L   F   A   G   R   A   V   T   A   I   L   P   P   G   V   N
    Query  TCT GGT TCT GGC CTG TTT GCT GGT CGG GCG GTA ACT GCC ATC TTG CCA CCC GGA GTG AAC
           ||| ||| ||| ||# ||| ||# ||* ||| ||* ||| ||* ||# ||* ||| #|| ||# ||* ||* ||* |||
Optimized  TCT GGT TCT GGT CTG TTC GCG GGT CGT GCG GTT ACC GCG ATC CTG CCG CCG GGT GTT AAC
```

```
           P   A   S   V   Q   N   P   G   V   F   P   H   A   F   I   D   A   R   T   T
    Query  CCC GCG AGT GTC CAA AAC CCT GGG GTT TTC CCT CAT GCC TTC ATT GAT GCA CGT ACC ACT
           ||* ||| **| ||# ||# ||| ||* ||* ||| ||| ||* ||# ||* ||| ||# ||# ||# ||| ||| ||#
Optimized  CCG GCG TCT GTT CAG AAC CCG GGT GTT TTC CCG CAC GCG TTC ATC GAC GCG CGT ACC ACC

           E   P   I   L   I   N   L   P   D   I   R   P   V   D   F   H   R   V   D   G
    Query  GAG CCA ATC TTG ATT AAT CTG CCA GAC ATT CGT CCT GTT GAC TTC CAC CGT GTA GAC GGA
           ||# ||# ||| #|| ||# ||# ||| ||# ||| ||# ||| ||* ||| ||| ||| ||| ||| ||* ||| ||*
Optimized  GAA CCG ATC CTG ATC AAC CTG CCG GAC ATC CGT CCG GTT GAC TTC CAC CGT GTT GAC GGT

           D   D   A   T   A   S   V   G   L   W   V   A   Q   P   L   I   N   P   F   Q
    Query  GAC GAT GCC ACG GCA TCT GTT GGA CTG TGG GTA GCT CAA CCC CTC ATC AAC CCA TTT CAG
           ||| ||# ||* ||* ||# ||| ||| ||* ||| ||| ||* ||* ||# ||* ||* ||| ||| ||# ||# |||
Optimized  GAC GAC GCG ACC GCG TCT GTT GGT CTG TGG GTT GCG CAG CCG CTG ATC AAC CCG TTC CAG

           T   G   P   V   S   T   C   W   L   S   F   E   T   R   P   G   P   D   F   D
    Query  ACA GGC CCT GTG TCC ACT TGT TGG TTG AGT TTT GAA ACA AGA CCC GGC CCT GAC TTT GAC
           ||* ||# ||* ||* ||# ||# ||# ||| #|| **| ||# ||| ||* *|* ||* ||# ||* ||| ||# |||
Optimized  ACC GGT CCG GTT TCT ACC TGC TGG CTG TCT TTC GAA ACC CGT CCG GGT CCG GAC TTC GAC

           F   C   L   L   K   A   P   E   Q   Q   M   D   N   G   I   S   P   A   S   L
    Query  TTT TGT CTG TTG AAG GCC CCA GAG CAG CAG ATG GAC AAT GGA ATT TCG CCC GCC TCT TTG

           L   P   R   R   L   G   R   S   R   G   N   R   M   G   G   R   I   V   G   L
    Query  TTG CCC CGT AGG CTC GGA CGT TCC CGA GGC AAC AGA ATG GGT GGG CGA ATT GTG GGA CTG
           #|| ||* ||| *|* ||* ||* ||| ||# ||* ||# ||| *|* ||| ||| ||* ||* ||# ||* ||* |||
Optimized  CTG CCG CGT CGT CTG GGT CGT TCT CGT GGT AAC CGT ATG GGT GGT CGT ATC GTT GGT CTG

           V   V   V   A   A   A   E   Q   V   N   H   H   F   D   A   R   S   T   T   L
    Query  GTT GTG GTG GCG GCT GCG GAA CAG GTG AAC CAT CAC TTT GAT GCC CGG TCA ACA ACA TTA
           ||| ||* ||* ||| ||* ||| ||| ||| ||* ||| ||# ||| ||# ||# ||* ||* ||* ||* ||* #|#
Optimized  GTT GTT GTT GCG GCG GCG GAA CAG GTT AAC CAC CAC TTC GAC GCG CGT TCT ACC ACC CTG

           G   W   S   T   L   P   V   E   P   I   A   G   D   I   S   W   Y   G   D   A
    Query  GGG TGG TCC ACA TTG CCT GTT GAA CCT ATT GCA GGG GAT ATA TCC TGG TAT GGT GAT GCT
           ||* ||| ||# ||* #|| ||* ||| ||| ||* ||# ||# ||* ||# ||* ||# ||| ||# ||| ||# ||*
Optimized  GGT TGG TCT ACC CTG CCG GTT GAA CCG ATC GCG GGT GAC ATC TCT TGG TAC GGT GAC GCG

           G   N   K   S   I   R   G   L   V   S   A   Q   G   K   G   I   I   F   P   N
    Query  GGG AAC AAG TCA ATC CGA GGG CTT GTT AGT GCT CAG GGC AAA GGT ATA ATA TTT CCA AAC
           ||* ||| ||# ||* ||| ||* ||* ||* ||| **| ||* ||| ||# ||| ||| ||* ||* ||# ||# |||
Optimized  GGT AAC AAA TCT ATC CGT GGT CTG GTT TCT GCG CAG GGT AAA GGT ATC ATC TTC CCG AAC
```

```
                I   V   N   H   W   T   D   V   A   L   S   S   K   T   S   N   T   T   T   I
Query           ATA GTC AAC CAC TGG ACT GAC GTT GCA CTG TCC TCC AAG ACA TCT AAC ACC ACA ACC ATA
                ||* ||# ||| ||| ||| ||# ||| ||| ||# ||| ||# ||# ||# ||* ||| ||| ||| ||* ||| ||*
Optimized       ATC GTT AAC CAC TGG ACC GAC GTT GCG CTG TCT TCT AAA ACC TCT AAC ACC ACC ACC ATC

                P   T   D   T   S   T   L   G   N   L   P   G   A   S   G   P   L   V   T   F
Query           CCA ACT GAC ACA TCT ACT CTT GGC AAT TTA CCA GGT GCC TCT GGA CCA CTT GTC ACT TTT
                ||# ||# ||| ||* ||| ||# ||* ||# ||# #|# ||# ||| ||* ||| ||* ||# ||* ||# ||# ||#
Optimized       CCG ACC GAC ACC TCT ACC CTG GGT AAC CTG CCG GGT GCG TCT GGT CCG CTG GTT ACC TTC

                A   D   N   G   D   V   N   E   S   S   A   Q   N   A   I   L   T   A   A   N
Query           GCT GAC AAT GGG GAT GTT AAT GAG AGT TCC GCC CAA AAT GCC ATA TTG ACA GCT GCA AAT
                ||* ||| ||# ||* ||# ||| ||# ||# **| ||# ||* ||# ||# ||* ||* #|| ||* ||* ||# ||#
Optimized       GCG GAC AAC GGT GAC GTT AAC GAA TCT TCT GCG CAG AAC GCG ATC CTG ACC GCG GCG AAC

                Q   N   F   T   S   F   S   P   T   F   D   A   A   G   I   W   V   W   M   P
Query           CAG AAC TTC ACA TCA TTC TCT CCA ACT TTT GAT GCG GCA GGG ATA TGG GTG TGG ATG CCT
                ||| ||| ||| ||* ||* ||| ||| ||# ||# ||# ||# ||| ||# ||* ||* ||| ||* ||| ||| ||*
Optimized       CAG AAC TTC ACC TCT TTC TCT CCG ACC TTC GAC GCG GCG GGT ATC TGG GTT TGG ATG CCG

                W   A   T   D   R   P   G   A   S   D   S   N   I   Y   I   S   P   T   W   V
Query           TGG GCC ACG GAT CGT CCA GGT GCG TCA GAC AGC AAC ATC TAC ATT AGC CCC ACC TGG GTA
                ||| ||* ||* ||# ||| ||# ||| ||| ||* ||| **# ||| ||| ||| ||# **# ||* ||| ||| ||*
Optimized       TGG GCG ACC GAC CGT CCG GGT GCG TCT GAC TCT AAC ATC TAC ATC TCT CCG ACC TGG GTT

                N   G   N   P   S   H   P   I   H   E   K   C   T   N   M   I   G   T   N   F
Query           AAT GGC AAT CCC TCC CAC CCA ATC CAT GAA AAA TGC ACT AAC ATG ATT GGC ACA AAC TTT
                ||# ||# ||# ||* ||# ||| ||# ||| ||# ||| ||| ||| ||# ||| ||| ||# ||# ||* ||| ||#
Optimized       AAC GGT AAC CCG TCT CAC CCG ATC CAC GAA AAA TGC ACC AAC ATG ATC GGT ACC AAC TTC

                Q   F   G   G   T   G   T   N   N   I   M   L   W   Q   E   Q   H   F   T   S
Query           CAG TTT GGA GGG ACC GGC ACC AAC AAC ATC ATG TTG TGG CAG GAA CAG CAC TTC ACA TCC
                ||| ||# ||* ||* ||| ||# ||| ||| ||| ||| ||| #|| ||| ||| ||| ||| ||| ||| ||* ||#
Optimized       CAG TTC GGT GGT ACC GGT ACC AAC AAC ATC ATG CTG TGG CAG GAA CAG CAC TTC ACC TCT

                W   P   G   A   A   E   V   Y   C   S   Q   L   E   S   T   A   E   I   F   Q
Query           TGG CCC GGT GCA GCA GAG GTG TAC TGC TCG CAA CTG GAA AGC ACT GCC GAG ATT TTC CAG
                ||| ||* ||| ||# ||# ||# ||* ||| ||| ||* ||# ||| ||| **# ||# ||* ||# ||# ||| |||
Optimized       TGG CCG GGT GCG GCG GAA GTT TAC TGC TCT CAG CTG GAA TCT ACC GCG GAA ATC TTC CAG

                N   N   I   V   N   I   P   M   N   Q   M   A   V   F   N   V   E   T   A   G
Query           AAC AAC ATT GTT AAC ATC CCA ATG AAC CAA ATG GCA GTG TTT AAT GTT GAG ACT GCA GGT
                ||| ||| ||# ||| ||| ||| ||# ||| ||| ||# ||| ||# ||* ||# ||# ||| ||# ||# ||# |||
Optimized       AAC AAC ATC GTT AAC ATC CCG ATG AAC CAG ATG GCG GTT TTC AAC GTT GAA ACC GCG GGT
```

```
           N   S   F   Q   I   A   I   L   P   N   G   Y   C   V   T   N   A   P   V   G
    Query  AAT TCA TTC CAA ATA GCC ATC TTG CCC AAT GGT TAT TGT GTC ACC AAC GCA CCA GTG GGA
           ||# ||* ||| ||# ||* ||* ||| #|| ||* ||# ||| ||# ||# ||# ||| ||| ||# ||# ||* ||*
Optimized  AAC TCT TTC CAG ATC GCG ATC CTG CCG AAC GGT TAC TGC GTT ACC AAC GCG CCG GTT GGT

           T   H   Q   L   L   D   Y   E   T   S   F   K   F   V   G   L   F   P   Q   S
    Query  ACA CAC CAA CTT CTT GAC TAT GAG ACT AGC TTC AAA TTT GTA GGA CTC TTC CCC CAA AGC
           ||* ||| ||# ||* ||* ||| ||# ||# ||# **# ||| ||| ||# ||* ||* ||* ||| ||* ||# **#
Optimized  ACC CAC CAG CTG CTG GAC TAC GAA ACC TCT TTC AAA TTC GTT GGT CTG TTC CCG CAG TCT

           T   S   L   Q   G   P   H   G   N   S   G   R   A   V   R   F   L   E   .
    Query  ACT TCA CTT CAG GGG CCC CAT GGG AAC AGT GGC CGG GCC GTT AGG TTC TTA GAA TAA
           ||# ||* ||* ||| ||* ||* ||# ||* ||| **| ||# ||* ||* ||| *|* ||| #|# ||| |||
Optimized  ACC TCT CTG CAG GGT CCG CAC GGT AAC TCT GGT CGT GCG GTT CGT TTC CTG GAA TAA
```

**C12 (AY603425.1)** – 1,686 nucleotides; 1,285 unchanged, 200 transition and 201 transversion changes.

```
          M   E   G   V   P   R   P   E   G   P   K   A   N   S   N   E   N   V   P   L
    Query ATG GAG GGT GTA CCC CGC CCA GAG GGC CCC AAG GCC AAC TCC AAT GAA AAT GTG CCT CTT
          ||| ||# ||| ||* ||* ||# ||# ||# ||# ||* ||# ||* ||| ||# ||# ||| ||# ||* ||* ||*
Optimized ATG GAA GGT GTT CCG CGT CCG GAA GGT CCG AAA GCG AAC TCT AAC GAA AAC GTT CCG CTG

          A   S   P   Q   D   T   I   G   P   N   A   A   L   L   L   P   T   Q   I   E
    Query GCG TCA CCA CAA GAC ACA ATT GGT CCA AAC GCG GCA CTT CTC CTT CCC ACG CAG ATT GAG
          ||| ||* ||# ||# ||| ||* ||# ||| ||# ||| ||| ||# ||* ||* ||* ||* ||* ||| ||# ||#
Optimized GCG TCT CCG CAG GAC ACC ATC GGT CCG AAC GCG GCG CTG CTG CTG CCG ACC CAG ATC GAA

          T   P   N   G   A   A   Q   R   V   E   M   A   A   A   T   G   A   V   S   N
    Query ACA CCC AAC GGA GCT GCA CAA AGG GTG GAG ATG GCT GCA GCG ACA GGG GCC GTT AGC AAC
          ||* ||* ||| ||* ||* ||# ||# *|* ||* ||# ||| ||* ||# ||| ||* ||* ||* ||| **# |||
Optimized ACC CCG AAC GGT GCG GCG CAG CGT GTT GAA ATG GCG GCG GCG ACC GGT GCG GTT TCT AAC

          N   V   P   M   C   V   R   E   C   F   A   S   V   T   T   L   P   W   T   T
    Query AAC GTC CCC ATG TGT GTG AGG GAG TGC TTT GCT AGT GTG ACC ACG CTA CCT TGG ACA ACA
          ||| ||# ||* ||| ||# ||* *|* ||# ||| ||# ||* **| ||* ||| ||* ||# ||* ||| ||* ||*
Optimized AAC GTT CCG ATG TGC GTT CGT GAA TGC TTC GCG TCT GTT ACC ACC CTG CCG TGG ACC ACC

          R   Q   A   S   N   T   F   L   G   A   I   H   L   G   P   R   I   N   P   Y
    Query CGC CAA GCG TCC AAC ACC TTT CTT GGC GCC ATC CAT CTT GGT CCA CGC ATT AAC CCT TAC
          ||# ||# ||| ||# ||| ||| ||# ||* ||# ||* ||| ||# ||* ||| ||# ||# ||# ||| ||* |||
Optimized CGT CAG GCG TCT AAC ACC TTC CTG GGT GCG ATC CAC CTG GGT CCG CGT ATC AAC CCG TAC

          T   A   H   L   S   A   M   F   A   G   W   G   G   S   F   Q   I   R   V   T
    Query ACT GCA CAT CTG AGT GCC ATG TTT GCT GGT TGG GGC GGC AGT TTC CAA ATA CGG GTC ACC
          ||# ||# ||# ||| **| ||* ||| ||# ||* ||| ||| ||# ||# **| ||| ||# ||* ||* ||# |||
Optimized ACC GCG CAC CTG TCT GCG ATG TTC GCG GGT TGG GGT GGT TCT TTC CAG ATC CGT GTT ACC

          L   S   G   S   G   L   Y   A   G   R   A   V   V   A   V   L   P   P   G   V
    Query CTG TCT GGT TCC GGG CTG TAC GCA GGC CGA GCA GTT GTT GCA GTG CTA CCT CCA GGT GTC
          ||| ||| ||| ||# ||* ||| ||| ||# ||# ||* ||# ||| ||| ||# ||* ||# ||* ||# ||| ||#
Optimized CTG TCT GGT TCT GGT CTG TAC GCG GGT CGT GCG GTT GTT GCG GTT CTG CCG CCG GGT GTT
```

```
           N   P   A   N   V   Q   N   P   G   V   F   P   H   A   F   I   D   A   R   T
    Query  AAT CCA GCC AAT GTC CAA AAC CCT GGA GTT TTT CCA CAT GCT TTC ATT GAT GCC CGC ACC
           ||# ||# ||* ||# ||# ||# ||| ||* ||* ||| ||# ||# ||# ||* ||| ||# ||# ||* ||# |||
Optimized  AAC CCG GCG AAC GTT CAG AAC CCG GGT GTT TTC CCG CAC GCG TTC ATC GAC GCG CGT ACC

           V   D   P   I   L   I   N   L   P   D   I   R   A   V   D   Y   H   R   V   D
    Query  GTG GAC CCC ATC CTC ATC AAC CTC CCC GAC ATC CGT GCA GTG GAC TAT CAC CGC GTG GAT
           ||* ||| ||* ||| ||* ||| ||| ||* ||* ||| ||| ||| ||# ||* ||| ||# ||| ||# ||* ||#
Optimized  GTT GAC CCG ATC CTG ATC AAC CTG CCG GAC ATC CGT GCG GTT GAC TAC CAC CGT GTT GAC

           G   D   E   Q   T   A   T   V   G   L   W   V   A   Q   P   L   I   N   P   F
    Query  GGT GAT GAG CAA ACT GCA ACC GTT GGG CTC TGG GTT GCT CAA CCT CTC ATC AAC CCG TTT
           ||| ||# ||# ||# ||# ||# ||| ||| ||* ||* ||| ||| ||* ||# ||* ||* ||| ||| ||| ||#
Optimized  GGT GAC GAA CAG ACC GCG ACC GTT GGT CTG TGG GTT GCG CAG CCG CTG ATC AAC CCG TTC

           Q   T   G   S   I   S   T   C   W   L   T   F   E   T   R   P   G   P   D   F
    Query  CAA ACT GGA TCC ATC TCC ACG TGT TGG TTG ACA TTT GAG ACT AGA CCT GGA CCT GAT TTT
           ||# ||# ||* ||# ||| ||# ||* ||# ||| #|| ||* ||# ||# ||# *|* ||* ||* ||* ||# ||#
Optimized  CAG ACC GGT TCT ATC TCT ACC TGC TGG CTG ACC TTC GAA ACC CGT CCG GGT CCG GAC TTC

           D   F   C   L   L   K   A   P   E   Q   E   M   D   N   G   I   S   P   A   N
    Query  GAC TTT TGC CTT CTC AAG GCC CCA GAG CAA GAA ATG GAC AAT GGC ATA TCA CCT GCC AAC
           ||| ||# ||| ||* ||* ||# ||* ||# ||# ||# ||| ||| ||| ||# ||# ||* ||* ||* ||* |||
Optimized  GAC TTC TGC CTG CTG AAA GCG CCG GAA CAG GAA ATG GAC AAC GGT ATC TCT CCG GCG AAC

           L   L   P   R   R   L   G   R   S   R   G   N   R   L   G   G   R   V   V   G
    Query  TTA CTC CCT CGT CGC CTC GGG CGA TCT CGA GGC AAC AGG CTT GGT GGC CGG GTG GTG GGT
           #|# ||* ||* ||| ||# ||* ||* ||* ||| ||* ||# ||| *|* ||* ||| ||# ||* ||* ||* |||
Optimized  CTG CTG CCG CGT CGT CTG GGT CGT TCT CGT GGT AAC CGT CTG GGT GGT CGT GTT GTT GGT

           L   V   V   V   A   V   A   E   Q   V   N   H   H   F   A   A   N   S   T   T
    Query  CTC GTG GTG GTG GCC GTT GCG GAA CAA GTT AAC CAC CAC TTT GCC GCT AAT TCC ACC ACG
           ||* ||* ||* ||* ||* ||| ||| ||| ||# ||| ||| ||| ||| ||# ||* ||* ||# ||# ||| ||*
Optimized  CTG GTT GTT GTT GCG GTT GCG GAA CAG GTT AAC CAC CAC TTC GCG GCG AAC TCT ACC ACC

           L   G   W   S   T   L   P   I   E   P   I   A   G   A   I   S   W   Y   Q   N
    Query  CTC GGT TGG TCG ACA CTC CCA ATT GAG CCC ATT GCT GGT GCC ATT TCC TGG TAT CAG AAC
           ||* ||| ||| ||* ||* ||* ||# ||# ||# ||* ||# ||* ||| ||* ||# ||# ||| ||# ||| |||
Optimized  CTG GGT TGG TCT ACC CTG CCG ATC GAA CCG ATC GCG GGT GCG ATC TCT TGG TAC CAG AAC

           T   T   P   G   I   S   T   R   G   L   L   S   A   E   G   K   G   I   I   F
    Query  ACC ACA CCA GGC ATT AGC ACA CGC GGC CTG CTC AGT GCT GAA GGT AAA GGC ATC ATC TTT
           ||| ||* ||# ||# ||# **# ||* ||# ||# ||| ||* **| ||* ||| ||| ||| ||# ||| ||| ||#
Optimized  ACC ACC CCG GGT ATC TCT ACC CGT GGT CTG CTG TCT GCG GAA GGT AAA GGT ATC ATC TTC
```

```
           P   N   I   V   N   H   W   T   D   V   A   L   S   S   K   T   S   G   R   T
    Query  CCA AAC ATA GTC AAC CAC TGG ACT GAC GTT GCC CTC TCC AGC AAA ACT TCC GGG AGA ACC
           ||# ||| ||* ||# ||| ||| ||| ||# ||| ||| ||* ||* ||# **# ||| ||# ||# ||* *|* |||
Optimized  CCG AAC ATC GTT AAC CAC TGG ACC GAC GTT GCG CTG TCT TCT AAA ACC TCT GGT CGT ACC

           T   V   P   T   D   Q   A   N   L   N   Q   C   P   G   A   S   G   P   V   V
    Query  ACT GTA CCA ACT GAC CAG GCC AAC CTT AAC CAG TGT CCT GGG GCC TCT GGT CCC GTC GTG
           ||# ||* ||# ||# ||| ||| ||* ||| ||* ||| ||| ||# ||* ||* ||* ||| ||| ||* ||# ||*
Optimized  ACC GTT CCG ACC GAC CAG GCG AAC CTG AAC CAG TGC CCG GGT GCG TCT GGT CCG GTT GTT

           M   F   Q   N   N   G   D   V   N   E   T   S   A   N   N   C   V   L   T   A
    Query  ATG TTC CAA AAC AAT GGG GAT GTG AAT GAG ACA AGT GCC AAC AAC TGT GTG TTA ACA GCT
           ||| ||| ||# ||| ||# ||* ||# ||* ||# ||# ||* **| ||* ||| ||| ||# ||* #|# ||* ||*
Optimized  ATG TTC CAG AAC AAC GGT GAC GTT AAC GAA ACC TCT GCG AAC AAC TGC GTT CTG ACC GCG

           A   S   H   D   F   V   N   L   S   S   N   F   D   A   A   G   M   W   V   W
    Query  GCC TCA CAT GAT TTT GTC AAC CTG TCT AGT AAT TTT GAT GCG GCT GGC ATG TGG GTT TGG
           ||* ||* ||# ||# ||# ||# ||| ||| ||| **| ||# ||# ||# ||| ||* ||# ||| ||| ||| |||
Optimized  GCG TCT CAC GAC TTC GTT AAC CTG TCT TCT AAC TTC GAC GCG GCG GGT ATG TGG GTT TGG

           L   P   W   T   T   T   K   P   D   A   T   I   N   R   N   V   Y   I   T   P
    Query  CTC CCT TGG ACA ACT ACC AAG CCT GAT GCC ACA ATA AAC CGC AAC GTT TAC ATC ACC CCA
           ||* ||* ||| ||* ||# ||| ||# ||* ||# ||* ||* ||* ||| ||# ||| ||| ||| ||| ||| ||#
Optimized  CTG CCG TGG ACC ACC ACC AAA CCG GAC GCG ACC ATC AAC CGT AAC GTT TAC ATC ACC CCG

           T   W   I   N   G   D   P   S   R   P   I   H   G   K   C   T   N   M   V   G
    Query  ACC TGG ATC AAT GGT GAC CCC TCT CGA CCA ATC CAT GGC AAG TGT ACC AAT ATG GTT GGA
           ||| ||| ||| ||# ||| ||| ||* ||| ||* ||# ||| ||# ||# ||# ||# ||| ||# ||| ||| ||*

           T   N   F   Q   F   G   G   T   G   T   N   N   I   M   L   W   Q   E   Q   H
    Query  ACT AAC TTT CAG TTT GGT GGC ACT GGT ACT AAC AAC ATT ATG CTT TGG CAG GAA CAG CAC
           ||# ||| ||# ||| ||# ||| ||# ||# ||| ||# ||| ||| ||# ||| ||* ||| ||| ||| ||| |||
Optimized  ACC AAC TTC CAG TTC GGT GGT ACC GGT ACC AAC AAC ATC ATG CTG TGG CAG GAA CAG CAC

           F   T   S   F   P   G   A   A   E   V   Y   C   S   Q   L   E   S   T   A   E
    Query  TTC ACA TCT TTT CCT GGA GCA GCA GAA GTC TAC TGT TCA CAG CTT GAG AGC ACT GCT GAG
           ||| ||* ||| ||# ||* ||* ||# ||# ||| ||# ||| ||# ||* ||| ||* ||# **# ||# ||* ||#
Optimized  TTC ACC TCT TTC CCG GGT GCG GCG GAA GTT TAC TGC TCT CAG CTG GAA TCT ACC GCG GAA

           M   F   Q   N   N   V   V   N   I   P   A   N   Q   M   A   V   F   N   V   E
    Query  ATG TTC CAG AAC AAT GTT GTT AAT ATA CCA GCT AAT CAG ATG GCA GTC TTC AAT GTG GAA
           ||| ||| ||| ||| ||# ||| ||| ||# ||* ||# ||* ||# ||| ||| ||# ||# ||| ||# ||* |||
Optimized  ATG TTC CAG AAC AAC GTT GTT AAC ATC CCG GCG AAC CAG ATG GCG GTT TTC AAC GTT GAA
```

```
           T   A   G   N   T   F   Q   I   A   I   M   P   N   G   Y   C   V   T   N   A
    Query  ACT GCG GGC AAC ACA TTT CAA ATT GCT ATC ATG CCA AAT GGT TAT TGT GTT ACC AAT GCA
           ||# ||| ||# ||| ||* ||# ||# ||# ||* ||| ||| ||# ||# ||| ||# ||# ||| ||| ||# ||#
Optimized  ACC GCG GGT AAC ACC TTC CAG ATC GCG ATC ATG CCG AAC GGT TAC TGC GTT ACC AAC GCG

           A   V   G   T   H   Q   L   L   D   Y   E   T   S   F   R   F   V   G   L   F
    Query  GCT GTT GGT ACT CAC CAG CTC CTG GAT TAC GAG ACA AGT TTT AGG TTT GTT GGA CTT TTC
           ||* ||| ||| ||# ||| ||| ||* ||| ||# ||| ||# ||* **| ||# *|* ||# ||| ||* ||* |||
Optimized  GCG GTT GGT ACC CAC CAG CTG CTG GAC TAC GAA ACC TCT TTC CGT TTC GTT GGT CTG TTC

           P   Q   S   T   S   L   Q   G   P   N   G   N   A   G   R   A   V   R   F   L
    Query  CCC CAA AGC ACT AGT CTT CAA GGG CCC AAT GGG AAC GCT GGA CGG GCC GTC AGA TTC CTT
           ||* ||# **# ||# **| ||* ||# ||* ||* ||# ||* ||| ||* ||* ||* ||* ||# *|* ||| ||*
Optimized  CCG CAG TCT ACC TCT CTG CAG GGT CCG AAC GGT AAC GCG GGT CGT GCG GTT CGT TTC CTG

           E   .
    Query  GAA TAA
           ||| |||
Optimized  GAA TAA
```

**NGY-1 (AB775659.2)** – 1,716 nucleotides; 1,323 unchanged, 220 transition and 173 transversion changes.

```
           M   E   G   N   G   S   P   R   G   G   N   Q   P   Q   S   H   N   G   G   T
Query      ATG GAG GGC AAT GGC TCC CCA CGA GGA GGT AAC CAG CCT CAA TCC CAC AAT GGT GGC ACT
           ||| ||# ||# ||# ||# ||# ||# ||* ||* ||| ||| ||| ||* ||# ||# ||| ||# ||| ||# ||#
Optimized  ATG GAA GGT AAC GGT TCT CCG CGT GGT GGT AAC CAG CCG CAG TCT CAC AAC GGT GGT ACC

           S   P   A   Q   A   A   P   P   G   T   T   G   P   A   E   A   P   L   V   P
Query      TCA CCT GCT CAG GCT GCA CCG CCT GGC ACG ACT GGC CCT GCG GAA GCA CCC CTT GTA CCT
           ||* ||* ||* ||| ||* ||# ||| ||* ||# ||* ||# ||# ||* ||| ||| ||# ||* ||* ||* ||*
Optimized  TCT CCG GCG CAG GCG GCG CCG CCG GGT ACC ACC GGT CCG GCG GAA GCG CCG CTG GTT CCG

           V   N   P   E   Q   P   N   S   I   A   Q   R   M   E   L   A   V   A   T   G
Query      GTT AAT CCT GAA CAG CCC AAT TCC ATT GCC CAA CGC ATG GAG CTG GCT GTT GCC ACA GGA
           ||| ||# ||* ||| ||| ||* ||# ||# ||# ||* ||# ||# ||| ||# ||| ||* ||| ||* ||* ||*
Optimized  GTT AAC CCG GAA CAG CCG AAC TCT ATC GCG CAG CGT ATG GAA CTG GCG GTT GCG ACC GGT

           A   T   T   S   N   V   P   E   C   V   R   N   C   F   A   L   L   R   T   I
Query      GCA ACA ACC TCA AAT GTC CCT GAA TGT GTG CGG AAC TGC TTT GCT CTC CTT CGT ACG ATT
           ||# ||* ||| ||* ||# ||# ||* ||| ||# ||* ||* ||| ||| ||# ||* ||* ||* ||| ||* ||#
Optimized  GCG ACC ACC TCT AAC GTT CCG GAA TGC GTT CGT AAC TGC TTC GCG CTG CTG CGT ACC ATC

           P   W   N   S   R   Q   P   Q   G   S   L   L   T   A   V   S   L   H   P   D
Query      CCT TGG AAT TCT CGA CAG CCC CAG GGA TCT CTT CTC ACA GCT GTT TCT TTA CAC CCT GAC
           ||* ||| ||# ||| ||* ||| ||* ||| ||* ||| ||* ||* ||* ||* ||| ||| #|# ||| ||* |||
Optimized  CCG TGG AAC TCT CGT CAG CCG CAG GGT TCT CTG CTG ACC GCG GTT TCT CTG CAC CCG GAC

           I   N   P   Y   T   K   H   L   A   Q   M   F   A   G   W   G   G   A   M   D
Query      ATC AAC CCG TAC ACA AAA CAT CTT GCT CAA ATG TTT GCC GGA TGG GGA GGG GCA ATG GAT
           ||| ||| ||| ||| ||* ||| ||# ||* ||* ||# ||| ||# ||* ||* ||| ||* ||* ||# ||| ||#
Optimized  ATC AAC CCG TAC ACC AAA CAC CTG GCG CAG ATG TTC GCG GGT TGG GGT GGT GCG ATG GAC

           V   R   V   T   I   S   G   S   G   L   F   A   G   K   L   V   C   G   I   L
Query      GTC CGT GTC ACG ATC TCA GGC TCG GGC TTA TTT GCA GGG AAG CTT GTG TGT GGG ATA CTT
           ||# ||| ||# ||* ||| ||* ||# ||* ||# #|# ||# ||# ||* ||# ||* ||* ||# ||* ||* ||*
Optimized  GTT CGT GTT ACC ATC TCT GGT TCT GGT CTG TTC GCG GGT AAA CTG GTT TGC GGT ATC CTG
```

```
          P   P   G   V   N   P   T   L   V   S   D   P   G   V   L   P   H   A   L   V
Query     CCT CCT GGA GTT AAT CCC ACT CTT GTC AGC GAT CCG GGG GTT CTG CCA CAC GCT TTG GTA
          ||* ||* ||* ||| ||# ||* ||# ||* ||# **# ||# ||| ||* ||| ||| ||# ||| ||* #|| ||*
Optimized CCG CCG GGT GTT AAC CCG ACC CTG GTT TCT GAC CCG GGT GTT CTG CCG CAC GCG CTG GTT

          D   A   R   V   T   E   P   A   C   F   N   V   P   D   V   R   A   V   D   Y
Query     GAT GCC CGC GTG ACT GAA CCA GCA TGC TTC AAC GTG CCT GAT GTC CGC GCC GTC GAC TAC
          ||# ||* ||# ||* ||# ||| ||# ||# ||| ||| ||| ||* ||* ||# ||# ||# ||* ||# ||| |||
Optimized GAC GCG CGT GTT ACC GAA CCG GCG TGC TTC AAC GTT CCG GAC GTT CGT GCG GTT GAC TAC

          H   R   T   D   G   D   E   A   T   A   T   L   G   I   W   V   L   Q   P   L
Query     CAT CGC ACT GAC GGC GAT GAG GCA ACT GCA ACA CTT GGC ATT TGG GTG CTC CAA CCA CTA
          ||# ||# ||# ||| ||# ||# ||# ||# ||# ||# ||* ||* ||# ||# ||| ||* ||* ||# ||# ||#
Optimized CAC CGT ACC GAC GGT GAC GAA GCG ACC GCG ACC CTG GGT ATC TGG GTT CTG CAG CCG CTG

          I   N   P   F   S   T   E   A   V   S   T   A   W   I   S   I   E   T   K   P
Query     ATC AAC CCC TTC TCC ACT GAA GCT GTT TCA ACC GCT TGG ATT TCA ATT GAG ACC AAA CCA
          ||| ||| ||* ||| ||# ||# ||| ||* ||| ||* ||| ||* ||| ||# ||* ||# ||# ||| ||| ||#
Optimized ATC AAC CCG TTC TCT ACC GAA GCG GTT TCT ACC GCG TGG ATC TCT ATC GAA ACC AAA CCG

          G   G   D   F   D   L   C   L   M   K   P   P   N   Q   A   M   D   N   G   T
Query     GGT GGT GAT TTT GAT CTT TGC TTG ATG AAA CCT CCT AAC CAA GCC ATG GAT AAC GGC ACA
          ||| ||| ||# ||# ||# ||* ||| #|| ||| ||| ||* ||* ||| ||# ||* ||| ||# ||| ||# ||*
Optimized GGT GGT GAC TTC GAC CTG TGC CTG ATG AAA CCG CCG AAC CAG GCG ATG GAC AAC GGT ACC

          S   P   S   F   L   L   P   R   R   L   Q   R   S   R   G   N   R   A   G   G
Query     TCT CCA TCG TTC CTC TTG CCA CGC CGT TTA CAA CGT TCA AGA GGC AAC CGC GCT GGT GGC
          ||| ||# ||* ||| ||* #|| ||# ||# ||| #|# ||# ||| ||* *|* ||# ||| ||# ||* ||| ||#
Optimized TCT CCG TCT TTC CTG CTG CCG CGT CGT CTG CAG CGT TCT CGT GGT AAC CGT GCG GGT GGT

          Y   A   V   G   M   V   I   V   G   S   A   H   Q   V   N   R   H   F   T   A
Query     TAT GCA GTT GGA ATG GTC ATT GTG GGC TCA GCA CAC CAG GTG AAC CGC CAC TTC ACT GCT
          ||# ||# ||| ||* ||| ||# ||# ||* ||# ||* ||# ||| ||| ||* ||| ||# ||| ||| ||# ||*
Optimized TAC GCG GTT GGT ATG GTT ATC GTT GGT TCT GCG CAC CAG GTT AAC CGT CAC TTC ACC GCG

          L   G   T   T   F   G   W   S   T   A   P   Y   E   P   M   R   C   A   F   G
Query     CTT GGC ACA ACA TTT GGT TGG TCT ACC GCA CCT TAC GAA CCC ATG CGA TGT GCT TTT GGT
          ||* ||# ||* ||* ||# ||| ||| ||| ||| ||# ||* ||| ||| ||* ||| ||* ||# ||* ||# |||
Optimized CTG GGT ACC ACC TTC GGT TGG TCT ACC GCG CCG TAC GAA CCG ATG CGT TGC GCG TTC GGT

          G   V   H   Q   G   R   D   T   N   P   K   I   G   Y   Y   W   E   V   G   A
Query     GGA GTA CAT CAA GGG CGT GAC ACC AAC CCA AAA ATT GGC TAC TAC TGG GAG GTT GGT GCT
          ||* ||* ||# ||# ||* ||| ||| ||| ||| ||# ||| ||# ||# ||| ||| ||| ||# ||| ||| ||*
Optimized GGT GTT CAC CAG GGT CGT GAC ACC AAC CCG AAA ATC GGT TAC TAC TGG GAA GTT GGT GCG
```

```
          D   Q   R   G   P   L   F   P   N   I   V   N   H   W   P   D   F   A   V   N
    Query GAC CAG CGT GGC CCG CTT TTT CCA AAC ATT GTT AAT CAT TGG CCT GAT TTT GCA GTC AAC
          ||| ||| ||| ||# ||| ||* ||# ||# ||| ||# ||| ||# ||# ||| ||* ||# ||# ||# ||# |||
Optimized GAC CAG CGT GGT CCG CTG TTC CCG AAC ATC GTT AAC CAC TGG CCG GAC TTC GCG GTT AAC

          T   K   Y   T   W   P   D   A   D   Y   I   P   H   S   A   V   V   G   T   L
    Query ACC AAA TAC ACA TGG CCA GAT GCC GAC TAT ATA CCC CAT AGT GCG GTT GTT GGA ACC TTG
          ||| ||| ||| ||* ||| ||# ||# ||* ||| ||# ||* ||* ||# **| ||| ||| ||| ||* ||| #||
Optimized ACC AAA TAC ACC TGG CCG GAC GCG GAC TAC ATC CCG CAC TCT GCG GTT GTT GGT ACC CTG

          V   S   F   Q   D   N   G   D   V   S   E   D   Q   V   A   T   A   F   A   I
    Query GTT TCC TTC CAG GAC AAT GGA GAT GTT TCT GAG GAT CAA GTG GCT ACT GCA TTT GCA ATT
          ||| ||# ||| ||| ||| ||# ||* ||# ||| ||| ||# ||# ||# ||* ||* ||# ||# ||# ||# ||#
Optimized GTT TCT TTC CAG GAC AAC GGT GAC GTT TCT GAA GAC CAG GTT GCG ACC GCG TTC GCG ATC

          S   M   N   T   P   S   G   S   T   T   G   R   G   T   V   R   E   A   F   D
    Query TCA ATG AAC ACT CCA AGT GGG AGC ACA ACT GGA CGC GGC ACA GTC AGA GAA GCA TTT GAT
          ||* ||| ||| ||# ||# **| ||* **# ||* ||# ||* ||# ||# ||* ||# *|* ||| ||# ||# ||#
Optimized TCT ATG AAC ACC CCG TCT GGT TCT ACC ACC GGT CGT GGT ACC GTT CGT GAA GCG TTC GAC

          P   S   T   M   H   L   V   R   T   N   G   T   T   Q   P   S   G   W   P   T
    Query CCA TCC ACA ATG CAT TTG GTC CGC ACC AAT GGT ACC ACT CAA CCA TCT GGG TGG CCA ACC
          ||# ||# ||* ||| ||# #|| ||# ||# ||| ||# ||| ||| ||# ||# ||# ||| ||* ||| ||# |||
Optimized CCG TCT ACC ATG CAC CTG GTT CGT ACC AAC GGT ACC ACC CAG CCG TCT GGT TGG CCG ACC

          G   S   N   T   G   N   G   Y   F   T   P   M   W   G   H   G   Q   G   N   A
    Query GGC TCA AAC ACT GGA AAT GGT TAC TTC ACA CCA ATG TGG GGT CAT GGT CAG GGT AAT GCA
          ||# ||* ||| ||# ||* ||# ||| ||| ||| ||* ||# ||| ||| ||| ||# ||| ||| ||| ||# ||#

          I   N   D   K   I   T   N   M   E   G   A   N   Y   T   F   G   G   S   G   Q
    Query ATT AAT GAT AAG ATC ACC AAC ATG GAG GGG GCT AAT TAT ACA TTT GGA GGC TCT GGC CAA
          ||# ||# ||# ||# ||| ||| ||| ||| ||# ||* ||* ||# ||# ||* ||# ||* ||# ||| ||# ||#
Optimized ATC AAC GAC AAA ATC ACC AAC ATG GAA GGT GCG AAC TAC ACC TTC GGT GGT TCT GGT CAG

          N   N   I   V   L   W   V   E   R   I   F   S   D   H   P   G   K   T   T   L
    Query AAC AAC ATA GTC TTG TGG GTT GAA AGA ATC TTC TCA GAC CAT CCT GGT AAA ACC ACA CTT
          ||| ||| ||* ||# #|| ||| ||| ||| *|* ||| ||| ||* ||| ||# ||* ||| ||| ||| ||* ||*
Optimized AAC AAC ATC GTT CTG TGG GTT GAA CGT ATC TTC TCT GAC CAC CCG GGT AAA ACC ACC CTG

          Y   S   S   Q   L   D   S   T   A   A   I   F   Q   S   G   P   V   N   I   P
    Query TAC TCA TCA CAA TTG GAC AGC ACT GCG GCC ATT TTC CAG TCA GGC CCA GTC AAT ATA CCT
          ||| ||* ||* ||# #|| ||| **# ||# ||| ||* ||# ||| ||| ||* ||# ||# ||# ||# ||* ||*
Optimized TAC TCT TCT CAG CTG GAC TCT ACC GCG GCG ATC TTC CAG TCT GGT CCG GTT AAC ATC CCG
```

```
           E   N   M   M   A   V   Y   N   V   T   T   N   G   A   D   F   Q   V   G   I
    Query  GAG AAC ATG ATG GCT GTC TAC AAT GTC ACC ACT AAT GGG GCT GAC TTT CAA GTT GGC ATC
           ||# ||| ||| ||| ||* ||# ||| ||# ||# ||| ||# ||# ||* ||* ||| ||# ||# ||| ||# |||
Optimized  GAA AAC ATG ATG GCG GTT TAC AAC GTT ACC ACC AAC GGT GCG GAC TTC CAG GTT GGT ATC

           R   R   D   G   Y   M   V   T   S   G   T   I   G   T   Q   Q   E   L   D   P
    Query  CGT CGT GAT GGC TAC ATG GTC ACA TCT GGA ACA ATT GGT ACT CAG CAG GAG CTT GAC CCT
           ||| ||| ||# ||# ||| ||| ||# ||* ||| ||* ||* ||# ||| ||# ||| ||| ||# ||* ||| ||*
Optimized  CGT CGT GAC GGT TAC ATG GTT ACC TCT GGT ACC ATC GGT ACC CAG CAG GAA CTG GAC CCG

           D   T   T   F   T   Y   V   G   L   F   P   L   S   A   S   L   V   G   P   H
    Query  GAC ACC ACT TTC ACT TAT GTT GGA CTT TTC CCC CTT TCT GCC TCA TTG GTT GGC CCA CAT
           ||| ||| ||# ||| ||# ||# ||| ||* ||* ||| ||* ||* ||| ||* ||* #|| ||| ||# ||# ||#
Optimized  GAC ACC ACC TTC ACC TAC GTT GGT CTG TTC CCG CTG TCT GCG TCT CTG GTT GGT CCG CAC

           G   N   S   G   R   A   Q   I   A   W   S   .
    Query  GGG AAT TCT GGA CGG GCC CAG ATA GCA TGG TCA TGA
           ||* ||# ||| ||* ||* ||* ||| ||* ||# ||| ||* |||
Optimized  GGT AAC TCT GGT CGT GCG CAG ATC GCG TGG TCT TGA
```
