## Supplemental Figure S2 for "In silico codon optimization and phylogenetic analysis of various strains of Sapovirus"

**Supplementary Figure S2.** Sequence alignment of the native-type (Query) and codon-optimized VP2 gene sequences of twelve human Sapovirus strains (GenBank accession numbers AY237423.2, AY237422.3, AY646853.2, AY646854.2, KP298674.1, AY694184.2, HM002617.1, AJ606693.2, LC190463.1, AY237420.2, AY603425.1 and AB775659.2).

**The specific-coloured symbols stand for (*): Transversion change, (#): Transition change and (|): Unchanged nucleotide.*

**N21 (AY237423.2)** – 498 nucleotides; 382 unchanged, 59 transition and 57 transversion changes.

```
           M   S   W   L   V   G   A   L   Q   T   F   G   S   L   A   D   V   A   G   T
    Query  ATG AGT TGG TTA GTG GGT GCC CTC CAA ACC TTC GGG TCA TTG GCT GAC GTT GCT GGC ACT
           ||| **| ||| #|# ||* ||| ||* ||* ||# ||| ||| ||* ||* #|| ||* ||| ||| ||* ||# ||#
Optimized  ATG TCT TGG CTG GTT GGT GCG CTG CAG ACC TTC GGT TCT CTG GCG GAC GTT GCG GGT ACC

           V   S   N   I   V   Y   Q   Q   R   Q   A   A   Q   L   E   K   Q   N   E   L
    Query  GTA TCC AAT ATA GTG TAC CAG CAG CGC CAG GCT GCT CAG TTG GAA AAA CAA AAT GAA CTG
           ||* ||# ||# ||* ||* ||| ||| ||| ||# ||| ||* ||* ||| #|| ||| ||| ||# ||# ||| |||
Optimized  GTT TCT AAC ATC GTT TAC CAG CAG CGT CAG GCG GCG CAG CTG GAA AAA CAG AAC GAA CTG

           M   E   T   W   I   N   K   Q   E   A   L   Q   K   S   Q   M   E   L   T   R
    Query  ATG GAA ACA TGG ATA AAC AAG CAA GAA GCA CTG CAG AAA TCC CAA ATG GAA TTA ACC CGA
           ||| ||| ||* ||| ||* ||| ||# ||# ||| ||# ||| ||| ||| ||# ||# ||| ||| #|# ||| ||*
Optimized  ATG GAA ACC TGG ATC AAC AAA CAG GAA GCG CTG CAG AAA TCT CAG ATG GAA CTG ACC CGT

           D   L   S   I   N   G   P   A   A   R   V   Q   S   A   L   D   A   G   F   D
    Query  GAC CTC TCC ATT AAT GGT CCA GCT GCC CGG GTG CAA TCA GCA CTC GAC GCG GGT TTT GAT
           ||| ||* ||# ||# ||# ||| ||# ||* ||* ||* ||* ||# ||* ||# ||* ||| ||| ||| ||# ||#
Optimized  GAC CTG TCT ATC AAC GGT CCG GCG GCG CGT GTT CAG TCT GCG CTG GAC GCG GGT TTC GAC

           E   V   S   A   R   R   I   A   G   S   G   E   R   V   I   W   G   N   L   D
    Query  GAA GTG AGT GCA CGG CGA ATA GCT GGT TCA GGT GAG CGT GTG ATA TGG GGA AAC TTG GAT
           ||| ||* **| ||# ||* ||* ||* ||* ||| ||* ||| ||# ||| ||* ||* ||| ||* ||| #|| ||#
Optimized  GAA GTT TCT GCG CGT CGT ATC GCG GGT TCT GGT GAA CGT GTT ATC TGG GGT AAC CTG GAC

           R   P   I   M   H   A   G   T   M   D   S   I   R   Q   T   K   H   L   D   S
    Query  CGA CCC ATC ATG CAC GCT GGC ACT ATG GAT AGC ATA CGC CAA ACC AAA CAT TTG GAC TCT
           ||* ||* ||| ||| ||| ||* ||# ||# ||| ||# **# ||* ||# ||# ||| ||| ||# #|| ||| |||
Optimized  CGT CCG ATC ATG CAC GCG GGT ACC ATG GAC TCT ATC CGT CAG ACC AAA CAC CTG GAC TCT

           L   S   H   S   L   A   T   F   K   N   G   T   P   F   G   K   P   A   P   P
    Query  TTG AGT CAC TCA TTG GCT ACA TTC AAA AAT GGG ACT CCA TTT GGG AAA CCT GCG CCC CCA
           #|| **| ||| ||* #|| ||* ||* ||| ||| ||# ||* ||# ||# ||# ||* ||| ||* ||| ||* ||#
Optimized  CTG TCT CAC TCT CTG GCG ACC TTC AAA AAC GGT ACC CCG TTC GGT AAA CCG GCG CCG CCG

           T   A   K   F   G   K   P   Q   A   T   T   A   Q   V   N   I   G   H   N   P
    Query  ACA GCA AAA TTT GGC AAG CCC CAG GCC ACA ACT GCA CAA GTT AAC ATT GGT CAC AAC CCT
           ||* ||# ||| ||# ||# ||# ||* ||| ||* ||* ||# ||# ||# ||| ||| ||# ||| ||| ||| ||*
Optimized  ACC GCG AAA TTC GGT AAA CCG CAG GCG ACC ACC GCG CAG GTT AAC ATC GGT CAC AAC CCG

           G   S   S   N   V   .
    Query  GGC AGC TCC AAT GTC TAA
           ||# **# ||# ||# ||# |||
Optimized  GGT TCT TCT AAC GTT TAA
```

**Mc114 (AY237422.3)** – 498 nucleotides; 384 unchanged, 58 transition and 56 transversion changes.

```
           M   S   W   L   V   G   A   L   Q   T   F   G   S   L   A   D   V   A   G   T
Query      ATG AGT TGG TTA GTG GGT GCC CTC CAG ACC TTC GGG TCA TTG GCT GAC GTT GCT GGC ACT
           ||| **| ||| #|# ||* ||| ||* ||* ||| ||| ||| ||* ||* #|| ||* ||| ||| ||* ||# ||#
Optimized  ATG TCT TGG CTG GTT GGT GCG CTG CAG ACC TTC GGT TCT CTG GCG GAC GTT GCG GGT ACC

           V   S   N   I   V   Y   Q   Q   R   Q   A   A   Q   L   E   K   Q   N   E   L
Query      GTG TCT AAT ATA GTG TAC CAG CAG CGC CAG GCT GCT CAG TTG GAA AAA CAA AAT GAA CTG
           ||* ||| ||# ||* ||* ||| ||| ||| ||# ||| ||* ||* ||| #|| ||| ||| ||# ||# ||| |||
Optimized  GTT TCT AAC ATC GTT TAC CAG CAG CGT CAG GCG GCG CAG CTG GAA AAA CAG AAC GAA CTG

           M   E   T   W   M   N   K   Q   E   A   L   Q   K   S   Q   M   E   L   T   R
Query      ATG GAA ACA TGG ATG AAC AAG CAA GAA GCA TTG CAG AAA TCC CAA ATG GAA TTA ACC CGA
           ||| ||| ||* ||| ||| ||| ||# ||# ||| ||# #|| ||| ||| ||# ||# ||| ||| #|# ||| ||*
Optimized  ATG GAA ACC TGG ATG AAC AAA CAG GAA GCG CTG CAG AAA TCT CAG ATG GAA CTG ACC CGT

           D   L   S   I   N   G   P   A   A   R   V   Q   S   A   L   D   A   G   F   D
Query      GAC CTC TCC ATT AAT GGT CCA GCT GCC CGG GTG CAA TCA GCA CTC GAC GCG GGT TTC GAT
           ||| ||* ||# ||# ||# ||| ||# ||* ||* ||* ||* ||# ||* ||# ||* ||| ||| ||| ||| ||#
Optimized  GAC CTG TCT ATC AAC GGT CCG GCG GCG CGT GTT CAG TCT GCG CTG GAC GCG GGT TTC GAC

           E   V   S   A   R   R   I   A   G   S   G   E   R   V   I   W   G   N   L   D
Query      GAA GTG AGT GCA CGG CGA ATA GCT GGT TCA GGT GAG CGT GTG ATA TGG GGG AAC TTG GAT
           ||| ||* **| ||# ||* ||* ||* ||* ||| ||* ||| ||# ||| ||* ||* ||| ||* ||| #|| ||#
Optimized  GAA GTT TCT GCG CGT CGT ATC GCG GGT TCT GGT GAA CGT GTT ATC TGG GGT AAC CTG GAC

           R   P   I   M   H   A   G   T   M   D   S   I   R   Q   T   K   H   L   D   S
Query      CGA CCC ATC ATG CAC GCT GGC ACT ATG GAT AGC ATA CGC CAA ACC AAA CAT TTG GAC TCT
           ||* ||* ||| ||| ||| ||* ||# ||# ||| ||# **# ||* ||# ||# ||| ||| ||# #|| ||| |||
Optimized  CGT CCG ATC ATG CAC GCG GGT ACC ATG GAC TCT ATC CGT CAG ACC AAA CAC CTG GAC TCT

           L   S   H   S   L   A   T   F   K   N   G   T   P   F   G   K   P   A   P   P
Query      TTG AGT CAC TCA TTG GCT ACG TTC AAA AAT GGA ACT CCA TTT GGG AAA CCT GCG CCC CCA
           #|| **| ||| ||* #|| ||* ||* ||| ||| ||# ||* ||# ||# ||# ||* ||| ||* ||| ||* ||#
Optimized  CTG TCT CAC TCT CTG GCG ACC TTC AAA AAC GGT ACC CCG TTC GGT AAA CCG GCG CCG CCG

           T   A   K   F   G   K   P   Q   A   T   T   A   Q   I   N   I   G   H   N   P
Query      ACA GCA AAA TTT GGC AAG CCC CAG GCC ACA ACT GCA CAA ATT AAC ATT GGT CAC AAC CCT
           ||* ||# ||| ||# ||# ||# ||* ||| ||* ||* ||# ||# ||# ||# ||| ||# ||| ||| ||| ||*
Optimized  ACC GCG AAA TTC GGT AAA CCG CAG GCG ACC ACC GCG CAG ATC AAC ATC GGT CAC AAC CCG

           G   S   S   N   V   .
Query      GGC AGC TCC AAT GTC TAA
           ||# **# ||# ||# ||# |||
Optimized  GGT TCT TCT AAC GTT TAA
```

**NongKhai-50 (AY646853.2)** – 498 nucleotides; 383 unchanged, 59 transition and 56 transversion changes.

```
           M   S   W   L   V   G   A   L   Q   T   F   G   S   L   A   D   V   A   G   T
    Query  ATG AGT TGG TTG GTA GGT GCT CTG CAA ACT TTT GGG TCT TTG GCT GAT GTT GCT GGC ACT
           ||| **| ||| #|| ||* ||| ||* ||| ||# ||# ||# ||* ||| #|| ||* ||# ||| ||* ||# ||#
Optimized  ATG TCT TGG CTG GTT GGT GCG CTG CAG ACC TTC GGT TCT CTG GCG GAC GTT GCG GGT ACC

           V   S   N   I   V   Y   Q   Q   R   Q   A   A   Q   L   E   K   Q   N   E   L
    Query  GTT TCC AAC ATA GTG TAC CAG CAG CGC CAG GCC GCT CAG TTG GAA AAA CAA AAT GAA CTT
           ||| ||# ||| ||* ||* ||| ||| ||| ||# ||| ||* ||* ||| #|| ||| ||| ||# ||# ||| ||*
Optimized  GTT TCT AAC ATC GTT TAC CAG CAG CGT CAG GCG GCG CAG CTG GAA AAA CAG AAC GAA CTG

           M   E   T   W   M   N   K   Q   E   A   L   Q   K   S   Q   M   E   L   T   R
    Query  ATG GAG ACA TGG ATG AAC AAA CAA GAG GCA CTG CAG AAG TCT CAA ATG GAA TTA ACC CGA
           ||| ||# ||* ||| ||| ||| ||| ||# ||# ||# ||| ||| ||# ||| ||# ||| ||| #|# ||| ||*
Optimized  ATG GAA ACC TGG ATG AAC AAA CAG GAA GCG CTG CAG AAA TCT CAG ATG GAA CTG ACC CGT

           D   L   S   I   N   G   P   A   A   R   V   Q   S   A   L   D   A   G   F   D
    Query  GAC CTC TCC ATC AAT GGT CCA GCC GCC CGG GTA CAA TCA GCA CTT GAT GCA GGT TTT GAT
           ||| ||* ||# ||| ||# ||| ||# ||* ||* ||* ||* ||# ||* ||# ||* ||# ||# ||| ||# ||#
Optimized  GAC CTG TCT ATC AAC GGT CCG GCG GCG CGT GTT CAG TCT GCG CTG GAC GCG GGT TTC GAC

           E   V   S   A   R   R   I   A   G   S   G   E   R   V   I   W   G   N   L   D
    Query  GAG GTA AGT GCA CGG CGC ATT GCT GGT TCA GGT GAG CGC GTG ATT TGG GGG AAC TTG GAT
           ||# ||* **| ||# ||* ||# ||# ||* ||| ||* ||| ||# ||# ||* ||# ||| ||* ||| #|| ||#
Optimized  GAA GTT TCT GCG CGT CGT ATC GCG GGT TCT GGT GAA CGT GTT ATC TGG GGT AAC CTG GAC

           R   P   I   M   H   A   G   T   M   D   S   I   R   Q   T   K   H   L   D   S
    Query  CGA CCC ATC ATG CAC GCT GGC ACT ATG GAC AGC ATA CGC CAG ACT AAA CAT TTG GAC TCT
           ||* ||* ||| ||| ||| ||* ||# ||# ||| ||| **# ||* ||# ||| ||# ||| ||# #|| ||| |||
Optimized  CGT CCG ATC ATG CAC GCG GGT ACC ATG GAC TCT ATC CGT CAG ACC AAA CAC CTG GAC TCT

           L   S   H   S   L   A   T   F   K   N   G   T   P   F   G   K   P   A   P   P
    Query  CTG AGT CAT TCA TTG GCC ACA TTT AAA AAT GGA ACT CCA TTC GGA AAA CCT GCA CCC CCA
           ||| **| ||# ||* #|| ||* ||* ||# ||| ||# ||* ||# ||# ||| ||* ||| ||* ||# ||* ||#
Optimized  CTG TCT CAC TCT CTG GCG ACC TTC AAA AAC GGT ACC CCG TTC GGT AAA CCG GCG CCG CCG

           T   A   K   S   G   R   P   Q   A   T   T   A   K   I   T   I   G   H   N   P
    Query  ACA GCA AAA TCT GGA AGG CCC CAG GCC ACA ACT GCA AAA ATC ACC ATT GGT CAC AAC CCT
           ||* ||# ||| ||| ||* *|* ||* ||| ||* ||* ||# ||# ||| ||| ||| ||# ||| ||| ||| ||*
Optimized  ACC GCG AAA TCT GGT CGT CCG CAG GCG ACC ACC GCG AAA ATC ACC ATC GGT CAC AAC CCG

           G   S   T   S   V   .
    Query  GGT AGC ACC AGT GTT TAA
           ||| **# ||| **| ||| |||
Optimized  GGT TCT ACC TCT GTT TAA
```

**Chanthaburi-74 (AY646854.2)** – 498 nucleotides; 380 unchanged, 64 transition and 54 transversion changes.

```
           M   S   W   L   V   G   A   L   Q   T   F   G   S   L   A   D   V   A   G   T
    Query  ATG AGT TGG TTG GTG GGT GCT CTG CAG ACT TTT GGG TCT TTG GCA GAC GTT GCT GGC ACT
           ||| **| ||| #|| ||* ||| ||* ||| ||| ||# ||# ||* ||| #|| ||# ||| ||| ||* ||# ||#
Optimized  ATG TCT TGG CTG GTT GGT GCG CTG CAG ACC TTC GGT TCT CTG GCG GAC GTT GCG GGT ACC

           V   S   N   I   V   Y   Q   Q   R   Q   A   A   Q   L   E   K   Q   N   E   L
    Query  GTT TCC AAC ATA GTG TAC CAG CAG CGC CAG GCT GCT CAG TTG GAA AAA CAA AAT GAA CTA
           ||| ||# ||| ||* ||* ||| ||| ||| ||# ||| ||* ||* ||| #|| ||| ||| ||# ||# ||| ||#
Optimized  GTT TCT AAC ATC GTT TAC CAG CAG CGT CAG GCG GCG CAG CTG GAA AAA CAG AAC GAA CTG

           M   E   T   W   M   N   K   Q   E   A   L   Q   K   S   Q   M   E   L   T   R
    Query  ATG GAG ACA TGG ATG AAT AAA CAA GAG GCA CTG CAG AAA TCT CAA ATG GAA TTA ACT CGG
           ||| ||# ||* ||| ||| ||# ||| ||# ||# ||# ||| ||| ||| ||| ||# ||| ||| #|# ||# ||*
Optimized  ATG GAA ACC TGG ATG AAC AAA CAG GAA GCG CTG CAG AAA TCT CAG ATG GAA CTG ACC CGT

           D   L   S   I   N   G   P   A   A   R   V   Q   S   A   L   D   A   G   F   D
    Query  GAC CTC TCC ATC AAT GGT CCA GCC GCC CGG GTG CAA TCA GCA CTC GAT GCA GGT TTT GAT
           ||| ||* ||# ||| ||# ||| ||# ||* ||* ||* ||* ||# ||* ||# ||* ||# ||# ||| ||# ||#
Optimized  GAC CTG TCT ATC AAC GGT CCG GCG GCG CGT GTT CAG TCT GCG CTG GAC GCG GGT TTC GAC

           E   V   S   A   R   R   I   A   G   S   G   E   R   V   I   W   G   N   L   D
    Query  GAA GTG AGT GCA CGG CGC ATT GCT GGT TCA GGT GAA CGC GTG ATA TGG GGG AAT TTG GAT
           ||| ||* **| ||# ||* ||# ||# ||* ||| ||* ||| ||| ||# ||* ||* ||| ||* ||# #|| ||#
Optimized  GAA GTT TCT GCG CGT CGT ATC GCG GGT TCT GGT GAA CGT GTT ATC TGG GGT AAC CTG GAC

           R   P   I   M   H   A   G   T   M   D   S   I   R   Q   T   K   H   L   D   S
    Query  CGA CCC ATT ATG CAC GCT GGC ACT ATG GAT AGC ATA CGC CAA ACT AAA CAT TTG GAC TCT
           ||* ||* ||# ||| ||| ||* ||# ||# ||| ||# **# ||* ||# ||# ||# ||| ||# #|| ||| |||
Optimized  CGT CCG ATC ATG CAC GCG GGT ACC ATG GAC TCT ATC CGT CAG ACC AAA CAC CTG GAC TCT

           L   S   H   S   L   A   T   F   K   N   G   T   P   F   G   K   P   A   P   P
    Query  CTG AGT CAT TCT TTG GCC ACA TTC AAA AAT GGA ACT CCA TTT GGA AAA CCT GCA CCC CCA
           ||| **| ||# ||| #|| ||* ||* ||| ||| ||# ||* ||# ||# ||# ||* ||| ||* ||# ||* ||#
Optimized  CTG TCT CAC TCT CTG GCG ACC TTC AAA AAC GGT ACC CCG TTC GGT AAA CCG GCG CCG CCG

           T   A   K   S   G   R   P   Q   A   T   T   A   Q   I   N   I   G   H   N   P
    Query  ACA GCA AAG TCT GGA AGG CCT CAG GCC ACA ACT GCA CAA ATC AAT ATT GGC CAC AAC CCT
           ||* ||# ||# ||| ||* *|* ||* ||| ||* ||* ||# ||# ||# ||| ||# ||# ||# ||| ||| ||*
Optimized  ACC GCG AAA TCT GGT CGT CCG CAG GCG ACC ACC GCG CAG ATC AAC ATC GGT CAC AAC CCG

           G   S   T   S   V   .
    Query  GGT AGT ACC AGT GTT TAA
           ||| **| ||| **| ||| |||
Optimized  GGT TCT ACC TCT GTT TAA
```

**ROK62 (KP298674.1)** – 498 nucleotides; 382 unchanged, 61 transition and 55 transversion changes.

```
           M   S   W   L   V   G   A   L   Q   T   F   G   S   L   A   D   V   A   G   T
    Query  ATG AGT TGG TTG GTA GGC GCC CTC CAG ACC TTC GGG TCA TTG GCT GAC GTT GCT GGC ACT
           ||| **| ||| #|| ||* ||# ||* ||* ||| ||| ||| ||* ||* #|| ||* ||| ||| ||* ||# ||#
Optimized  ATG TCT TGG CTG GTT GGT GCG CTG CAG ACC TTC GGT TCT CTG GCG GAC GTT GCG GGT ACC

           V   S   N   I   V   Y   Q   Q   R   Q   A   A   Q   L   E   K   Q   N   E   L
    Query  GTG TCT AAC ATA GTG TAC CAG CAG CGT CAG GCT GCT CAG TTG GAA AAA CAG AAT GAA CTA
           ||* ||| ||| ||* ||* ||| ||| ||| ||| ||| ||* ||* ||| #|| ||| ||| ||| ||# ||| ||#
Optimized  GTT TCT AAC ATC GTT TAC CAG CAG CGT CAG GCG GCG CAG CTG GAA AAA CAG AAC GAA CTG

           M   E   T   W   M   N   K   Q   E   A   L   Q   K   S   Q   M   E   L   T   R
    Query  ATG GAG ACA TGG ATG AAC AAG CAA GAA GCA TTG CAA AAG TCC CAA ATG GAA TTA ACC CGA
           ||| ||# ||* ||| ||| ||| ||# ||# ||| ||# #|| ||# ||# ||# ||# ||| ||| #|# ||| ||*
Optimized  ATG GAA ACC TGG ATG AAC AAA CAG GAA GCG CTG CAG AAA TCT CAG ATG GAA CTG ACC CGT

           D   L   S   I   N   G   P   A   A   R   V   Q   S   A   L   D   A   G   F   D
    Query  GAC CTC TCC ATT AAT GGT CCA GCT GCC CGG GTG CAA TCA GCA CTT GAC GCA GGT TTT GAT
           ||| ||* ||# ||# ||# ||| ||# ||* ||* ||* ||* ||# ||* ||# ||* ||| ||# ||| ||# ||#
Optimized  GAC CTG TCT ATC AAC GGT CCG GCG GCG CGT GTT CAG TCT GCG CTG GAC GCG GGT TTC GAC

           A   V   S   A   R   R   I   A   G   S   G   E   R   V   I   W   G   N   L   D
    Query  GCA GTG AGT GCA CGG CGT ATT GCT GGT TCA GGT GAG CGC GTG ATA TGG GGC AAT CTT GAT
           ||# ||* **| ||# ||* ||| ||# ||* ||| ||* ||| ||# ||# ||* ||* ||| ||# ||# ||* ||#
Optimized  GCG GTT TCT GCG CGT CGT ATC GCG GGT TCT GGT GAA CGT GTT ATC TGG GGT AAC CTG GAC

           R   P   I   M   H   A   G   T   M   D   S   I   R   Q   T   K   H   L   D   F
    Query  CGG CCC ATC ATG CAC GCT GGC ACT ATG GAC AGC ATA CGC CAA ACT AAA CAC TTG GAC TTT
           ||* ||* ||| ||| ||| ||* ||# ||# ||| ||| **# ||* ||# ||# ||# ||| ||| #|| ||| ||#
Optimized  CGT CCG ATC ATG CAC GCG GGT ACC ATG GAC TCT ATC CGT CAG ACC AAA CAC CTG GAC TTC

           L   S   H   S   L   A   T   F   K   N   G   T   P   F   G   K   P   A   P   P
    Query  TTG AGT CAC TCA TTG GCC ACA TTC AAA AAT GGA ACT CCA TTT GGT AAA CCT GCA CCC CCA
           #|| **| ||| ||* #|| ||* ||* ||| ||| ||# ||* ||# ||# ||# ||| ||| ||* ||# ||* ||#
Optimized  CTG TCT CAC TCT CTG GCG ACC TTC AAA AAC GGT ACC CCG TTC GGT AAA CCG GCG CCG CCG

           T   A   K   F   G   K   P   Q   A   T   I   A   K   I   N   I   G   H   N   P
    Query  ACA GCA AAA TTT GGC AAA CCT CAG GCC ACA ATT GCA AAA ATC AAC ATT GGT CAC AAC CCT
           ||* ||# ||| ||# ||# ||| ||* ||| ||* ||* ||# ||# ||| ||| ||| ||# ||| ||| ||| ||*
Optimized  ACC GCG AAA TTC GGT AAA CCG CAG GCG ACC ATC GCG AAA ATC AAC ATC GGT CAC AAC CCG

           G   S   S   S   V   .
    Query  GGT AGC TCC AGT GTC TAA
           ||| **# ||# **| ||# |||
Optimized  GGT TCT TCT TCT GTT TAA
```

**Dresden (AY694184.2)** – 498 nucleotides; 379 unchanged, 63 transition and 56 transversion changes.

```
           M   S   W   L   V   G   A   L   Q   T   F   G   S   L   A   D   V   A   G   T
Query      ATG AGT TGG TTG GTA GGC GCC CTG CAG ACT TTC GGG TCT TTG GCT GAT GTC GCT GGC ACT
           ||| **| ||| #|| ||* ||# ||* ||| ||| ||# ||| ||* ||| #|| ||* ||# ||# ||* ||# ||#
Optimized  ATG TCT TGG CTG GTT GGT GCG CTG CAG ACC TTC GGT TCT CTG GCG GAC GTT GCG GGT ACC

           V   S   N   I   V   Y   Q   Q   R   Q   A   A   Q   L   E   K   Q   N   E   L
Query      GTC TCC AAC ATA GTG TAC CAG CAG CGC CAG GCC GCC CAG TTG GAA AAA CAG AAT GAA CTA
           ||# ||# ||| ||* ||* ||| ||| ||| ||# ||| ||* ||* ||| #|| ||| ||| ||| ||# ||| ||#
Optimized  GTT TCT AAC ATC GTT TAC CAG CAG CGT CAG GCG GCG CAG CTG GAA AAA CAG AAC GAA CTG

           M   E   T   W   M   N   K   Q   E   A   L   Q   K   S   Q   M   E   L   T   R
Query      ATG GAG ACA TGG ATG AAC AAA CAA GAG GCA CTG CAG AAG TCC CAG ATG GAG TTA ACT CGA
           ||| ||# ||* ||| ||| ||| ||| ||# ||# ||# ||| ||| ||# ||# ||| ||| ||# #|# ||# ||*
Optimized  ATG GAA ACC TGG ATG AAC AAA CAG GAA GCG CTG CAG AAA TCT CAG ATG GAA CTG ACC CGT

           D   L   S   I   N   G   P   A   A   R   V   Q   S   A   L   D   A   G   F   D
Query      GAC CTC TCC ATC AAT GGT CCA GCC GCC CGG GTG CAA TCG GCA CTT GAT GCA GGT TTT GAT
           ||| ||* ||# ||| ||# ||| ||# ||* ||* ||* ||* ||# ||* ||# ||* ||# ||# ||| ||# ||#
Optimized  GAC CTG TCT ATC AAC GGT CCG GCG GCG CGT GTT CAG TCT GCG CTG GAC GCG GGT TTC GAC

           E   V   S   A   R   R   I   A   G   S   G   E   R   V   I   W   G   N   L   D
Query      GAA GTG AGT GCA CGG CGT ATT GCT GGT TCA GGT GAG CGT GTG ATC TGG GGG AAT TTG GAT
           ||| ||* **| ||# ||* ||| ||# ||* ||| ||* ||| ||# ||| ||* ||| ||| ||* ||# #|| ||#
Optimized  GAA GTT TCT GCG CGT CGT ATC GCG GGT TCT GGT GAA CGT GTT ATC TGG GGT AAC CTG GAC

           R   P   I   M   H   A   G   T   M   D   S   I   R   Q   T   R   H   L   D   S
Query      CGA CCC ATC ATG CAT GCT GGC ACT ATG GAC AGC ATA CGC CAG ACT AGA CAT TTA GAC TCT
           ||* ||* ||| ||| ||# ||* ||# ||# ||| ||| **# ||* ||# ||| ||# *|* ||# #|# ||| |||
Optimized  CGT CCG ATC ATG CAC GCG GGT ACC ATG GAC TCT ATC CGT CAG ACC CGT CAC CTG GAC TCT

           L   S   H   S   L   A   T   F   K   N   G   T   P   F   G   K   P   A   P   P
Query      CTG AGT CAT TCA TTG GCT ACA TTC AAG AAT GGA ACT CCA TTC GGG AAA CCA GCG CCC CCA
           ||| **| ||# ||* #|| ||* ||* ||| ||# ||# ||* ||# ||# ||| ||* ||| ||# ||| ||* ||#
Optimized  CTG TCT CAC TCT CTG GCG ACC TTC AAA AAC GGT ACC CCG TTC GGT AAA CCG GCG CCG CCG

           T   A   K   S   G   R   P   Q   A   T   T   A   Q   I   T   I   G   H   N   P
Query      ACA GCA AAA TCT GGA AGG CCC CAG GCC ACA ACT GCA CAA ATC ACC ATT GGT CAT AAC CCT
           ||* ||# ||| ||| ||* *|* ||* ||| ||* ||* ||# ||# ||# ||| ||| ||# ||| ||# ||| ||*
Optimized  ACC GCG AAA TCT GGT CGT CCG CAG GCG ACC ACC GCG CAG ATC ACC ATC GGT CAC AAC CCG

           G   S   T   S   V   .
Query      GGT AGC ACC AGT GTT TAA
           ||| **# ||| **| ||| |||
Optimized  GGT TCT ACC TCT GTT TAA
```

**Sapporo-MT-2010 (HM002617.1)** – 498 nucleotides; 382 unchanged, 62 transition and 54 transversion changes.

```
           M   S   W   L   V   G   A   L   Q   T   F   G   S   L   A   D   V   A   G   T
     Query ATG AGT TGG TTG GTG GGT GCC CTT CAA ACT TTT GGG TCT TTG GCT GAC GTT GCT GGC ACT
           ||| **| ||| #|| ||* ||| ||* ||* ||# ||# ||# ||* ||| #|| ||* ||| ||| ||* ||# ||#
 Optimized ATG TCT TGG CTG GTT GGT GCG CTG CAG ACC TTC GGT TCT CTG GCG GAC GTT GCG GGT ACC

           V   S   N   I   V   Y   Q   Q   R   Q   A   A   Q   L   E   K   Q   N   E   L
     Query GTT TCT AAC ATA GTG TAC CAG CAG CGC CAG GCC GCT CAA TTG GAG AAA CAA AAT GAA CTA
           ||| ||| ||| ||* ||* ||| ||| ||| ||# ||| ||* ||* ||# #|| ||# ||| ||# ||# ||| ||#
 Optimized GTT TCT AAC ATC GTT TAC CAG CAG CGT CAG GCG GCG CAG CTG GAA AAA CAG AAC GAA CTG

           M   E   T   W   M   N   K   Q   E   A   L   Q   K   S   Q   M   E   L   T   R
     Query ATG GAG ACA TGG ATG AAC AAA CAA GAG GCA TTG CAA AAA TCC CAA ATG GAA TTG ACC CGA
           ||| ||# ||* ||| ||| ||| ||| ||# ||# ||# #|| ||# ||| ||# ||# ||| ||| #|| ||| ||*
 Optimized ATG GAA ACC TGG ATG AAC AAA CAG GAA GCG CTG CAG AAA TCT CAG ATG GAA CTG ACC CGT

           D   L   S   I   N   G   P   A   A   R   V   Q   S   A   L   D   A   G   F   D
     Query GAC CTC TCC ATC AAT GGT CCA GCT GCC CGG GTG CAA TCA GCA CTC GAC GCA GGT TTC GAT
           ||| ||* ||# ||| ||# ||| ||# ||* ||* ||* ||* ||# ||* ||# ||* ||| ||# ||| ||| ||#
 Optimized GAC CTG TCT ATC AAC GGT CCG GCG GCG CGT GTT CAG TCT GCG CTG GAC GCG GGT TTC GAC

           A   V   S   A   R   R   I   A   G   S   G   E   R   V   I   W   G   N   L   D
     Query GCA GTG AGT GCT CGG CGC ATT GCT GGT TCA GGT GAG CGC GTG ATA TGG GGG AAC TTG GAT
           ||# ||* **| ||* ||* ||# ||# ||* ||| ||* ||| ||# ||# ||* ||* ||| ||* ||| #|| ||#
 Optimized GCG GTT TCT GCG CGT CGT ATC GCG GGT TCT GGT GAA CGT GTT ATC TGG GGT AAC CTG GAC

           R   P   I   M   H   A   G   T   M   D   S   I   R   Q   T   K   H   L   D   S
     Query CGC CCC ATC ATG CAC GCT GGC ACC ATG GAC AGC ATA CGC CAA ACC AAA CAT TTG GAC TCT
           ||# ||* ||| ||| ||| ||* ||# ||| ||| ||| **# ||* ||# ||# ||| ||| ||# #|| ||| |||
 Optimized CGT CCG ATC ATG CAC GCG GGT ACC ATG GAC TCT ATC CGT CAG ACC AAA CAC CTG GAC TCT

           L   S   H   S   L   A   T   F   K   N   G   T   P   F   G   K   P   A   P   P
     Query CTG AGT CAC TCA TTG GCC ACA TTT AAA AAT GGA ACC CCA TTC GGA AAA CCT GCG CCC CCA
           ||| **| ||| ||* #|| ||* ||* ||# ||| ||# ||* ||| ||# ||| ||* ||| ||* ||| ||* ||#
 Optimized CTG TCT CAC TCT CTG GCG ACC TTC AAA AAC GGT ACC CCG TTC GGT AAA CCG GCG CCG CCG

           T   A   K   F   G   K   P   Q   A   T   T   A   Q   I   T   I   G   H   N   P
     Query ACA GCA AAA TTT GGT AAG CCC CAA GCC ACA ACC GCA CAA ATC ACT ATT GGC CAT AAC CCT
           ||* ||# ||| ||# ||| ||# ||* ||# ||* ||* ||| ||# ||# ||| ||# ||# ||# ||# ||| ||*
 Optimized ACC GCG AAA TTC GGT AAA CCG CAG GCG ACC ACC GCG CAG ATC ACC ATC GGT CAC AAC CCG

           G   S   S   S   V   .
     Query GGT AGC TCC AGT GTC TAA
           ||| **# ||# **| ||# |||
 Optimized GGT TCT TCT TCT GTT TAA
```

**Chiba/000496 (AJ606693.2)** – 498 nucleotides; 378 unchanged, 68 transition and 52 transversion changes.

```
            M   S   W   L   V   G   A   L   Q   T   A   G   S   L   V   D   T   A   G   T
    Query   ATG AGT TGG TTA GTT GGT GCA CTT CAA ACG GCA GGC TCC CTC GTG GAC ACT GCT GGT ACA
            ||| **| ||| #|# ||| ||| ||# ||* ||# ||* ||# ||# ||# ||* ||* ||| ||# ||* ||| ||*
Optimized   ATG TCT TGG CTG GTT GGT GCG CTG CAG ACC GCG GGT TCT CTG GTT GAC ACC GCG GGT ACC

            I   S   N   I   V   Y   Q   Q   R   Q   A   A   Q   L   E   K   Q   N   Q   L
    Query   ATT TCC AAC ATT GTG TAT CAA CAG AGA CAA GCA GCT CAA TTG GAA AAA CAG AAT CAG CTG
            ||# ||# ||| ||# ||* ||# ||# ||| *|* ||# ||# ||* ||# #|| ||| ||| ||| ||# ||| |||
Optimized   ATC TCT AAC ATC GTT TAC CAG CAG CGT CAG GCG GCG CAG CTG GAA AAA CAG AAC CAG CTG

            M   E   M   W   M   N   K   Q   E   A   L   Q   K   S   Q   M   E   L   S   R
    Query   ATG GAG ATG TGG ATG AAC AAA CAA GAA GCA CTG CAA AAA TCC CAG ATG GAA CTG TCA CGT
            ||| ||# ||| ||| ||| ||| ||| ||# ||| ||# ||| ||# ||| ||# ||| ||| ||| ||| ||* |||
Optimized   ATG GAA ATG TGG ATG AAC AAA CAG GAA GCG CTG CAG AAA TCT CAG ATG GAA CTG TCT CGT

            D   L   A   V   N   G   P   A   L   R   V   Q   A   A   L   D   A   G   F   D
    Query   GAT TTG GCT GTC AAT GGC CCG GCA CTC CGT GTA CAG GCT GCG TTG GAT GCT GGC TTC GAC
            ||# #|| ||* ||# ||# ||# ||| ||# ||* ||| ||* ||| ||* ||| #|| ||# ||* ||# ||| |||
Optimized   GAC CTG GCG GTT AAC GGT CCG GCG CTG CGT GTT CAG GCG GCG CTG GAC GCG GGT TTC GAC

            A   V   S   A   R   R   M   A   G   S   G   E   R   V   I   W   G   N   L   D
    Query   GCT GTT AGT GCT CGA AGA ATG GCA GGT TCC GGT GAG AGG GTT ATA TGG GGC AAT TTA GAC
            ||* ||| **| ||* ||* *|* ||| ||# ||| ||# ||| ||# *|* ||| ||* ||| ||# ||# #|# |||
Optimized   GCG GTT TCT GCG CGT CGT ATG GCG GGT TCT GGT GAA CGT GTT ATC TGG GGT AAC CTG GAC

            R   P   L   M   H   A   G   T   M   D   S   I   R   N   T   K   H   L   E   S
    Query   CGC CCA CTT ATG CAT GCT GGC ACC ATG GAC AGC ATC CGT AAC ACC AAA CAC CTT GAG TCC
            ||# ||# ||* ||| ||# ||* ||# ||| ||| ||| **# ||| ||| ||| ||| ||| ||| ||* ||# ||#
Optimized   CGT CCG CTG ATG CAC GCG GGT ACC ATG GAC TCT ATC CGT AAC ACC AAA CAC CTG GAA TCT

            I   S   H   S   L   A   T   F   K   N   G   T   P   F   G   K   S   A   P   P
    Query   ATC AGC CAC TCC CTT GCC ACG TTC AAA AAT GGC ACC CCA TTT GGC AAA TCA GCA CCA CCT
            ||| **# ||| ||# ||* ||* ||* ||| ||| ||# ||# ||| ||# ||# ||# ||| ||* ||# ||# ||*
Optimized   ATC TCT CAC TCT CTG GCG ACC TTC AAA AAC GGT ACC CCG TTC GGT AAA TCT GCG CCG CCG

            Q   F   K   Q   G   R   P   A   A   T   T   A   Q   I   Q   L   G   P   K   P
    Query   CAA TTC AAA CAG GGC AGG CCA GCT GCC ACA ACT GCA CAA ATC CAA TTA GGA CCT AAA CCT
            ||# ||| ||| ||| ||# *|* ||# ||* ||* ||* ||# ||# ||# ||| ||# #|# ||* ||* ||| ||*
Optimized   CAG TTC AAA CAG GGT CGT CCG GCG GCG ACC ACC GCG CAG ATC CAG CTG GGT CCG AAA CCG

            G   S   S   S   V   .
    Query   GGT AGC TCC AGC GTC TAA
            ||| **# ||# **# ||# |||
Optimized   GGT TCT TCT TCT GTT TAA
```

**Kashiwa1 (LC190463.1)** – 501 nucleotides; 376 unchanged, 66 transition and 59 transversion changes.

```
           M   S   W   F   N   A   A   A   L   G   L   G   S   V   V   D   M   A   G   T
    Query  ATG TCT TGG TTC AAT GCT GCA GCA CTT GGT CTT GGC TCT GTC GTG GAC ATG GCC GGG ACG
           ||| ||| ||| ||| ||# ||* ||# ||# ||* ||| ||* ||# ||| ||# ||* ||| ||| ||* ||* ||*
Optimized  ATG TCT TGG TTC AAC GCG GCG GCG CTG GGT CTG GGT TCT GTT GTT GAC ATG GCG GGT ACC

           I   S   S   I   V   A   Q   H   R   Q   I   D   M   M   A   E   A   N   R   I
    Query  ATT TCC TCC ATT GTT GCC CAA CAT CGG CAA ATA GAC ATG ATG GCT GAA GCT AAC AGA ATA
           ||# ||# ||# ||# ||| ||* ||# ||# ||* ||# ||* ||| ||| ||| ||* ||| ||* ||| *|* ||*
Optimized  ATC TCT TCT ATC GTT GCG CAG CAC CGT CAG ATC GAC ATG ATG GCG GAA GCG AAC CGT ATC

           Q   A   D   W   V   R   R   Q   E   Q   L   L   M   R   G   Q   D   M   T   R
    Query  CAG GCA GAT TGG GTT CGC CGC CAA GAA CAG TTG CTC ATG CGT GGG CAG GAT ATG ACA CGT
           ||| ||# ||# ||| ||| ||# ||# ||# ||| ||| #|| ||* ||| ||| ||* ||| ||# ||| ||* |||
Optimized  CAG GCG GAC TGG GTT CGT CGT CAG GAA CAG CTG CTG ATG CGT GGT CAG GAC ATG ACC CGT

           D   L   S   I   N   G   P   A   Q   R   V   E   S   L   V   N   A   G   F   T
    Query  GAT TTG TCA ATC AAT GGC CCC GCA CAG CGT GTG GAG TCC CTT GTC AAC GCA GGG TTC ACT
           ||# #|| ||* ||| ||# ||# ||* ||# ||| ||| ||* ||# ||# ||* ||# ||| ||# ||* ||| ||#
Optimized  GAC CTG TCT ATC AAC GGT CCG GCG CAG CGT GTT GAA TCT CTG GTT AAC GCG GGT TTC ACC

           P   T   D   A   R   R   M   V   G   G   S   E   T   V   S   Y   G   L   L   D
    Query  CCC ACG GAC GCC CGC CGC ATG GTG GGT GGC TCC GAA ACA GTC AGT TAT GGG TTG CTG GAT
           ||* ||* ||| ||* ||# ||# ||| ||* ||| ||# ||# ||| ||* ||# **| ||# ||* #|| ||| ||#
Optimized  CCG ACC GAC GCG CGT CGT ATG GTT GGT GGT TCT GAA ACC GTT TCT TAC GGT CTG CTG GAC

           R   P   V   L   Q   R   S   I   L   A   G   I   S   E   T   R   H   L   Q   N
    Query  CGA CCC GTG CTG CAA AGG AGC ATC TTG GCA GGC ATC AGT GAA ACT CGC CAT CTC CAA AAC
           ||* ||* ||* ||| ||# *|* **# ||| #|| ||# ||# ||| **| ||| ||# ||# ||# ||* ||# |||
Optimized  CGT CCG GTT CTG CAG CGT TCT ATC CTG GCG GGT ATC TCT GAA ACC CGT CAC CTG CAG AAC

           M   Q   G   T   L   S   A   F   K   T   G   T   G   V   G   T   K   P   A   P
    Query  ATG CAG GGG ACA CTC AGT GCC TTC AAG ACT GGC ACG GGT GTC GGC ACT AAG CCA GCA CCT
           ||| ||| ||* ||* ||* **| ||* ||| ||# ||# ||# ||* ||| ||# ||# ||# ||# ||# ||# ||*
Optimized  ATG CAG GGT ACC CTG TCT GCG TTC AAA ACC GGT ACC GGT GTT GGT ACC AAA CCG GCG CCG

           A   G   F   G   N   P   N   Y   K   T   G   P   P   K   I   N   L   G   F   N
    Query  GCC GGA TTT GGT AAT CCA AAT TAC AAA ACT GGC CCC CCT AAA ATC AAC CTT GGA TTC AAT
           ||* ||* ||# ||| ||# ||# ||# ||| ||| ||# ||# ||* ||* ||| ||| ||| ||* ||* ||| ||#
Optimized  GCG GGT TTC GGT AAC CCG AAC TAC AAA ACC GGT CCG CCG AAA ATC AAC CTG GGT TTC AAC

           P   P   S   T   N   V   .
    Query  CCC CCT AGC ACT AAT GTC TAA
           ||* ||* **# ||# ||# ||# |||
Optimized  CCG CCG TCT ACC AAC GTT TAA
```

**Mc10 (AY237420.2)** – 501 nucleotides; 378 unchanged, 65 transition and 58 transversion changes.

```
           M   S   W   F   T   G   A   S   L   A   A   G   S   L   V   D   M   A   G   T
    Query  ATG TCT TGG TTT ACT GGA GCA TCT CTG GCT GCC GGT TCA CTC GTG GAC ATG GCA GGC ACC
           ||| ||| ||| ||# ||# ||* ||# ||| ||| ||* ||* ||| ||* ||* ||* ||| ||| ||# ||# |||
Optimized  ATG TCT TGG TTC ACC GGT GCG TCT CTG GCG GCG GGT TCT CTG GTT GAC ATG GCG GGT ACC

           I   S   S   I   V   A   Q   Q   R   Q   I   D   L   M   A   E   A   N   R   I
    Query  ATT TCA TCA ATT GTG GCA CAA CAA AGA CAA ATT GAT CTG ATG GCA GAA GCA AAT AGA ATC
           ||# ||* ||* ||# ||* ||# ||# ||# *|* ||# ||# ||# ||| ||| ||# ||| ||# ||# *|* |||
Optimized  ATC TCT TCT ATC GTT GCG CAG CAG CGT CAG ATC GAC CTG ATG GCG GAA GCG AAC CGT ATC

           Q   A   D   W   V   R   R   Q   E   A   L   Q   I   R   G   Q   D   I   S   R
    Query  CAG GCA GAT TGG GTG CGC CGT CAA GAG GCA CTA CAA ATC CGT GGC CAG GAC ATC TCA CGG
           ||| ||# ||# ||| ||* ||# ||| ||# ||# ||# ||# ||# ||| ||| ||# ||| ||| ||| ||* ||*
Optimized  CAG GCG GAC TGG GTT CGT CGT CAG GAA GCG CTG CAG ATC CGT GGT CAG GAC ATC TCT CGT

           D   L   A   V   N   G   T   A   Q   R   V   E   S   L   V   N   A   G   F   T
    Query  GAT CTT GCT GTT AAC GGC ACT GCC CAG CGT GTT GAG TCT TTA GTC AAT GCA GGC TTC ACA
           ||# ||* ||* ||| ||| ||# ||# ||* ||| ||| ||| ||# ||| #|# ||# ||# ||# ||# ||| ||*
Optimized  GAC CTG GCG GTT AAC GGT ACC GCG CAG CGT GTT GAA TCT CTG GTT AAC GCG GGT TTC ACC

           P   V   D   A   R   R   L   A   G   G   T   E   T   V   S   Y   G   L   L   D
    Query  CCC GTG GAC GCA CGT CGG CTG GCC GGC GGA ACG GAA ACG GTG AGT TAC GGC CTA CTG GAT
           ||* ||* ||| ||# ||| ||* ||| ||* ||# ||* ||* ||| ||* ||* **| ||| ||# ||# ||| ||#
Optimized  CCG GTT GAC GCG CGT CGT CTG GCG GGT GGT ACC GAA ACC GTT TCT TAC GGT CTG CTG GAC

           R   P   I   L   Q   R   G   I   L   S   G   I   T   E   T   R   H   L   Q   A
    Query  CGC CCT ATC CTA CAA CGG GGC ATC CTT TCT GGC ATC ACT GAG ACA CGA CAC CTC CAG GCC
           ||# ||* ||| ||# ||# ||* ||# ||| ||* ||| ||# ||| ||# ||# ||* ||* ||| ||* ||| ||*
Optimized  CGT CCG ATC CTG CAG CGT GGT ATC CTG TCT GGT ATC ACC GAA ACC CGT CAC CTG CAG GCG

           M   Q   G   A   L   S   A   F   K   N   G   A   S   Y   G   A   P   P   A   P
    Query  ATG CAG GGC GCT CTA AGT GCA TTC AAA AAT GGT GCC TCT TAC GGA GCC CCG CCA GCC CCA
           ||| ||| ||# ||* ||# **| ||# ||| ||| ||# ||| ||* ||| ||| ||* ||* ||| ||# ||* ||#
Optimized  ATG CAG GGT GCG CTG TCT GCG TTC AAA AAC GGT GCG TCT TAC GGT GCG CCG CCG GCG CCG

           S   G   F   V   N   P   N   Y   Q   P   S   P   P   R   L   K   L   G   P   R
    Query  TCA GGC TTT GTG AAT CCA AAT TAT CAA CCT TCA CCT CCG AGA TTG AAA CTA GGC CCT AGG
           ||* ||# ||# ||* ||# ||# ||# ||# ||# ||* ||* ||* ||| *|* #|| ||| ||# ||# ||* *|*
Optimized  TCT GGT TTC GTT AAC CCG AAC TAC CAG CCG TCT CCG CCG CGT CTG AAA CTG GGT CCG CGT

           P   P   S   T   N   V   .
    Query  CCC CCT AGC ACC AAT GTT TGA
           ||* ||* **# ||| ||# ||| |||
Optimized  CCG CCG TCT ACC AAC GTT TGA
```

**C12 (AY603425.1)** – 501 nucleotides; 378 unchanged, 62 transition and 61 transversion changes.

```
           M   S   W   F   T   G   A   A   L   A   A   G   S   L   V   D   M   A   G   T
Query      ATG TCT TGG TTC ACA GGA GCA GCT CTT GCA GCC GGG TCC CTC GTG GAC ATG GCT GGT ACT
           ||| ||| ||| ||| ||* ||* ||# ||* ||* ||# ||* ||* ||# ||* ||* ||| ||| ||* ||| ||#
Optimized  ATG TCT TGG TTC ACC GGT GCG GCG CTG GCG GCG GGT TCT CTG GTT GAC ATG GCG GGT ACC

           V   S   S   I   V   A   Q   H   R   Q   I   D   L   M   A   Q   A   N   Q   I
Query      GTA TCA TCC ATT GTG GCA CAA CAC CGC CAA ATT GAC TTG ATG GCA CAA GCA AAC CAG ATT
           ||* ||* ||# ||# ||* ||# ||# ||| ||# ||# ||# ||| #|| ||| ||# ||# ||# ||| ||| ||#
Optimized  GTT TCT TCT ATC GTT GCG CAG CAC CGT CAG ATC GAC CTG ATG GCG CAG GCG AAC CAG ATC

           Q   R   D   W   V   N   K   Q   E   A   L   M   R   R   G   Q   D   I   S   R
Query      CAA AGA GAC TGG GTC AAC AAG CAA GAG GCA CTC ATG CGG CGA GGT CAG GAC ATC TCA CGC
           ||# *|* ||| ||| ||# ||| ||# ||# ||# ||# ||* ||| ||* ||* ||| ||| ||| ||| ||* ||#
Optimized  CAG CGT GAC TGG GTT AAC AAA CAG GAA GCG CTG ATG CGT CGT GGT CAG GAC ATC TCT CGT

           D   L   A   V   N   G   T   A   Q   R   V   E   S   L   V   G   A   G   F   T
Query      GAC CTT GCG GTC AAC GGC ACT GCC CAA CGG GTG GAG TCG CTG GTA GGT GCA GGT TTC ACC
           ||| ||* ||| ||# ||| ||# ||# ||* ||# ||* ||* ||# ||* ||| ||* ||| ||# ||| ||| |||
Optimized  GAC CTG GCG GTT AAC GGT ACC GCG CAG CGT GTT GAA TCT CTG GTT GGT GCG GGT TTC ACC

           P   V   D   A   R   R   L   A   G   S   T   E   T   V   Q   Y   G   L   L   D
Query      CCT GTT GAT GCT CGA CGG CTG GCC GGG AGC ACA GAA ACA GTC CAG TAT GGC CTG CTG GAT
           ||* ||| ||# ||* ||* ||* ||| ||* ||* **# ||* ||| ||* ||# ||| ||# ||# ||| ||| ||#
Optimized  CCG GTT GAC GCG CGT CGT CTG GCG GGT TCT ACC GAA ACC GTT CAG TAC GGT CTG CTG GAC

           R   P   I   L   Q   R   E   V   L   A   G   I   S   E   T   H   H   L   Q   S
Query      CGC CCA ATT CTA CAA CGG GAA GTC CTA GCG GGC ATC AGT GAA ACA CAT CAC CTA CAA TCT
           ||# ||# ||# ||# ||# ||* ||| ||# ||# ||| ||# ||| **| ||| ||* ||# ||| ||# ||# |||
Optimized  CGT CCG ATC CTG CAG CGT GAA GTT CTG GCG GGT ATC TCT GAA ACC CAC CAC CTG CAG TCT

           M   Q   A   S   L   S   A   F   K   K   G   S   S   Y   G   A   P   P   A   P
Query      ATG CAA GCT TCT TTG AGT GCA TTC AAG AAA GGG TCC AGT TAT GGT GCC CCA CCT GCC CCT
           ||| ||# ||* ||| #|| **| ||# ||| ||# ||| ||* ||# **| ||# ||| ||* ||# ||* ||* ||*
Optimized  ATG CAG GCG TCT CTG TCT GCG TTC AAA AAA GGT TCT TCT TAC GGT GCG CCG CCG GCG CCG

           V   G   F   A   N   P   N   F   Q   P   A   P   P   R   V   N   L   G   F   R
Query      GTT GGG TTT GCT AAC CCA AAC TTC CAA CCG GCC CCT CCT AGG GTG AAT TTA GGG TTT AGG
           ||| ||* ||# ||* ||| ||# ||| ||| ||# ||| ||* ||* ||* *|* ||* ||# #|# ||* ||# *|*
Optimized  GTT GGT TTC GCG AAC CCG AAC TTC CAG CCG GCG CCG CCG CGT GTT AAC CTG GGT TTC CGT

           P   P   S   T   N   V   .
Query      CCT CCA TCA ACC AAT GTT TAG
           ||* ||# ||* ||| ||# ||| |||
Optimized  CCG CCG TCT ACC AAC GTT TAG
```

**NGY-1 (AB775659.2)** – 504 nucleotides; 375 unchanged, 78 transition and 51 transversion changes.

```
           M   S   W   L   V   G   T   L   Q   S   L   G   G   L   T   D   V   A   S   T
    Query  ATG AGC TGG TTG GTA GGC ACT TTG CAA TCT CTT GGT GGG CTC ACT GAC GTT GCG TCC ACC
           ||| **# ||| #|| ||* ||# ||# #|| ||# ||| ||* ||| ||* ||* ||# ||| ||| ||| ||# |||
Optimized  ATG TCT TGG CTG GTT GGT ACC CTG CAG TCT CTG GGT GGT CTG ACC GAC GTT GCG TCT ACC

           I   S   G   M   V   Y   Q   H   R   H   L   D   Q   L   K   R   Q   N   D   L
    Query  ATT TCT GGC ATG GTT TAT CAA CAT AGA CAT CTT GAC CAA CTG AAA AGA CAG AAT GAT CTA
           ||# ||| ||# ||| ||| ||# ||# ||# *|* ||# ||* ||| ||# ||| ||| *|* ||| ||# ||# ||#
Optimized  ATC TCT GGT ATG GTT TAC CAG CAC CGT CAC CTG GAC CAG CTG AAA CGT CAG AAC GAC CTG

           Q   E   Q   W   M   A   R   N   E   Q   L   Q   R   D   A   M   Q   L   T   Q
    Query  CAA GAA CAG TGG ATG GCC CGC AAT GAA CAA CTG CAG CGA GAT GCA ATG CAA TTA ACT CAG
           ||# ||| ||| ||| ||| ||* ||# ||# ||| ||# ||| ||| ||* ||# ||# ||| ||# #|# ||# |||
Optimized  CAG GAA CAG TGG ATG GCG CGT AAC GAA CAG CTG CAG CGT GAC GCG ATG CAG CTG ACC CAG

           D   L   A   V   N   A   P   A   M   R   V   Q   A   A   L   N   A   G   F   D
    Query  GAT TTG GCA GTC AAT GCC CCA GCG ATG AGA GTG CAG GCC GCT CTC AAT GCT GGG TTT GAT
           ||# #|| ||# ||# ||# ||* ||# ||| ||| *|* ||* ||| ||* ||* ||* ||# ||* ||* ||# ||#
Optimized  GAC CTG GCG GTT AAC GCG CCG GCG ATG CGT GTT CAG GCG GCG CTG AAC GCG GGT TTC GAC

           V   V   S   A   R   Q   L   A   G   S   T   E   R   R   I   N   G   Y   L   D
    Query  GTG GTT AGT GCG CGC CAG CTT GCC GGT TCA ACT GAG CGA AGG ATC AAC GGC TAT TTG GAT
           ||* ||| **| ||| ||# ||| ||* ||* ||| ||* ||# ||# ||* *|* ||| ||| ||# ||# #|| ||#
Optimized  GTT GTT TCT GCG CGT CAG CTG GCG GGT TCT ACC GAA CGT CGT ATC AAC GGT TAC CTG GAC

           Q   P   I   R   T   I   D   Q   A   M   A   V   Q   S   R   G   N   L   T   S
    Query  CAG CCA ATT CGC ACT ATT GAT CAG GCA ATG GCA GTG CAG TCT AGG GGC AAT TTG ACT TCG
           ||| ||# ||# ||# ||# ||# ||# ||| ||# ||| ||# ||* ||| ||| *|* ||# ||# #|| ||# ||*
Optimized  CAG CCG ATC CGT ACC ATC GAC CAG GCG ATG GCG GTT CAG TCT CGT GGT AAC CTG ACC TCT

           L   S   N   A   L   A   T   Y   Q   K   G   T   Q   F   G   L   K   Q   P   K
    Query  TTG TCG AAC GCC CTT GCC ACT TAC CAA AAA GGC ACT CAA TTT GGA CTC AAA CAA CCC AAA
           #|| ||* ||| ||* ||* ||* ||# ||| ||# ||| ||# ||# ||# ||# ||* ||* ||| ||# ||* |||
Optimized  CTG TCT AAC GCG CTG GCG ACC TAC CAG AAA GGT ACC CAG TTC GGT CTG AAA CAG CCG AAA

           G   F   K   S   P   I   A   A   E   Q   S   R   G   P   T   I   T   L   G   P
    Query  GGA TTT AAG AGC CCA ATT GCT GCT GAA CAA TCT CGT GGT CCA ACC ATT ACA CTT GGC CCT
           ||* ||# ||# **# ||# ||# ||* ||* ||| ||# ||| ||| ||| ||# ||| ||# ||* ||* ||# ||*
Optimized  GGT TTC AAA TCT CCG ATC GCG GCG GAA CAG TCT CGT GGT CCG ACC ATC ACC CTG GGT CCG

           P   P   P   S   T   N   L   .
    Query  CCC CCA CCA TCT ACT AAT CTA TAA
           ||* ||# ||# ||| ||# ||# ||# |||
Optimized  CCG CCG CCG TCT ACC AAC CTG TAA
```
